# Mendelian randomization for multiple exposures and outcomes with Bayesian Directed Acyclic Graphs exploration and causal effects estimation

**DOI:** 10.1101/2024.06.18.599498

**Authors:** Verena Zuber, Toinét Cronjé, Na Cai, Dipender Gill, Leonardo Bottolo

## Abstract

Current Mendelian randomization (MR) methods do not reflect complex relationships among multiple exposures and outcomes as is typical for real-life applications. We introduce MrDAG the first MR method to model dependency relations within the exposures, the outcomes, and between them to improve causal effects estimation. MrDAG combines three causal inference strategies in a unified manner. It uses genetic variation as instrumental variables to account for unmeasured confounders. It performs structure learning to detect and orientate the direction of the dependencies within exposures and outcomes. Finally, interventional calculus is employed to derive principled causal effect estimates. MrDAG was motivated to unravel how lifestyle and behavioural exposures impact mental health. It highlights education and smoking as key effective points of intervention given their down-stream effects on mental health. These insights would have been difficult to delineate without modelling the causal paths between multiple exposures and outcomes at once.

## Introduction

Genetic evidence is increasingly used to infer causal relationships between human traits in Mendelian randomization (MR) analysis. The standard MR paradigm, one exposure and one outcome, can be biased by unobserved pleiotropy. It occurs when the genetic variants used as instruments in the MR analysis act *via* separate pathways to the exposure under investigation. Extensions to consider multiple exposures [1] along with multi-response [2] of standard MR allow to model pleiotropy acting either *via* any of the exposures or any of the outcomes or both, respectively.

Yet, to date, there is no MR approach which can estimate the dependency relations within the exposures and the outcomes to enhance the detection of causal effects between them and improve their accuracy. As we show in our motivating data application on mental health phenotypes, it is a common problem in practical applications that the effect of an exposure on an outcome can be confounded or (partially or completely) mediated by another exposure [3] or mediated by another outcome, or both. However, this structure is latent and not known and consequently needs to be learned from the data. This problem has been overlooked in the literature and current MR implementations which do not account for these dependencies likely produce spurious findings which are often claimed as supporting causality in applied analysis.

Here, we address this gap by proposing the MrDAG model, the first Mendelian randomization method with Directed Acyclic Graphs (DAGs) exploration and causal effects estimation, which utilises summary-level genetic associations from genome-wide association studies to learn how interrelated exposures affect multiple outcomes which, in turn, are interconnected in a complex fashion. MrDAG is a Bayesian causal graphical model that combines three causal inference strategies in a unified manner. First, the MR paradigm which uses genetic variation as instrumental variables (IVs) [4, 5] to ensure unconfoundedness. Second, structure learning [6], *i*.*e*., graphical models selection to define the graphs that best describe the dependency structure in a given data set under the constraint on edges’ orientation from the exposures to the outcomes implied by the MR paradigm. Third, interventional calculus to derive principled causal effects estimates [7] from the exposures to the outcomes.

Our motivating real data application considers the impact of six common modifiable lifestyle and behavioural exposures on seven mental health phenotypes. Mental health describes patterns of cognitive, emotional, and behavioural disregulations that limit daily functioning and cause distress. One in eight individuals suffers from one or more mental health phenotypes worldwide, most commonly anxiety, attention-deficit hyperactivity, autism spectrum, bipolar, eating, personality or schizophrenia-related diseases [8]. Collectively, they contribute to more than 15% of total years lived with disability [9]. Clinically, mental health phenotypes are notoriously difficult to disentangle and diagnose due to the lack of objective biological biomarkers and distinct disease impressions [10]. No symptom can be uniquely ascribed to one disease, and each disease comprises experiencing a group of interrelated traits. In research, this complexity is reinforced by the multifaceted mechanisms that cause and sustain mental health [10, 11]. In addition to genetic liability, numerous behavioural and lifestyle factors such as alcohol consumption, smoking, sleep hygiene, physical activity and education contribute to the risk of developing a mental health trait [11, 12]. Notably, these factors are also affected by existing disease and treatment [13]. It is essential to appreciate these complexities when attempting to identify distinct and shared underlying mechanisms of mental health. While MR studies have been effective in circumventing some of the limitations of traditional epidemiology such as environmental confounding and reverse causation, MR remains largely unable to fully disentangle the interplay between traits that cause or result from mental health [14]. The complexity of such an example demonstrates the limitations of current MR solutions to offer a more comprehensive picture of causal mechanisms between complex phenotypes and provides a suitable test ground for the application of the proposed methodology.

## Results

### Causal inferential strategies in MrDAG

MrDAG combines three causal inference strategies.

First, MR has pioneered the ability to use genetic data as IVs to derive causal statements from observational data despite the presence of unmeasured confounders [15, 16].

Second, in its standard formulation of one exposure and one outcome, the conditional dependencies between the outcome *Y*, the exposure *X*, the IV *G* and the unmeasured confounder *U* are all given as well as their graphical representation [5]. When multiple exposures ***X*** [1] and multiple outcomes ***Y*** [2] are considered along with multiple IVs ***G***, (partial) correlation between ***X*** and conditional dependencies between ***Y*** are included in the models to perform the selection of important exposures whose causal effects can be shared or are distinct across the responses. However, to date, no dependency relations within the exposures and within the outcomes are estimated, although, in practical applications, the effect of an exposure on an outcome can be confounded or (partially or completely) mediated by another exposure or mediated by another outcome, or both, see Figures 1A-B for an illustration. Moreover, dependency relations are also important to derive principled estimates of the causal effects [7].

**Figure 1.**
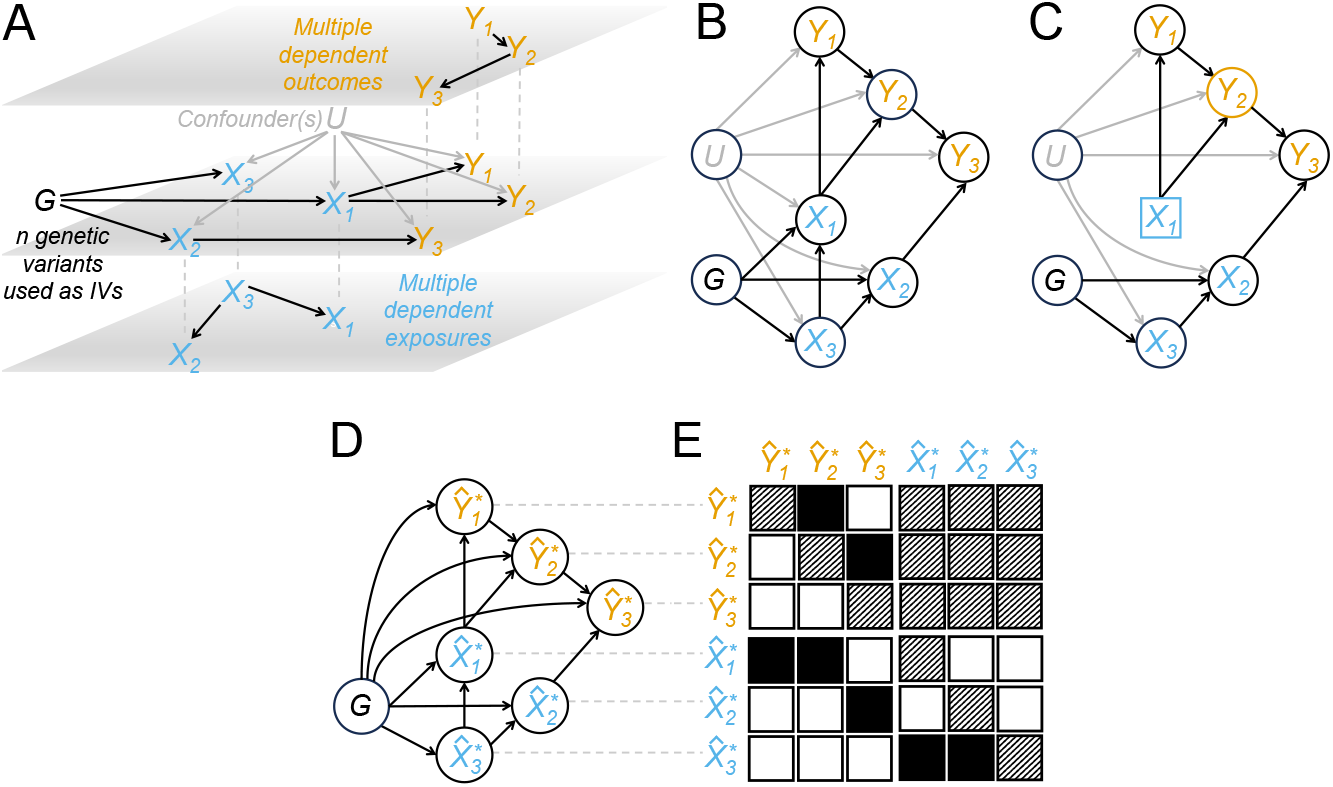
Directed Acyclic Graph (DAG) representation of the proposed multiple exposures and multiple outcomes Mendelian randomization model and causal effects estimation. (**A**) Middle panel: Multivariable Mendelian randomization for multiple responses with ***G*** = (*G*_1_, …, *G*_*n*_)^⊤^: Genetic variants (black) or instrumental variables (IVs); ***X*** = (*X*_1_, *X*_2_, *X*_3_)^⊤^: Exposures (blue); ***Y*** = (*Y*_1_, *Y*_2_, *Y*_3_)^⊤^: Responses (orange); *U* : Unmeasured confounder(s) (grey). True (unconfounded by *U*) exposure-outcome dependency relations depicted in the middle panel are classified as follows: *X*_1_ has *shared* causal effect on responses *Y*_1_ and *Y*_2_, while *X*_2_ has a *distinct* causal effect on response *Y*_3_. *X*_3_ does not have any effect on the outcomes. Bottom panel: True fork structure within the exposures with *X*_3_ regarded as the common cause of *X*_1_ and *X*_2_. Top panel: True chain structure within the outcomes, where *Y*_1_ affects *Y*_3_ through *Y*_2_. (**B**) DAG is obtained by combining the true exposure-outcome dependency relations ((A) middle panel), the fork structure within the exposures ((A) bottom panel) and the chain structure within the responses ((A) top panel). When looking at the effect of *X*_1_ on *Y*_3_, *X*_3_ (along with ***G*** and *U*) is a confounder of *X*_1_ and *Y*_2_ is a complete mediator. Without conditioning on *Y*_2_, with the same set of confounders, a spurious association would be found between *X*_1_ and *Y*_3_. (**C**) Estimation of the causal effect under intervention in *X*_1_ on *Y*_2_, highlighted in blue and orange, respectively. The representation of *X*_1_ has changed from a circle to a square to emphasise that, under intervention, it is no longer a random variable and it is now set at 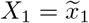. Intervention affects only the conditional distribution of *X*_1_, *i*.*e*., *X*_1_ | (*X*_3_, ***G***, *U*) and it leaves unaltered all the others. From a practical perspective, it would be sufficient to condition on *X*_3_, ***G*** and *U* (graphically, the directed edges to *X*_1_ from *X*_3_, ***G*** and *U* are removed) to guarantee that the association between *X*_1_ and *Y*_2_ is purely causative (see Supplementary Figure 1). However, since *U* is unobserved, the estimation of the causal effects cannot be obtained only by conditioning. (**D**) Genetically predicted exposures 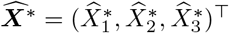 and outcomes 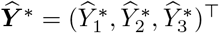 depend only on ***G*** which are chosen to be associated only with ***X*** and not with ***Y***.Graphically, no directed edges to 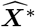 and 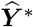 from *U* are pictured. True (unconfounded by *U*) dependency relations between the traits in the original (individual-level) data shown in (B) are obtained by the DAG estimated by using 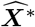 and 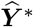. (**E**) Adjacency matrix describing the Markov properties of the DAG obtained by using genetically predicted exposures and outcomes (the variables in the *x*-axis are dependent on the variables in the *y*-axis) which are function of the IVs and the inverse-variance weighted (IVW) (depicted with a “*”) summary-level statistics 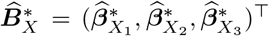 and 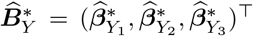. Neither reverse causation (top-right submatrix) nor feedback loops (main diagonal) are allowed. Colour code: Black, directed edge between variables; white, no causal relationship between variables; black-white strips, directed edge not allowed (feedback loop and reverse causation between exposures and outcomes).

In real data applications, complex dependency relations between the traits are generally not known in advance, and they need to be learned from the data. To detect them, we rely on Directed Acyclic Graphs (DAGs) and structure learning. Graphical models are multivariate distributions associated with a graph and are very effective for encoding conditional dependencies [17] between random variables. They are represented in a graph as nodes (vertices) while edges denote conditional dependence relationships between the corresponding random variables. A DAG is a directed graph, where each edge has an orientation with no directed cycles. Structure learning is a model selection problem [6] to estimate the DAG (or competing DAGs) that best describes the dependency structure in a given data set. However, without identifiability conditions [18], it is not possible to estimate uniquely the underlying DAG since its conditional independencies can be associated with several alternative DAGs. The set of DAGs that hold the same conditional independencies is known as Markov Equivalent Class and the best that can be done from observational data is to estimate this class (or competing classes). Thus, this paper aims to illustrate how to perform DAG exploration (whose importance will be apparent in the next paragraph) which belongs to the Markov Equivalent Classes that best fit the data under the constraint on the orientation of the edges, known as partial ordering [19], from the exposures to the outcomes implied by the MR paradigm.

Third, besides the identification of the exposure-outcome relations as well as the dependency patterns within the exposures and the outcomes, we are also interested in the causal effects estimation under intervention [7]. An intervention on the exposures can be made explicit by a suitable modification of the multivariate distribution associated with the DAG, under the assumption that the intervention does not affect any other variable in the joint distribution besides the conditional distribution of the exposure under intervention [20]. See Figure 1C for an example of intervention on an exposure and the estimation of the causal effect on an outcome.

In this formulation, all confounders should be measurable to perform structure learning and causal effects estimation (causal sufficiency assumption [21]). This assumption is usually not met in real data applications where, instead, unmeasured confounders are ubiquitous and affect exposures and responses at the same time. To solve this problem, we demonstrate (see Methods) and show in an extensive simulation study (see Results) that, under partial ordering, we can estimate the dependency structure that exists between the traits in the original (individual-level) data unconfounded by *U* by using their genetically predicted values. Since the genetically predicted traits depend only on the selected IVs, the confounders do not mask the true dependency relations required in causal effects estimation. See Figure 1D, where the graphical model estimated by using genetically predicted exposures and outcomes approximates the corresponding DAG in the individual-level data not affected by *U*. Our approach shares some similarities with methods based on the genetic correlation and developed to analyse the joint genetic architecture of complex traits [22] although, in the proposed MR framework, genetic variants are chosen to be valid IVs in contrast to genetic variants chosen for genome-wide [23] or local genetic correlation [24]. Computationally, given the duality between the Markov properties of the DAG and a non-symmetrical adjacency matrix (see Figure 1E), structure learning of the graphical model (or competing graphical models) that best fits the data is performed on a non-symmetrical adjacency matrix which incorporates the constraints on the orientation of the edges from the exposures to the outcomes.

Finally, for a given DAG, we extend results regarding the consistency of the effects of the regressions of the exposures and the outcomes on ***G*** which can be obtained without adjustment on *U* since the genetic variants used as IVs are randomly assigned [4], and show that it is possible to identify and estimate the causal effects between multiple exposures and multiple outcomes based on Pearl’s interventional calculus [7] (see Methods).

The MrDAG model can be summarised as follows:

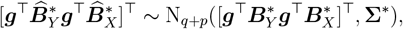

where ***g*** are the observed IVs after pruning or clumping, 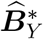 and 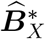 are the inverse-variance weighted (IVW) [25] estimated genetic associations with the outcomes and the exposures, 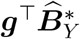 and 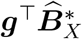 are the genetically predicted values of the outcomes 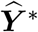 and exposures 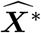 based on the IVs, respectively, which are normally distributed for large sample sizes, and **Σ**^*^ is the genetic covariance matrix that can be partitioned into 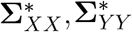 and 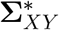, the genetic covariances within the exposures, the outcomes and between them. The MrDAG model allows us to find a solution to the two problems highlighted before. First, we perform DAGs exploration under partial ordering by using 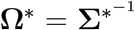, to learn the unconfounded dependency relations within the exposures, the outcomes and between them and to understand the genetic paths that link exposures and outcomes (see Methods). Second, estimate the causal effects of the intervention on the exposures as a function of trait-specific elements of the genetic associations 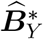 and 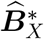 informed by the explored DAGs, unconfounded by any pleiotropic effects within the exposures and the outcomes and any unmeasured confounder.

### Selection of instrumental variables

MrDAG uses the same instrument selection employed in MVMR regardless of the multiple outcomes [2]. A genetic variant is considered a valid instrument for MVMR when three core conditions hold [3]: (IV1) Independence: The variant is independent of all confounders of each of the exposure-outcome associations; (IV2) Relevance: The variant must not be conditional independent of each exposure given the other exposures; (IV3) Exclusion restriction: The variant is independent of the outcome conditional on the exposures and confounders. In practice, only IV2 can be computationally evaluated from the available data. A recent solution to mitigate the effects of weak IVs in MVMR is presented in [26].

There is an important distinction between IV selection in MVMR, as used by MrDAG, and bidirectional MR. Let’s consider two traits A and B. In bidirectional MR, two MR analyses are conducted, one for trait *A* on trait *B*, and then *vice versa*. First, specific IVs are selected for trait A and the first MR model is fit. Then, another set of specific IVs is selected for trait B and the second MR model tests the opposite effects direction. In contrast, in MVMR, IVs are chosen to be the union of genome-wide significant genetic variants for any exposure. By combining MVMR IVs selection approach with DAG learning, MrDAG can infer the bidirectionality of the relationships within exposures based on 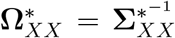 without repeated IVs selection and subsequent analyses. A similar comment can be made for the estimation of the bidirectionality of the relationships within the outcomes based on 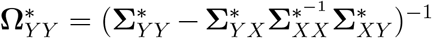 (see Methods). These dependencies should be interpreted as an indication of a violation of condition IV3, *i*.*e*., pleiotropy not explained by the estimated causal effects from the exposures to the outcomes [2]. The detected relationships within the exposures also suggest the existence of measured pleiotropy which, in the proposed framework, comprises confounding, mediation and independent pleiotropic pathways [3].

Overall, only the direction from exposures to outcomes is fixed in MrDAG, and no reverse causation is allowed, reflecting the standard MR paradigm.

### Simulation study

We compare MrDAG in an extensive and comprehensive simulation study where four different *in silico* scenarios have been generated on individual-level data for *N* = 100, 000 individuals. The simulated data sets include *n* = 100 independent genetic variants ***G***, an unmeasured confounder *U*, 15 exposures ***X*** and 5 outcomes ***Y***. All exposures ***X*** were measured on the same individuals in the first sample and have complete overlap as well as all outcomes ***Y*** were measured on the same individuals in the second sample independent of the first sample. In all simulations, the unconfounded dependency relations between the traits are simulated at the individual-level while the algorithms use as input the corresponding IVW summary-level statistics.

The four simulation scenarios are built by combining two different strategies we used to simulate the dependency patterns within the exposures and the responses:

i. “UndG_*X*_-Med_*Y*_ ”. A sparse undirected graphical model (“UndG_*X*_”) encodes the dependency pattern within the exposures ***X*** = (*X*_1_, …, *X*_15_). Regarding the responses ***Y*** = (*Y*_1_, …, *Y*_5_), one outcome is completed mediated by another one (“Med_*Y*_ “). For a visual representation of this scenario, see Figure 2A.
ii. “DAG_*X*_-Med_*Y*_ ”. The dependency relations within the exposures are more complex than in scenario (i) since a topologically ordered DAG within the exposures (“DAG_*X*_”) is simulated [27]. A complete mediation is still considered within the responses. This second scenario is illustrated in Figure 2B.
iii. “UndG_*X*_-DAG_*Y*_ ”. Here, a more complex dependency structure within the individual-level responses (“DAG_*Y*_ ”) is simulated. This scenario is represented in Figure 2C. An example of the complex dependency patterns generated in the simulation study between the traits for one replicate of scenario UndG_*X*_-DAG_*Y*_ is shown in Figure 3A.
iv. “DAG_*X*_-DAG_*Y*_ ”: This is the most complex simulated scenario where two independent topologically ordered DAGs have been simulated within the exposures and outcomes. Figure 2D presents a schematic illustration of this scenario, while Figure 3E shows the intricate dependency structure simulated between the traits for one replicate of DAG_*X*_-DAG_*Y*_ scenario.

**Figure 2.**
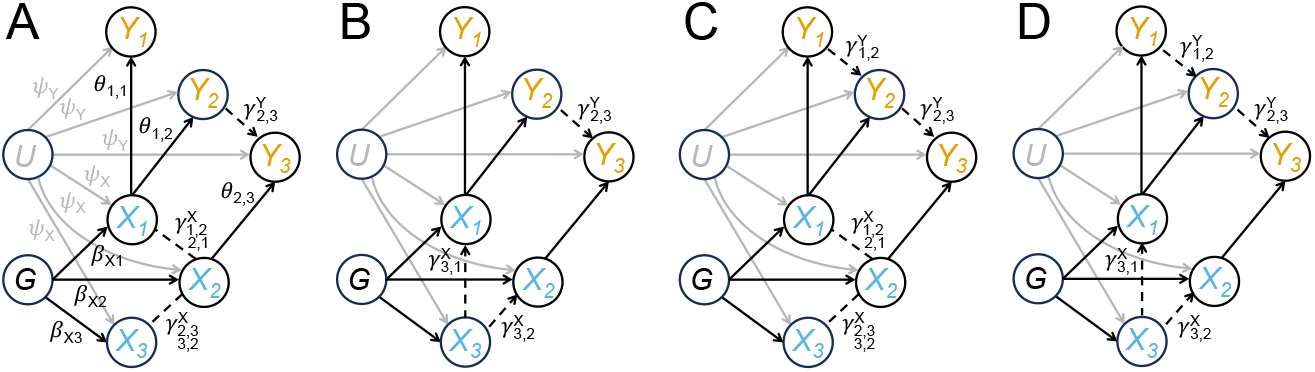
Schematic illustration of different dependency structures simulated between the traits at the individual-level data and the parameters employed in the simulation study. Directed edges indicate dependency relations, while undirected edges denote partial correlations. Dashed lines depict the true (unconfounded by *U*) dependency structure within the exposures and the outcomes, while solid lines indicate true causal effects between them. Parameters *ψ*_*Y*_ and *ψ*_*X*_ indicate the simulated effects of the unmeasured confounder *U* on the exposures and the outcomes, respectively, and 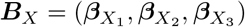 are the simulated genetic effects on the exposures. For simplicity, they are shown only on the left panel. **Θ** = (*θ*_1,1_, *θ*_1,2_, *θ*_2,3_) are the simulated causal effects from the exposures to the outcomes while 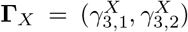 and 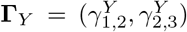 are the mediation parameters within the exposures and the outcomes, respectively, where the subscripts denote their directionality. When partial correlations are simulated within the exposures, bidirectional effects are depicted with double subscripts, *i*.*e*., 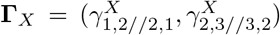. (**A**) Simulated scenario “UndG_*X*_ -Med_*Y*_ ”, where an undirected graph (“UndG_*X*_ ”) encodes the dependency pattern within ***X*** and, within the responses, an outcome (*Y*_3_) is completed mediated (“Med_*Y*_ ”) by another response (*Y*_2_) which, in turn, is affected by a different exposure (*X*_1_). Although there is another partial mediation between *X*_1_ and *Y*_3_ through *X*_2_, this mediation happens within ***X***, so it does not affect the definition of complete mediation within ***Y***. (**B**) Simulated scenario “DAG_*X*_ -Med_*Y*_ ”, where a topologically ordered DAG within the exposures (“DAG_*X*_”) is simulated. Specifically, in the example depicted, a fork structure is simulated, *i*.*e*., *X*_3_ affects both *X*_1_ and *X*_2_. A complete mediation is still considered within the responses. (**C**) Simulated scenario “UndG_*X*_ - DAG_*Y*_”. Here, the dependency structure between the individual-level responses is obtained by simulating a topologically ordered DAG (“DAG_*Y*_”). Specifically, a chain structure is considered, *i*.*e*., *Y*_1_ affects *Y*_2_ which, in turn, affects *Y*_3_, whereas an undirected graph encodes the dependency pattern within ***X***. (**D**) Simulated scenario “DAG_*X*_ -DAG_*Y*_ ”, where two topologically ordered DAGs are simulated within the exposures (fork structure) and outcomes (chain structure), respectively.

**Figure 3.**
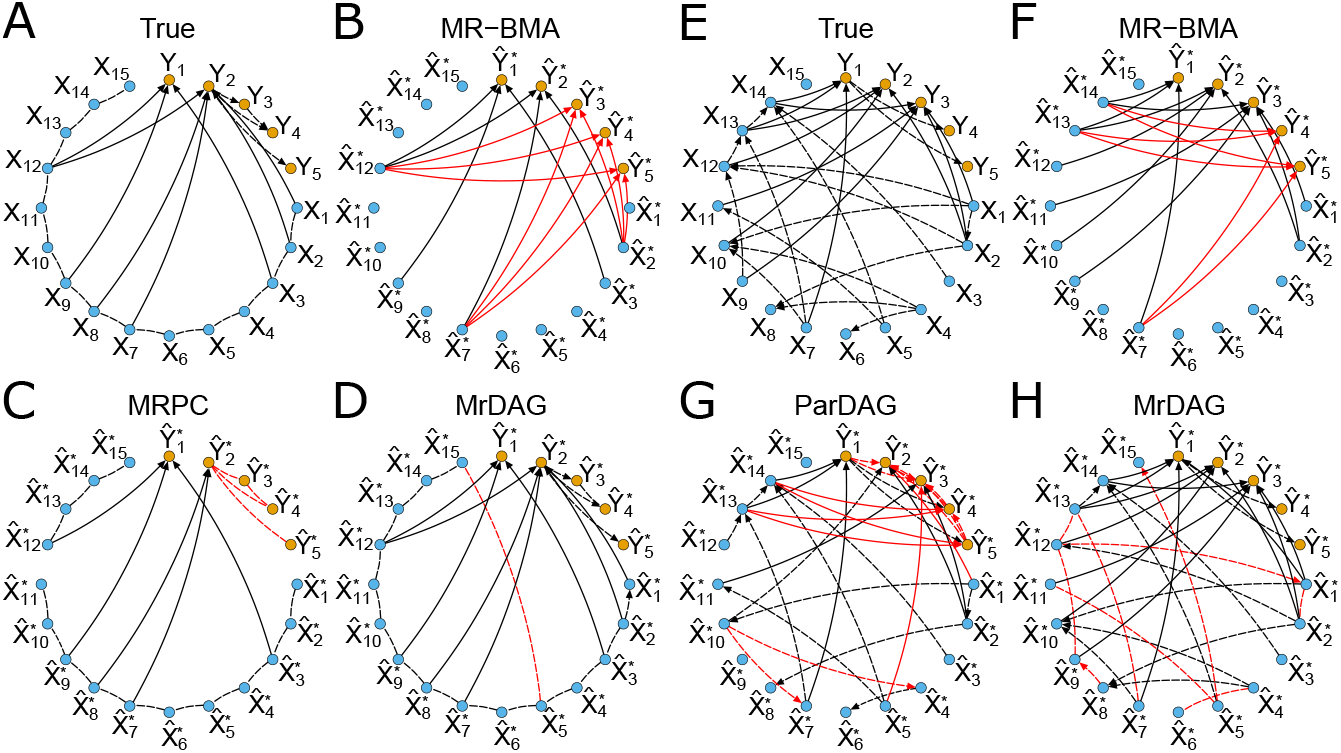
Examples of unconfounded dependency structure simulated at the individual-level data and estimated by using summary-level statistics within the exposures, the outcomes and between them in two different scenarios. In each panel, individual-level out-comes ***Y*** = (*Y*_1_, …, *Y*_5_) and exposures ***X*** = (*X*_1_, …, *X*_15_) as well as genetically predicted outcomes 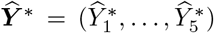 and exposures 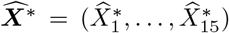 are represented with orange and blue nodes, respectively. Directed edges indicate dependency relations, while undirected edges denote partial correlation. Dashed lines depict the true (unconfounded by *U*) and estimated dependency structure within the exposures and the outcomes, while solid lines indicate true and estimated causal effects between them. Red colour denotes false positives, either falsely detected effects (regardless of the directionality) or wrong directionality of the edges. Besides the proposed model, alternative methods considered: Mendelian randomization with Bayesian Model Averaging (MR-BMA) [1], Mendelian randomization with PC algorithm (MRPC) [28], Partition-DAG (ParDAG) [30]. We report the results of MR-BMA and MrDAG without any threshold on the marginal posterior probability of inclusion (mPPI) and the posterior probability of edge inclusion (PPeI), respectively. MRPC Partially Directed Acyclic Graphs (PDAGs) are obtained by specifying the type I error rate for the conditional independence test at *α* = 0.01. ParDAG results are the solutions of causal effects estimation with Lasso penalisation set at *λ* = 0.9 after partitioning the traits into two groups and enforcing a constraint on the orientation of the edges between the exposures and the outcomes. (**A**-**D**) Single replicate of the simulated scenario UndG_*X*_ -DAG_*Y*_, where an undirected graph encodes the dependency pattern within ***X*** and a DAG represents the dependency relations within ***Y*** along with the simulated causal effects from the exposures to the outcomes, resulting in an overall partially oriented DAG. In this scenario, the strength of correlation between consecutive ***X*** is set at *r*_*X*_ = 0.6, and then decreases exponentially for non-consecutive exposures, and the average level of the mediation parameters within ***Y*** is set at *m*_*Y*_ = 1. (**E**-**H**) Single replicate of the simulated scenario DAG_*X*_ -DAG_*Y*_, where two topologically ordered DAGs have been independently simulated within ***X*** and ***Y*** along with the simulated causal effects from the exposures to the responses, resulting in an overall fully-oriented DAG. In this scenario, the average level of mediation parameters for ***X*** and ***Y*** are set at *r*_*X*_ = 0.6 and *m*_*Y*_ = 1, respectively.

Taken together, in scenarios (ii) and (iv), the overall individual-level DAGs, obtained by combining two different simulation strategies for ***X*** and ***Y***, are fully oriented while in scenarios (i) and (iii) the overall DAGs are partially oriented. Details regarding the parameters *ψ*_*X*_ and *ψ*_*Y*_, the simulated levels of the effects of the unmeasured confounder *U* on the responses and the outcomes, ***B***_*X*_, the simulated levels of the genetic effects on the exposures, and **Γ**_*X*_ and **Γ**_*Y*_, the simulated levels of the mediation parameters within the exposures and the outcomes are presented in Methods. Finally, all simulations are replicated 25 times and initialised with a different random seed.

We compare MrDAG with published multivariable MR methods and their software implementations excluding from the comparisons naïve one-exposure and one-outcome MR models since it has been shown that they are outperformed by multivariable MR methods when there is measured pleiotropy among exposures [3]. Specifically, we consider Mendelian randomization with Bayesian Model Averaging (MR-BMA) [1] an MVMR algorithm which allows for many exposures to be included, but does not model explicitly the dependency relations within the exposures [3]. MR-BMA estimates the sparse direct causal effects between the exposures and one outcome providing the marginal posterior probability of inclusion (mPPI) along with the posterior mean of the causal effects. We treat MR-BMA as the baseline algorithm for the comparisons since it analyses one outcome at-a-time. Secondly, we include Mendelian randomization with PC algorithm (MRPC) [28], which combines instrumental variables with the PC algorithm [29] for DAG estimation. At a specified type I error rate for the conditional independence test, MRPC returns the estimated Partially Directed Acyclic Graphs (PDAGs) [19] (see Methods) in which some undirected edges are present along with the directed ones as well as the *p*-values of all conditional independence tests. For a given PDAG detected by MRPC in each replicate and scenario, we utilise [27] to estimate the causal effects between the exposures and outcomes. Finally, Partition-DAG (ParDAG) [30] provides a solution to the structure learning problem once the summary-level statistics have been partitioned into two groups and the orientation of the edges from the exposures to the outcomes has been enforced. ParDAG computes the causal effects estimates under Lasso regularisation. It has not been combined with instrumental variable estimation and applied to genetic data to date. All methods use summary-level statistics as input after IVW. Finally, for each method and algorithmic implementation, details of the parameter settings are provided in Supplementary Information.

Regarding the evaluation criteria, we use a precision-recall curve (PRC) that shows the relationship between precision (*i*.*e*., positive predictive value, on the *y*-axis) and recall (*i*.*e*., sensitivity, on the *x*-axis) for every possible cut-off and it is not impacted by the over-representation of null effects. See Supplementary Information for a detailed discussion regarding how we implemented a fair comparison between the methods considered.

Finally, to evaluate the quality of the causal effects estimation, we calculate the sum of squared errors (SSE), defined as the sum of the squared differences between the estimated and the simulated causal effect. In contrast to the evaluation of the recovery obtained by each method of the simulated dependencies within the exposures, the outcomes and between them, we do not report the SSE of the mediation parameters **Γ**_*X*_ and **Γ**_*Y*_ since they are considered nuisance parameters in the proposed model (see Supplementary Information).

### MrDAG more accurately detects unconfounded dependency relations within the exposures and the outcomes and between them

Figure 3 presents the results of MrDAG and alternative methods for one replicate of the simulated scenario UndG_*X*_-DAG_*Y*_ (Figures 3A-D) and DAG_*X*_-DAG_*Y*_ (Figures 3E-F) for a particular choice of the parameters *r*_*X*_ = 0.6 and *m*_*Y*_ = 1 used in the simulation study to control the average value of the mediation parameters **Γ**_*X*_ within the exposures and **Γ**_*Y*_ within the outcomes, and *ψ*_*X*_ = 2 and *ψ*_*Y*_ = 1 for the level of confounding on the exposures and the outcomes, respectively (see Methods).

The general performance of competing algorithms is already apparent from it. In scenario UndG_*X*_-DAG_*Y*_, if a causal effect is simulated from an exposure to an outcome and there are dependency relations from this outcome to other responses (Figure 3A), MR-BMA adds erroneously causal effects to all linked responses with severe FP inflation (Figure 3B, FPs between 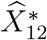 and 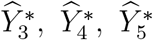 depicted in red). On the other hand, MR-BMA estimates neither the dependency pattern within ***X***, since (partial) correlation between summary-level exposures is assumed in the model [3] but not estimated, nor the dependencies within ***Y*** since MR-BMA considers one response at-a-time. MRPC infers correctly most of the dependencies within ***X***, but it does not have the power to detect all simulated causal effects **Θ** at the specified type I error rate for the conditional independence test (*α* = 0.01) with a few FNs (Figure 3C, FNs between 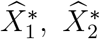 and 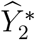) and well as FPs within 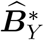 (FPs between 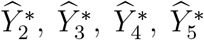, where bidirectionally is erroneously detected). MrDAG performs better than alternative methods to detect both directed and bidirected edges with only one FP between 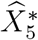 and 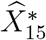 (Figure 3D).

Similar comments can be made for a particular replicate of scenario DAG_*X*_-DAG_*Y*_, although in this scenario the dependency patterns are more complex since a topological ordered DAG is simulated also within the outcomes (Figure 3E). MrDAG confirms its good performance except for the directionality of the dependency relations within ***X***, where bidirectional edges are found with a few FPs (Figure 3H, FPs between 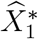 and 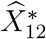 and between 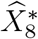 and 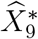).

Figure 4 generalises the results depicted in Figure 3, averaging the results over 25 replicates of the simulated scenarios UndG_*X*_-DAG_*Y*_ (Figures 4A-C) and DAG_*X*_-DAG_*Y*_ (Figures 4D-F) with the same parameters setting used in Figure 3. The results are presented separately for the simulated dependency structures from the exposures to the outcomes (Figures 4A and D), within the exposures (Figures 4B and E) and within the outcomes (Figures 4C and D), respectively.

**Figure 4.**
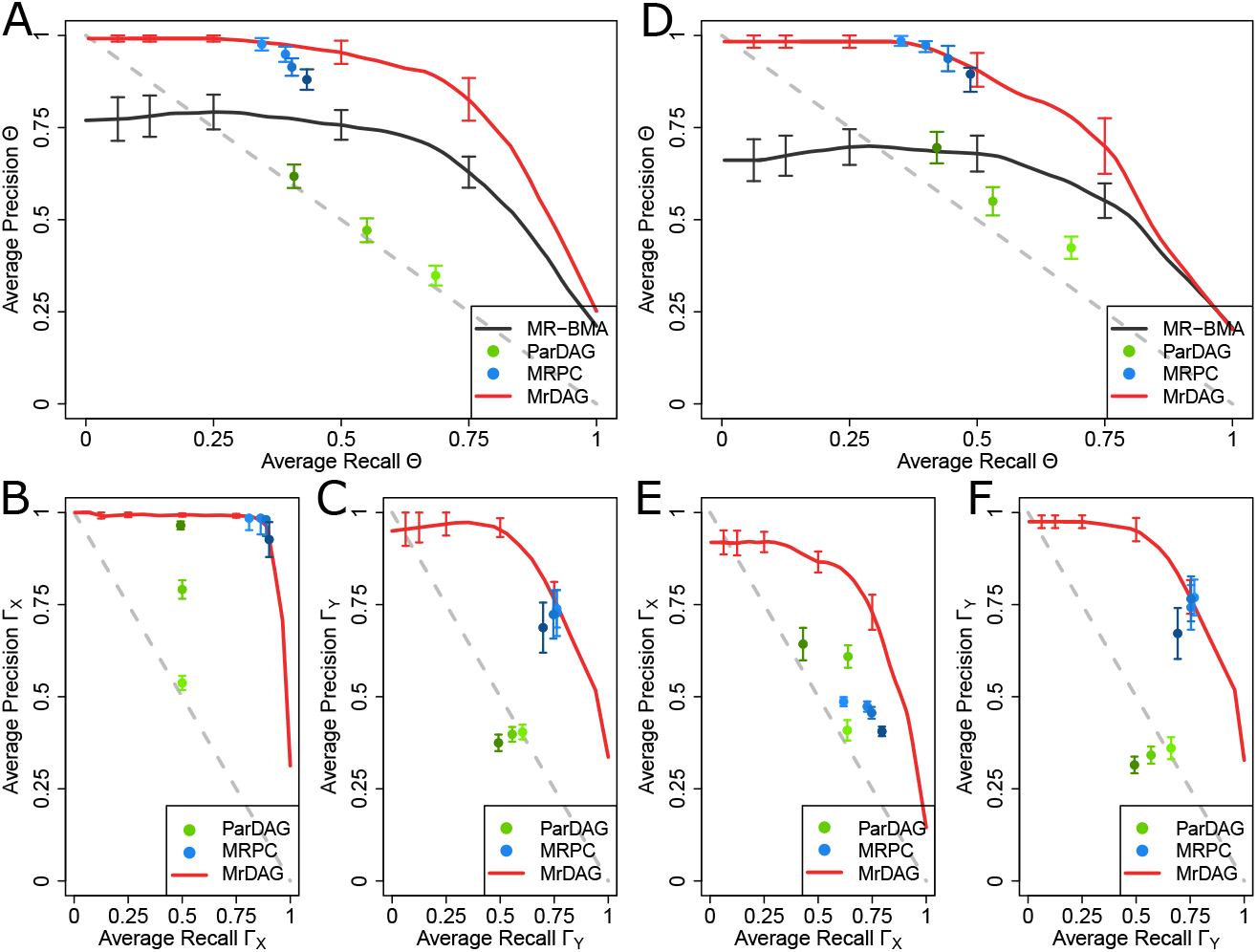
Precision Recall Curves (PRCs) for all methods considered in the simulated scenarios UndG_*X*_ -DAG_*Y*_ and DAG_*X*_ -DAG_*Y*_. show recall (= sensitivity = TP*/*(TP + FN)) in the *x*-axis and precision (= positive predictive value = TP*/*(TP + FP)) in the *y*-axis with TP = True Positive, FN = False Negative and FN = False Positive averaged over 25 replicates in each scenario. In scenario UndG_*X*_ -DAG_*Y*_ (**A**-**C**), the strength of correlation between consecutive ***X*** is set at *r*_*X*_ = 0.6, and then it decreases exponentially for non-consecutive exposures, and the average level of the mediation parameters within ***Y*** is set at *m*_*Y*_ = 1, while in scenario DAG_*X*_ -DAG_*Y*_ (**D**-**F**), the average level of the mediation parameters within ***X*** and ***Y*** is set at *r*_*X*_ = 0.6 and *m*_*Y*_ = 1, respectively. For details, see Methods. In both scenarios, the results are presented separately for the simulated dependency structures from the exposures to the outcomes (A and D), within the exposures (B and E) and the outcomes (C and D), respectively. Vertical bars in each PRC, at specific recall levels 0.0625, 0.125, 0.25, 0.50 and 0.75, indicate standard error. For the MRPC algorithm, type I error rate for the conditional independence test is set at *α* = {0.01, 0.05, 0.10, 0.20} (from light-to dark-blue dots) and for the ParDAG algorithm we specify three different values for the Lasso penalisation *λ* = {0.5, 0.7, 0.9} (from light-to dark-green dots). See Supplementary Information for details.

On average, MRPC and MrDAG have good performance in both simulated scenarios (Figures 4A and D). MRPC best results are obtained at a stringent type I error rate *α* = 0.01 for the conditional independent tests (blue dots) although they are quite similar across different values of *α* and thus robust to this choice. However, it fails to detect the simulated dependency pattern within ***X*** in scenario DAG_*X*_-DAG_*Y*_ (Figure 4B). The performance of MR-BMA can be only evaluated for the detection of the causal effects from the exposures to the outcomes (Figures 4A and D). As we noticed above, the large number of FPs degrades the results of this method which was not developed to deal with multiple related responses.

The performance of ParDAG is the worst among the methods considered for all types of designed relationships, slightly better within the exposures (Figures 4B and E) and between the exposures and outcomes (Figures 4A and D) and worse within the outcomes (Figures 4C and F). Since ParDAG detects only directed edges, in Figure 4B, where partial correlation between exposures is simulated, the method has 50% recall rate. The results seem also quite different according to the penalty parameter *λ*.

MrDAG has a strong performance in both scenarios. In contrast to MR-BMA, in scenario DAG_*X*_-DAG_*Y*_ (Figures 4A and C) there is only a small reduction of the precision in the estimation of the dependency relations between the exposures and the outcomes, and within the latter, compared to the scenario UndG_*X*_-DAG_*Y*_ (Figures 4D and F).

The comments above can be extended to the scenarios where the relationships within outcomes are completely mediated (UndG_*X*_-Med_*Y*_ depicted in Supplementary Figures 2A-C, and DAG_*X*_-Med_*Y*_ shown in Supplementary Figures 2D-F). In these scenarios, the mediation within the outcomes is easier to detect (Supplementary Figures 2C and F) than a topologically ordered DAG simulated within ***Y***.

Supplementary Figure 3 shows the results of the AUCPR to detect the causal effects **Θ** and the sensitivity of the methods to different specifications of *r*_*X*_ and *m*_*Y*_. MrDAG confirms to be uniformly the best method with stable AUCPR for any combination of *r*_*X*_ and *m*_*Y*_ with similar AUCPR when partial correlation or a topological ordered DAG is simulated within ***X*** (Supplementary Figures 3A and B). MR-BMA performs well, especially in the scenario UndG_*X*_-Med_*Y*_ (Supplementary Figure 3A) which is the scenario that is most compatible for this method as well as in scenario DAG_*X*_-Med_*Y*_ (Supplementary Figure 3C), where its performance slightly decreases. Both MRPC and ParDAG seem to be less precise at higher levels of *r*_*X*_ irrespective of the simulated scenario, with ParDAG also influenced by the value of *m*_*Y*_. Similarly, Supplementary Figures 4) and 5 show the sensitivity of the algorithms to detect the simulated patterns within ***X*** and within ***Y*** for different specifications of *r*_*X*_ and *m*_*Y*_.

### MrDAG improves the estimation of the causal effects over existing methods

Figure 5A shows the Sum of Squares Error (SSE) of the causal effects **Θ** between the exposures and the outcomes for all methods considered in the simulated scenario UndG_*X*_-DAG_*Y*_ and Figure 5B for the simulated scenario DAG_*X*_-DAG_*Y*_ across 25 replicates in each scenario with the same parameter setting and implementation of algorithms described above. For MRPC and ParDAG algorithms, we only show the results obtained at type I error rate for the conditional independence test *α* = 0.01 and Lasso penalisation *λ* = 0.9, respectively. These values provide the best results for the two algorithms as shown in Figure 4 and Supplementary Figure 2.

**Figure 5.**
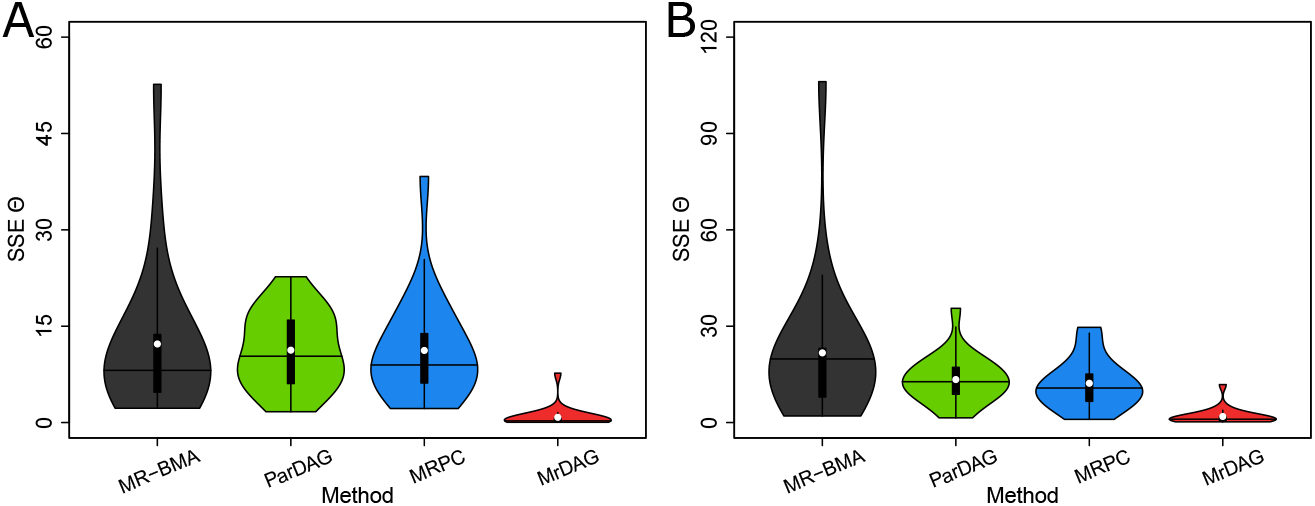
Violin plots of the Sum of Squares Error (SSE) of the causal effects Θ between the exposures and the outcomes for all methods considered in the simulated scenarios UndG_*X*_ -DAG_*Y*_ and DAG_*X*_ -DAG_*Y*_ across 25 replicates in each scenario. (**A**) In scenario UndG_*X*_ -DAG_*Y*_, the strength of correlation between consecutive ***X*** is set at *r*_*X*_ = 0.6, and then it decreases exponentially for non-consecutive exposures, and the average level of the mediation parameters within ***Y*** is set at *m*_*Y*_ = 1. (**B**) In scenario DAG_*X*_ -DAG_*Y*_, the average level of the mediation parameters within ***X*** and ***Y*** is set at *r*_*X*_ = 0.6 and *m*_*Y*_ = 1, respectively. For details, see Methods. In each violin plot, the vertical black thick line displays the interquartile range, the black horizontal line denotes the median and the white dot the mean. For MRPC and ParDAG algorithms, we only show the results obtained at type I error rate for the conditional independence test *α* = 0.01 and Lasso penalisation *λ* = 0.9, respectively. These values provide the best results for the two algorithms as shown in Figure 4 and Supplementary Figure 2.

MrDAG has the lowest SSE mean and median (white dots and horizontal black line, respectively) in both scenarios. As expected, when a topological ordered DAG is simulated within the exposures (Figure 5B), the violin plot have a wider range, showing more variable results, although the median is almost similar to the scenario with simulated partial correlation within ***X*** (Figure 5A). Alternative methods have larger SSE.

Similar comments can be made for simulated scenarios UndG_*X*_-Med_*Y*_ (Supplementary Figure 6A) and DAG_*X*_-Med_*Y*_ (Supplementary Figure 6B), where a complete mediation is considered within the outcomes. MrDAG is confirmed as the best method.

We conclude this section by inspecting the sensitivity of the SSE of the causal effects between the exposures and the outcomes for different values of the average level of the mediation parameters *r*_*X*_ and *m*_*Y*_. The estimation of the causal effects displayed in Supplementary Figure 7 shows that both MR-BMA and MRPC depend on the combination of *r*_*X*_ and *m*_*Y*_ with similar performance when a complete mediation is simulated (Supplementary Figures 7A and C) (Supplementary Figures 7B and D). Compared to the other methods, MrDAG is not only the best, but it is rather insensitive to different levels of the mediation parameters within ***X*** and ***Y***.

### Real data application: The impact of lifestyle and behavioural traits on mental health

We apply MrDAG to investigate its ability to detect the effect of lifestyle and behavioural exposures on the risk of mental health phenotypes as well as potential forms of interventions for their prevention. As exposures, we chose seven lifestyle and behavioural traits that have previously been investigated for their effects on mental health, including educa-tion (in years) (EDU), physical activity (PA), sleep duration (SP), alcohol consumption (ALC), lifetime smoking index (SM) and leisure screen time (LST). As outcomes, we select seven mental health phenotypes, including major depressive disorder (MDD), anorexia nervosa (AN), attention deficit hyperactivity disorder (ADHD), bipolar disorder (BD), autism spectrum disorder (ASD), schizophrenia (SCZ) and cognition (COG). See Supplementary Table 1 for the description of the summary-level statistics, the data sources, the number of IVs for each trait and Methods for the pre-processing steps. In a separate analysis, we also investigate the reverse direction, *i*.*e*., whether the same mental health phenotypes have an impact on the group of lifestyle and behavioural traits by selecting IVs for the mental health phenotypes, see Methods for the respective pre-processing steps.

Figure 6 presents the results of MrDAG. In particular, Figures 6A and C show the estimated posterior probability of edge inclusion (PPeI) (12) after structure learning and Figures 6B and D the posterior causal effects (95% credible intervals (CI)) between the exposures and the outcomes. Results on PPeI (and the posterior causal effects) are not thresholded and sparsity is enforced by assigning a prior on the number of expected edges. We set it at *π*^edge^ = 0.16, *i*.*e*., we expect *a priori* one edge for each of the 13 traits, see Methods and Supplementary Information.

**Figure 6.**
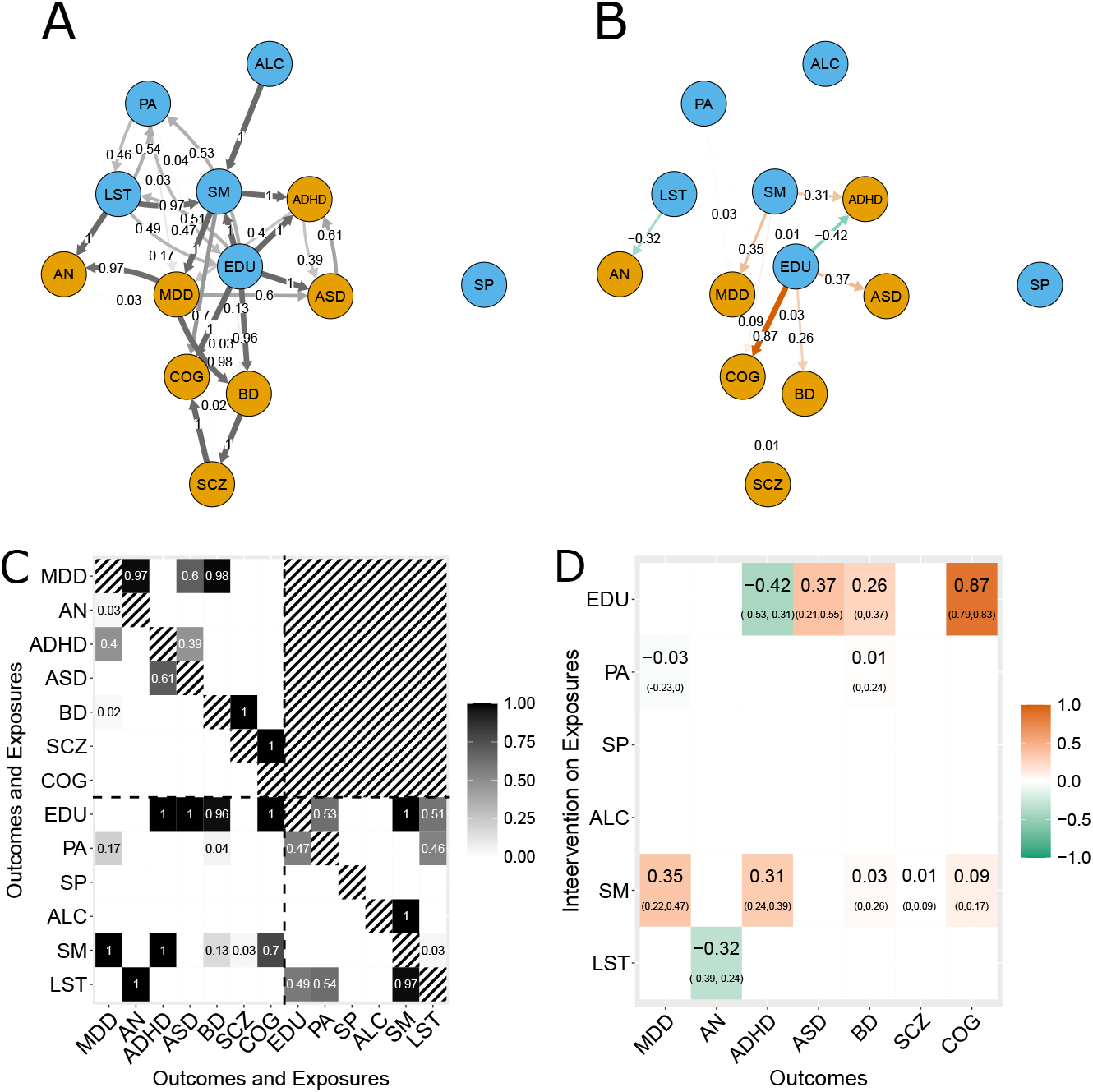
Results of MrDAG algorithm regarding how lifestyle and behavioural exposures impact mental health outcomes. (**A**) PDAG of the posterior probability of edge inclusion (PPeI) within the exposures (lifestyle and behavioural traits, blue nodes), the outcomes (mental health phenotypes, orange nodes) and between them. Undirected edges are represented as bidirectional edges, see, for instance, edges between PA (physical activity) and LST (leisure screen time) or ASD (autism spectrum disorder) and ADHD (attention deficit hyperactivity disorder). Neither reverse causation from the outcomes to the exposures nor feedback loops are allowed. (**B**) Posterior causal effects on the outcomes (orange nodes) under intervention on the exposures (blue nodes). Red and green edges indicate positive and negative posterior causal effects, respectively. (**C**) Posterior probability of edge inclusion (PPeI) for each combination of outcomes (mental health phenotypes) and exposures (lifestyle and behavioural traits). Horizontal and vertical dotted lines separate the exposures (bottom-right submatrix) from the outcomes (top-left submatrix). PPEIs between exposures and outcomes are depicted in the bottom-left submatrix. Neither reverse causation (top-right submatrix) nor feedback loops (main diagonal) are allowed (black-white strips). (**D**) Posterior causal effects (95% credible intervals) on the outcomes (*y*-axis) under intervention on the exposures (*x*-axis).

As shown in Figures 6C and D, there is one distinct exposure (LST) and two key shared exposures with important down-stream effects on mental health phenotypes, which are EDU and SM on which we focus our discussion. For each of them, we also describe how MrDAG can disentangle complex dependency relations within the exposures and the outcomes and detect (partial or complete) mediation which prevents spurious findings.

As could be expected due to its centrality in the global health agenda [31] and the high level of confounding of this phenotype with other genetically associated biological, behavioural and socioeconomic traits, genetically predicted EDU shows the most inter-exposure and exposure-outcome dependency relations (Figure 6C bottom part). Previous work has supported the broad mental health implications of education [32]. First, in keeping with previous findings [33, 34, 35, 36], our results show that EDU has a positive causal effect on COG, it is causally associated with an increased liability to ASD and BD as well as with a lower liability to ADHD. CIs show that the causal association with BD is markedly skewed to the right. In contrast, EDU has no effects on SP, the amount of ALC, or the liability to MDD [33], AN [37], or SCZ [36] (Figure 6D). Second, we investigate the detected dependency relations of EDU with other exposures that contribute to the reported causal associations. We find bidirectional relationships between genetically predicted EDU, PA and LST consistent with a large literature [33, 38]. Dependency relations have been also identified between EDU and SM [33, 39]. Supported by the existing literature, these results confirm the ability of MrDAG to disentangle complex relationships that exist between interrelated exposures.

We find that SM is second only to EDU in its causal association with several outcomes. Specifically, SM associates with an increased liability to MDD and ADHD as previously reported [40, 41]. It is also associated with COG, BD and SCZ, although these causal effects are small and CIs are skewed to the right. As discussed above, we also check the detected dependency relations of SM with other exposures. SM is related to PA as documented in epidemiological studies [42] and in standard MR analysis [43], the latter for objectively assessed average activity and number of cigarettes per day, respectively. Moreover, MrDAG appropriately identifies the relationship between ALC and SM, but not *vice versa*. In a recent MR publication [44], the opposite causal association is observed. However, in contrast to [44] who conceptualize SM with smoking initiation, we use a lifetime smoking index [40] which captures smoking duration, heaviness and cessation.

As important as the discussion of existing causal associations between the exposures and the outcomes, it is similarly insightful to discuss the absence of causal effects, especially those relationships that are reported in the literature or found by standard (one exposure and one outcome) MR models. For example, we do not replicate all previous evidence for positive causal effects of liability to SM on mental health phenotypes. Though we find a strong causal effect of SM on MDD [40], we do not find the same strong effect of SM on SCZ [40] as observed in observational studies [45, 46]. By looking at Figure 6C, this might be due to pleiotropic effects that have been identified by MrDAG within the mental health phenotypes. In line with prior findings, evidence from MrDAG supports dependency relations between genetic liability to MDD and AN, ASD and BD [47] as well as between genetic liability to BD and SCZ [48]. Lastly, in keeping with prior findings of possible bidirectional ASD-ADHD relationships [49], we observed genetic dependency relations between ASD and ADHD, and *vice versa*. These results suggest that the genetic effects of SM on SCZ can be mediated by pleiotropic effects within the responses. By considering the results above, we hypothesise that the SM to SCZ relationship is partly mediated first by MDD and then by BD. Moreover, there is another path that goes from the genetically predicted level of SM to SCZ through a positive weak causal association identified by MrDAG between SM and BD [50]. Both genetic paths are illustrated in Figure 6A. Conditionally on these relationships that are not considered in standard MR or MVMR, MrDAG does not detect a strong causal effect between SM and SCZ.

We further note that the causal effect of SM on ADHD is both direct and indirect, the latter mediated first by MDD and then by ASD. Thus, our analysis pinpoints the important role of MDD which partly or entirely accounts for many causal pathways within mental health phenotypes and their causal exposures. This might be due to the potentially high levels of confounding and non-specific genetic associations present in the original MDD GWAS [51, 52] as well as the high levels of symptom-level and therefore diagnostic overlap between MDD and all other psychiatric disorders [53]. Nonetheless, the implications of our results, assuming the validity of all GWAS findings, are that prevention and/or therapeutic intervention on MDD [54] can have a cascade of important effects for the prevention of several mental health phenotypes.

To investigate this hypothesis, Supplementary Figures 10A and B show the results of MrDAG when MDD is removed from the list of outcomes. Regarding the causal association between SM and ADHD, it is still present with the same strength and similar CI depicted in Figure 6D, suggesting that the indirect effect mediated first by MDD and then by ASD is negligible. Supplementary Figure 10B also shows that, after removing MDD, the genetically predicted SM is positively associated with SCZ as reported in the literature. Combined with our main findings, this result indicates that the absence of a link between SM and SCZ in the MrDAG model is likely due to the mediation of MDD and BD.

The risk of detecting spurious shared causal effects is very high when a standard MR method is used separately on each trait as well as when multiple exposures are considered for each outcome [1]. This problem has been highlighted in the simulation study and visually presented in Figures 3B and F. In Supplementary Table 2 we show the results MR-BMA algorithm when applied to the same data set. We notice an overestimation of the causal effects since MR-BMA tries to ascribe the whole effects to the exposures and, as expected from the simulation study, it also detects many more associations than MrDAG.

We conclude the analysis by assessing the validity of the results obtained by MrDAG. We divide this internal check into sensitivity to hyper-prior specification and robustness of structure learning. Regarding the first point, Supplementary Figure 11 show that the posterior causal effects as well as the 95% CIs for different values of the *a priori* probability of edge inclusion are not influenced by this choice. For the second internal check, we bootstrap MrDAG repeatedly on the data [55] (see Supplementary Information). In Supplementary Figures 12 we present the bootstrap frequency of edge inclusion for each permitted combination of exposures and outcomes and the scatterplot of the posterior probability of edge inclusion (PPeI) against the bootstrap frequency of edge inclusion. The results show that there is a satisfactory agreement between a single run of the algorithm and the bootstrap results for the causal associations. Extended results are presented in Supplementary Information.

For completeness, we have also tested reverse causation by selecting genetic variants to be associated with the mental health phenotypes. Figure 7 and Supplementary Figure 9 show the results of the analysis to detect the impact of mental health phenotypes on lifestyle and behavioural traits, where, besides the positive causal effect of genetically predicted COG on EDU [56], smoking is causally affected by the genetic liability to MDD [40] and ADHD, the latter well-documented in epidemiological studies [57] and recently confirmed in a randomised clinical trial of smoking cessation [58].

**Figure 7.**
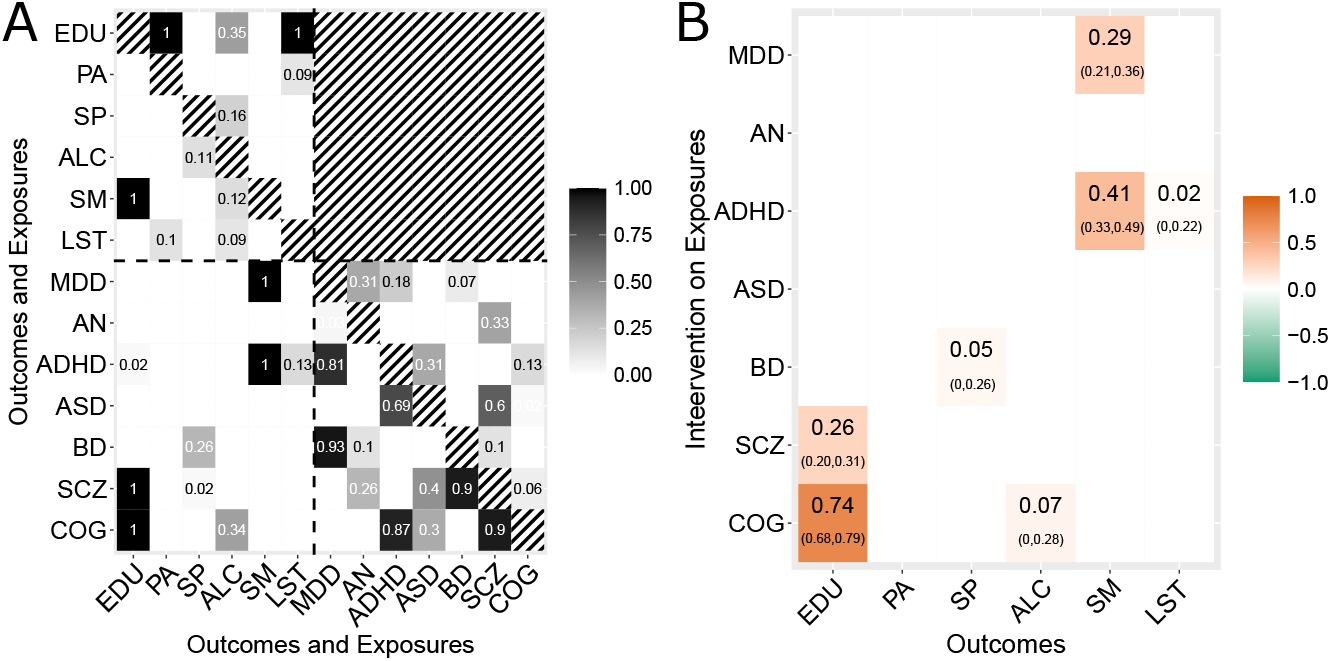
Results of MrDAG regarding how liability to mental health phenotypes affects lifestyle and behavioural traits. (**A**) Posterior probability of edge inclusion (PPeI) for each combination of outcomes (lifestyle and behavioural traits) and exposures (mental health phenotypes). Horizontal and vertical dotted lines separate the exposures (bottom-right submatrix) from the outcomes (top-left submatrix). PPEIs between exposures and outcomes are depicted in the bottom-left submatrix. Neither reverse causation (top-right submatrix) nor feedback loops (main diagonal) are allowed (black-white strips). (**B**) Posterior causal effects (95% credible intervals) on the outcomes (*y*-axis) under intervention on the exposures (*x*-axis).

## Discussion

Here, we have introduced MrDAG, the first Bayesian causal graphical MR model for multivariable and multiresponse that can detect dependency patterns within the exposures as well as within the outcomes thus allowing for a more precise estimation of the causal effects from the exposures to the outcomes. We showcased the advantage of the proposed method in a comprehensive simulation study and its utility in detecting how lifestyle and behavioural traits interact to cause mental health phenotypes, and *vice versa*. In the real data application, we highlighted how MrDAG can recover more information on the genetic paths that link exposures to outcomes compared to existing MR methods that ignore these dependency relations. Specifically, we highlighted education and smoking as key effective points of intervention given their distinct downstream effects on multiple mental health phenotypes.

These insights are possible since three methodological advances are considered in MrDAG. First, in structure learning, the hypothesis of no unobserved confounding is a fundamental underlying assumption. This assumption, known as causal sufficiency, is difficult to justify in real data applications and its violation produces biased results. By using IVs within the MR paradigm, we bypass the need to remove the effects of the unmeasured confounder from the individual-level data [21]. Instead, we solve this problem by employing genetically predicted exposures and outcomes which depend only on the genetic variants chosen as IVs. Genetically predicted exposures are key in the derivation of the two-stage least square causal effect estimator [25], but in MrDAG we have extended it to include genetically predicted outcomes. On both predicted traits, we perform DAG exploration to learn the unconfounded dependency relations that exist within the exposures, the outcomes and between them. Our second contribution is the estimation of causal effects under intervention on the exposures conditionally on a given DAG. We showed that they can be estimated based on Pearl’s interventional calculus [7]. Moreover, differently from [59] and its application in the MRPC algorithm [60], the estimation of the causal effects is averaged over the visited graphical models [61], thus taking into account the uncertainty regarding the graphs that best portray the dependency structure in a given data set. Third, MrDAG allows the possibility of including domain-knowledge relations between the traits. In the designed MrDAG model, constraints between the exposures and the outcomes descend directly from the MR paradigm. Our Bayesian implementation of structure learning under restrictions offers clear advantages over alternative methods [30]. Although not discussed here, other restrictions can be straightforwardly included, for instance, known relations regarding disease progression or time-dependent outcomes, *e*.*g*., smoking initiation and cessation [62].

In the real data application, while the use of existing summary-level statistics of genome-wide association studies facilitates the integration of diverse phenotypes measured in different cohorts, we are also limited by the biases suffered by the initial genome-wide association studies. Specifically, studies on mental health rely on the presence of a clinical diagnosis. Consequently, it is not truly the genetic liability of the disease itself as much as it is the probability of having access to diagnoses or treatment. Our findings on the relationship between higher genetically predicted educational attainment (EDU) and increased ASD and BD, but decreased ADHD risk provide an example of such bias. In these analyses, the predicted number of school years completed is unlikely to be causally implicated in the development of ASD traits. While the typical age of onset of ASD precedes the start of formal education (therefore unlikely to be caused by it), ASD-related traits are more likely to be recognized and referred, particularly in those who are undiagnosed or untreated, when individuals are within a schooling system where standardized testing and progress reports by peer comparison are performed. Moreover, current GWAS consider one trait or disease at-a-time and do not consider to what extent cases are comorbid with other diseases. Future GWAS on co-morbidity [63] may provide more fine-grained genetic associations allowing to disentangle some of these relationships. Alternatively, novel causal inference methodology designed for individual-level data in combination with large-scale biobank or cohort studies with genotype data could be used to triangulate evidence.

In conclusion, MrDAG represents an important step forward in how we can learn complex relationships among phenotypic traits and uncover causal pathways using ge-netic data. It provides analysts with the opportunity to derive a more comprehensive picture of causal mechanisms between complex phenotypes. The real data application is an example of the proposed holistic approach, where we leverage MrDAG and large-scale genome-wide association data to offer novel mechanistic insight into the causal behavioural determinants of mental health phenotypes to delineate between their overlapping pathophysiology and phenotypic presentation, toward translational progress in the field of mental health. Moving forward, MrDAG is ideally placed for the analysis of common causal exposures for multimorbid health conditions. This research into multimorbidity has been facilitated by the advent of large-scale biobanks being linked and followed up using electronic health records and routinely collected health care data. Using genotype data as genetic anchors offers a principled way for causal inference. MrDAG provides an addition to existing toolkits to map shared and distinct causes of disease, to understand trajectories, and to draw causal paths that link diseases.

## Methods

In the following, we denote with capital letters the random variables *Y*, *X, G* and *U* for the observed outcome, exposure, instrumental variable and unmeasured confounder, respectively, and with small letters *y, x, g* and *u* their corresponding observations. Multivariate random variables and corresponding observations are presented in bold. A marginal element of a vector of random variables is specified by a suitable subscript index, *e*.*g*., *Y*_*k*_, *k* ∈ *K* = {1, …, *q*}, *X*_*j*_, *j* ∈ *J* = {1, …, *p*}, and *G*_*i*_, *i* ∈ *I* = {1, …, *n*}. ***Y***_*\k*_ and ***Y***_*\j*_ consists of all the outcomes and exposures except those that are related to the *k*th response and *j*th exposure, respectively. Finally, vectors understood as columns vectors and matrices are indicated in bold, the latter also in capital letters.

We indicate with 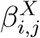 and 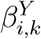 the effect of the genetic variant *i* ∈ *I* on the exposure *j* ∈ *J* and outcome *k* ∈ *K*, respectively, with 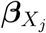 and 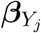 the *n*-dimensional vector of genetic effects on the *j*th exposure and *k*th outcome, respectively, and, finally, with ***B***_*X*_ and ***B***_*Y*_ the (*n × p*)- and (*n × q*)-dimensional matrices of the genetic effects on all exposures and outcomes. *θ*_*j,k*_ denotes the causal parameter of interest, *i*.*e*., the causal effect of *X*_*j*_ on *Y*_*k*_, and 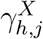 and 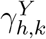 the mediation effect of *X*_*h*_ on *X*_*j*_, *h* ≠ *j* and *Y*_*j*_ on *Y*_*k*_, *h* ≠ *k*, respectively. **Θ, Γ**_*X*_ and **Γ**_*Y*_ indicate the corresponding (*p × q*)-, (*p × p*)- and (*q × q*)-dimensional matrices of the causal parameters of interest (**Θ**) and mediation parameters (**Γ**_*X*_ and **Γ**_*Y*_). The symbol “^” denotes the estimator of a parameter or its estimated value and “*” an IVW parameter.

Finally, let 𝒟 = (*V, E*) be a Directed Acyclic Graph (DAG), where *V* denote a set of vertices (nodes) and *E* = *V ×V* a set of directed edges, *i*.*e*., if (*z, v*) ∈ *E*, then (*z, v*) ∉ *E*. For a given DAG 𝒟, if *z* → *v*, then *z* is a parent of *v* and, conversely, *v* is a child of *z*. Moreover, if *z* → … → *v*, then *z* is an ancestor of *v* and *v* is a descendant of *z*. We denote the parent set of *v* in 𝒟 as pa_𝒟_(*v*) and *v* ∪ pa_𝒟_ (*v*) = fa_𝒟_ (*v*) the family of *v*. Unless otherwise stated, for ease of notation, we remove the subscript 𝒟.

In [5, 20, 64] key results regarding standard Mendelian randomization (single exposure with single instrumental variable and single outcome) are presented. Here, we use them to show that MrDAG is an extension of standard MR when (i) multiple exposures and outcomes are considered and (ii) the underlying dependency relations within and between them are not known (latent) and need to be estimated from the data. Technical details are provided in Supplementary Information.

### Multi-exposure and multi-outcome core conditions for instrumental variables

Let ***Y***, ***X*** and ***G*** be the *q*-, *p*- and *n*-dimensional vector of the outcomes, exposures and instruments (genotypes) random variables, respectively.

Let’s assume the following “multivariate core conditions” (MCC) for valid instrumental variables (IVs) which are the extensions of the core conditions that ***G*** has to satisfy in standard Mendelian randomization (MR) [5]:

(IV1) *G*_*i*_ ╨ *U*, ∀*i* ∈ *I, i*.*e*., *G*_*i*_ must be independent of *U* ;
(IV2) 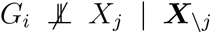, ∀*i* ∈ *I* and ∀*j* ∈ *J, i*.*e*., *G*_*i*_ must *not* be independent of *X*_*j*_ conditionally on ***X***_*\j*_;
(IV3) *G*_*i*_ ╨ *Y*_*k*_ | (***X***, *U*), ∀*i* ∈ *I* and ∀*k* ∈ *K, i*.*e*., *G*_*i*_ must be independent of *Y*_*k*_ conditionally on ***X*** and *U*.

The first multi-exposure and multi-outcome core condition (MCC) for instrumental variables is similar to the first CC in standard MR [5]. The second MCC imposes that *G*_*i*_ should be associated with *X*_*j*_ conditionally on the other exposures. The third MCC establishes that the instrumental variables and outcomes are conditionally independent given the exposures and the unmeasured confounder.

From the DAG 𝒟 involving ***Y, X, G*** and *U* that satisfies the MCC, the corresponding Markov properties say that *G*_*i*_ ╨ *U*, ∀*i* ∈ *I*, since *G*_*i*_ is not a descendant of *U* and *vice versa* and 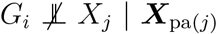, ∀*i* ∈ *I* and ∀*j* ∈ *J*, because *X*_*j*_ is a descendant of *G*_*i*_. The Markov property for the third MCC is *G*_*i*_ ╨ *Y*_*k*_ | (***Y***_pa(*k*)_, ***X***_pa(*k*)_, *U*), ∀*i* ∈ *I* and ∀*k* ∈ *K*, since *G*_*i*_ is a non-descendant of *Y*_*k*_ and (***Y***_pa(*k*)_, ***X***_pa(*k*)_, *U*) are the parents of *Y*_*k*_.

### Interventional distributions and causal effects estimation

The conditional dependencies associated with the multi-exposure and multi-outcome DAG 𝒟 lead to the following factorisation of the joint density of all random variables considered

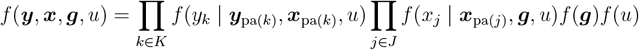

which is known as pre-intervention distribution and it is assumed to be faithful to the DAG [29], *i*.*e*., there are no conditional dependence relationships between the variables in the model that do not follow directly from the Markov properties.

The post-intervention distribution under intervention on the *h*th exposure sets to take the value 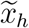 is obtained by the truncated factorisation [7]

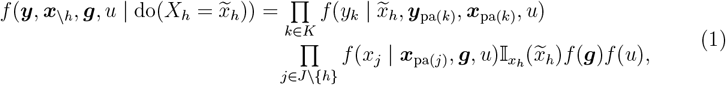

where 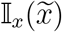 is the indicator function which is equal to one if 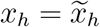 and zero otherwise. Graphically, the directed edges to *X*_*h*_ from its parents in ***X, G*** and *U* are removed.

A post-intervention distribution under intervention on the *h*the exposure is obtained from (1) by marginalising all variables but the selected outcome and the exposure on which an intervention is carried out

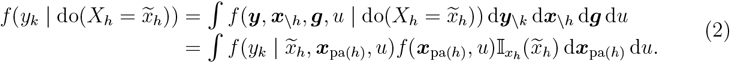

This result is derived from [7] and it follows directly from the Markov properties of the DAG. It establishes that the parents of the variable on which an intervention is carried out are the only variables that need to be measured to estimate the causal effect on an outcome [65].

The post-intervention distribution (2) can be summarised by taking the expectation and defining the causal effect of an intervention [59] as

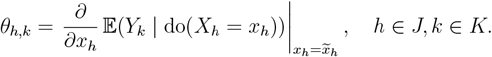

In Supplementary Information, we show the identifiability of the causal effect (Supplementary Proposition 2) and the derivation of its estimand in multiple exposures and multiple outcomes MR framework (Supplementary Proposition 3). We also show the consistency of the effects of the regressions of each outcome and exposure on ***G*** (Supplementary Proposition 1), *i*.*e*., the estimated genetic effects on the outcomes and exposures contain all information regarding the causal parameters of interest and the mediation parameters within the exposures and the outcomes.

Here, for a given DAG 𝒟, we report the IVW estimator of the causal effect of the intervention in *X*_*h*_ on *Y*_*k*_

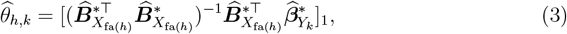

where the subscript indicates the first element of the solution of the linear least squares (LLS) regression since fa(*v*) = *v* ∪ pa(*v*), ***X***_fa(*h*)_ denotes the exposures that are the family of the exposure *X*_*h*_ under intervention, 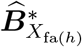 are the IVW estimated coefficients of the regressions of each exposure in ***X***_fa(*h*)_ on ***G*** and 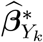 is the IVW estimated coefficient of a regression of *Y*_*k*_ on ***G***. (3) resembles the standard IVW estimator of the causal effect that approximates the estimate that would have been obtained if individual-level data were available [3]. However, in contrast to general proposed solutions in MVMR, in (3) the set of regressors is with regard to the family of the exposure under intervention.

### Dependency structure under the effect of unmeasured confounders

To estimate (3), structure learning of the graphical models needs to be performed to detect the parents ***X***_pa(*h*)_ of the exposure *X*_*h*_ under intervention. However, structure learning assumes causal sufficiency [21], *i*.*e*., it requires that there are no hidden (or latent) variables that are common causes of two or more traits. Instead, here we explicitly assume that an unmeasured confounder *U* acts on both outcomes and exposures.

Links between the genetic correlation and MR causal effect estimate have been already discussed in [23]. Here, we provide further connections with genetic covariance [24] which is key to show that, by working with summary-level statistics, it is possible to recover the dependency structure between the corresponding traits in the original (individual-level) data unconfounded by *U*.

Let’s assume that the genetic effect on a phenotypic trait is linear and consider two traits

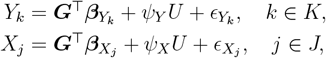

where ***G*** is a set of genetic variants, either spanning the whole genome, or region(s)- specific or selected to be associated with a trait, 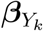 and 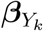 are the genetic effects, *U* is an unmeasured confounder that affects both traits with *ψ*_*Y*_ and *ψ*_*X*_ the effects sizes and 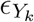 and 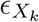 are white noises which can be interpreted as environmental effects. We assume that ***G*** ╨ *U* and, similarly, 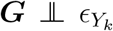 and 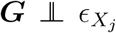. Finally, we assume that 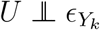 and 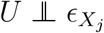, *i*.*e*., the unmeasured confounder *U* exerts its effect on both traits and it is distinct from other environmental factors. Under this model, the phenotypic covariance is

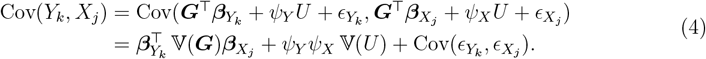

The phenotypic covariance can be decomposed into 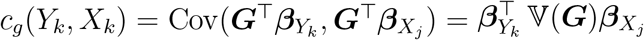, the genetic covariance between the two traits, *i*.*e*., the covariance between the genetic components of the two traits, 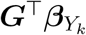 and 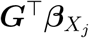, and the environmental covariance, *i*.*e*., the covariance between the environmental effects of two traits that we have split into the effect of the unmeasured confounder, *c*_*u*_(*Y*_*k*_, *X*_*k*_) = *ψ*_*Y*_ *ψ*_*X*_ 𝕍(*U*), and other environmental factors, 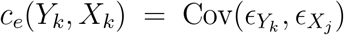. If the environmental factors are trait-specific since *U* includes all common confounding factors, *c*_*e*_(*Y*_*k*_, *X*_*k*_) = 0 and (4) shows that an estimand of the covariance between two traits unconfounded by *U* is *c*_*g*_. From an MR perspective, by using MCC with ***G*** a set of IVs, in Supplementary Proposition 5 we show that Cov(*Y*_*k*_, *X*_*h*_ | ***G*** = ***g***) is unconfounded by *U*.

Assuming that the individuals for the two phenotypic traits are drawn from the same population with LD matrix between the genetic variants ***V*** = ***G***^⊤^***G***, the sampling distribution of the genetic effects are 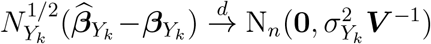 and 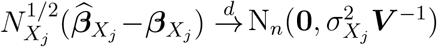, where “*d*” denotes convergence in distribution. Under infinite sample sizes, 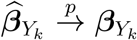 and 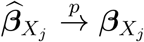, where “*p*” denotes convergence in probability, and an estimator of the genetic covariance between the two traits is

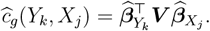

In the finite sample sizes case, the estimates of 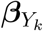 and 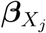 are noised and *ĉ*_*g*_(*Y*_*k*_, *X*_*j*_) is biased [24]

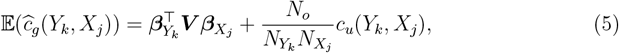

where *N*_*o*_ is the sample size overlap between the two traits. However, even in the scenario of complete overlap, the bias in (5) is negligible if the sample sizes of the two traits are large, as it usually happens in modern GWAS.

The same considerations can made for all phenotypic traits under investigation to reconstruct their joint genetic covariance unconfounded by *U*

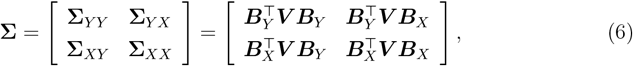

where **Σ**_*XX*_, **Σ**_*YY*_ and **Σ**_*XY*_ are the genetic covariances within the exposures, the outcomes and between them and ***B***_*Y*_ and ***B***_*X*_ are the coefficients of the regressions of the outcomes and the exposures on ***G***, respectively.

### MrDAG model

Assuming that the individuals for two phenotypic traits *Y*_*k*_ and *X*_*j*_ are drawn from the same population with LD matrix ***V***, we have 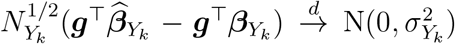 and 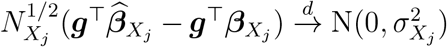, where ***g*** are the observed IVs, 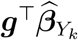 and 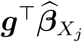 ar e the *k*th and the *j*th genetically predicted values of the outcome and exposure, *i*.*e*., 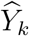 and 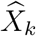, respectively.

The joint distribution of all genetically predicted values of the outcomes and exposures based on the IVs is

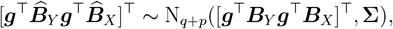

*i*.*e*., for large sample sizes they are normally distributed with mean [ ***g***^⊤^***B***_*Y*_ ***g***^⊤^***B***_*X*_]^⊤^ and covariance matrix **Σ** ∈ 𝒞_𝒟_, the space of the symmetric positive definite covariance matrices Markov with respect to the DAG 𝒟.

If we assume that IVW is performed on the estimated regression coefficients and IVs are independent after pruning or clumping, *i*.*e*., ***V*** = ***I***_*n*_, the MrDAG model becomes

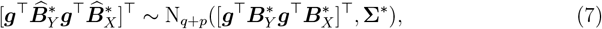

where 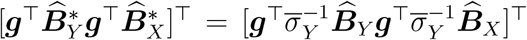 with 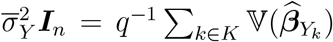 and similarly for 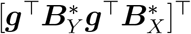. The covariance matrix can be can be partitioned into

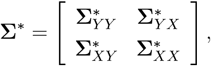

where 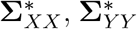 and 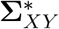 are the genetic covariances within the exposures, the outcomes and between them, and its inverse ([66], Theorem 8.5.11) into

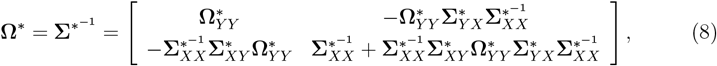

with **Ω**^*^ ∈ 𝒫_𝒟_, the space of the precision matrices Markov with respect to the DAG 𝒟 and 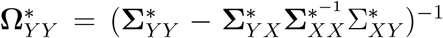. However, since by partial ordering 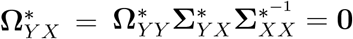, (8) becomes

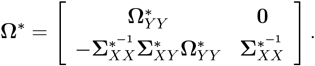

By using **Ω**^*^, Gaussian graphical models [17] can be used to estimate the conditional dependence relationships between the traits in the original (individual-level) data unconfounded by *U* since genetically predicted outcomes and exposures depend only on the selected IVs.

Finally, for a given DAG 𝒟, the estimand of the causal effect under intervention [67] is

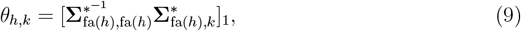

where 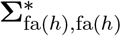 indicates the submatrix of **Σ**^*^ whose rows and columns are fa(*h*), 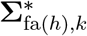 indicates the subvector of **Σ**^*^ whose rows are fa(*h*) and the column correspond to the *k*th outcome, and where the subscript indicates the first element of the vector. By using (6) which is a sufficient statistic for Σ^*^ after IVW and with ***V*** = ***I***_*n*_, (9) becomes

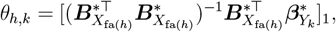

where 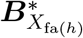 are the IVW coefficients of the regressions of each exposure in ***X***_fa(*h*)_ on ***G*** and 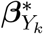 is the IVW coefficient of a regression of *Y*_*k*_ on ***G***. The corresponding estimator coincides with (3).

An important aspect of the MrDAG model is that the genetically predicted values of the outcomes and exposures do not need to be calculated since MrDAG uses as input (6) the sufficient statistic for Σ^*^. The only information that would be required from the original (individual-level) data is the LD matrix ***V***. However, this information is not necessary in MrDAG summary-level MR design since independent genetic variants are considered after pruning or clumping and thus ***V*** = ***I***_*n*_.

### MrDAG algorithm

#### Markov Equivalent Class, Completed Partially DAGs, Essential Graphs and Partially DAGs

The estimation of a DAG from observational data suffers the known problem of identifiability, *i*.*e*., it is not possible to estimate uniquely the underlying true DAG since its conditional independencies can be encoded in several alternative DAGs. This set of DAGs that hold the same conditional independencies is known as Markov Equivalent Class and the best that can be done from observational data is to estimate this class. All DAGs with the same conditional independencies can be represented by a Completed Partially DAG (CPDAG) [68] or Essential Graph (EG) [69]. EGs are Chain Graphs (CGs) whose chain components are decomposable undirected graphs [17]. A CPDAG or EG is a partially directed graph that might contain both directed and undirected edges without directed cycles. Finally, Partially DAG (PDAG) contain both directed and undirected edges and directed cycles might be present.

#### Posterior probability of edge inclusion

Technical details of the algorithm for graphical models exploration that we used to develop MrDAG algorithm are presented in [70]. Briefly, it is based on a Markov chain Monte Carlo (MCMC) algorithm devised to explore the space of EGs whose enumeration is infeasible since their number grows super-exponentially with the number of nodes. The EG 𝒢 is sampled from a proposal distribution which is accepted with a probability given by a Metropolis-Hastings (M-H) ratio defined to guarantee the convergence of the algorithm to the correct posterior distribution. The key ingredient in the M-H ratio is a closed-form expression for the marginal likelihood *m*_*G*_(data). This is based on a non-informative prior coupled with a fractional Bayes factor methodology and compatible priors building procedure. In practice, a specific DAG 𝒟( 𝒢), which belongs to the Markov Equivalent Class whose unique representative chain graph is the EG 𝒢, is proposed and, if accepted, its information stored as an adjacent matrix at each sweep of the MCMC algorithm.

In MrDAG algorithm, we added an acceptance/rejection step to guarantee that 𝒟(𝒢) satisfies the partial ordering that corresponds to the orientation of the edges from the exposures to the outcomes, see Figure 1E. To check the efficiency of this step, we also monitor its acceptance rate. We also included a tempering scheme [71] by considering an annealing parameter *T* in the M-H ratio to facilitate the convergence of the MCMC algorithm to the target distribution and the exploration of regions of high posterior mass. The temperature 1*/T* exponentiates the M-H ratio and its value increases linearly during the burn-in until *T* = 1 at the end of the burn-in.

Sparsity is enforced by assigning a prior to 𝒢 and specifically on 𝒢^*U*^, the skeleton of 𝒢 which contains the same edges of 𝒢 but without orientation

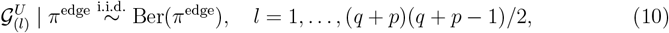

where 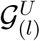 the *l*th element of the vectorized lower triangular part of the adjacency matrix of 𝒢^*U*^ and (*q* + *p*)(*q* + *p* − 1)*/*2 is the maximum number of edges in an EG on *q* + *p* nodes.

The posterior distribution of 𝒢 is

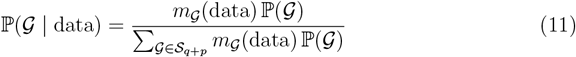

with 𝒮_*q*+*p*_ the set of all EGs with *q* + *p* nodes. The posterior probability of edge inclusion (PPeIs) is defined as

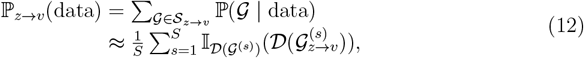

where 𝒮_*z*→*v*_ is the set of EGs containing the directed edge *z* → *v, S* is the number of sweeps after burn-in and 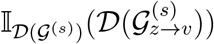 is the indicator function that is equal to one if the specific DAG considered at the *s*th sweep 𝒟( 𝒢^(*s*)^) contains the directed edge *z* → *v* and zero otherwise.

Note that, although MrDAG explores the space of EGs and stores a specific DAG that belongs to the sampled EG, the graphs obtained by thresholding the PPeIs might give rise to a PDAG [70].

#### Bayesian causal effects estimation

Here, we summarise the results reported in [67] that we employed to derive the Bayesian estimation of the causal effects under unmeasured confounders.

Let’s rewrite (7) as

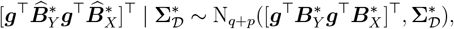

where 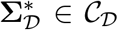, the space of s.p.d. covariance matrices Markov with respect to 𝒟. In the following, for ease of notation, we refer to 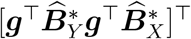 as the “data” and we also drop the subscript 𝒟.

Let 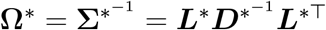 be the modified Cholesky decomposition of the precision **Ω**^*^. The DAG Cholesky parametrization of **Ω**^*^ is given by the node-parameters 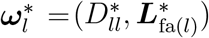, *l* = 1, …, *q* + *p*, with

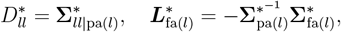

where 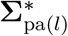 indicates the submatrix of **Σ**^*^ whose rows and columns are pa(*l*).

For a given DAG 𝒟, [67] derive the posterior distribution of 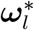, *l* = 1, …, *q* + *p*, in an objective Bayes framework which has the advantage of not depending on priors hyperparameters. In turn, the posterior draws of the Cholesky parameters 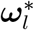 provide posterior draws from 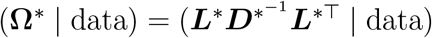 and finally, by using (9), posterior samples of the causal effects between the exposures and the outcomes.

In contrast to frequentist approaches [72] where, for an estimated EG 𝒢, the causal effects are calculated averaging over all (if numerical feasible) or a subset of DAGs within the Markov Equivalent Class 𝒢, here, we also consider the uncertainty related to the estimation of the EGs. Let {𝒢_*v*_, *v* = 1, …, *V* } the set of unique visited EGs by MrDAG. Based on (11), the posterior probability of 𝒢_*v*_ can be approximated by

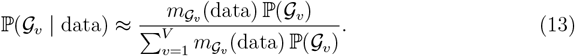

Averaging over the unique visited EGs, the posterior causal effect under intervention in the *h*th exposures on the *k*th outcomes is

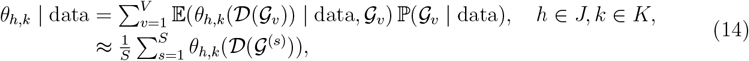

where 𝒢_*v*_ is one of the unique EGs visited during the MCMC, 𝔼 (*θ*_*h,k*_(𝒟( 𝒢_*v*_)) | data, 𝒢_*v*_) is the posterior expectation of the causal effect given 𝒢_*v*_, *i*.*e*., over *all* 𝒟( 𝒢_*v*_), ℙ (𝒢_*v*_ | data) is defined in (13) and *θ*_*h,k*_(*D*(*G*^(*s*)^) is the posterior causal effect conditioned on the recorded DAG at the *s*th sweep.

Finally, by a suitable modification of (14), credible intervals of the causal effects between the exposures and outcomes can be derived.

#### Simulation study

We share several aspects of the simulation study with [2]. It is formulated in a two-sample summary-level MR design, where *N* = 100, 000 independent individuals are simulated, of which *N*_*Y*_ = 50, 000 are used to compute the genetic associations with the exposures and *N*_*X*_ = 50, 000 to compute the genetic associations with the outcomes. Thus, we assume that the quantitative exposures *X*_*j*_, *j* ∈ *J* = {1, …, *p*}, and the quantitative responses *Y*_*k*_, *k* ∈ *K* = {1, …, *q*}, are measured on the same individuals *N*_*X*_ and *N*_*Y*_, respectively, with 100% sample overlap, but independent of each other.

In all simulated scenarios, we consider *p* = 15 exposures, *q* = 5 outcomes and *n* = 100 independent genetic variants as IVs. Genotypes for the *i*th genetic variant and each individual 𝓁 are simulated independently according to a binomial distribution with minor allele frequency (MAF) equal to 0.05, *i*.*e*., 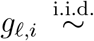 Bin(2, 0.05), 𝓁 ∈ *L* = {1, …, *N*}, *i* ∈ *I* = {1, …, *n*}. The resulting matrix of genotypes ***G*** is split into two equally sized groups, ***G***_*X*_ and ***G***_*Y*_, of dimension *N*_*X*_ *× n* and *N*_*Y*_ *× n*, respectively. Thus, no IVW is needed in the simulation study given that the same MAF at 5% is used to simulate the genotypes.

Overall, the data generation process consists of two stages. In the first stage, the raw data for the exposures ***X*** and the outcomes ***Y*** are simulated. Then, in the second stage, summary-level statistics are obtained as the linear regression coefficients 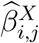 from a univariable linear regression in which the *j*th exposure is regressed on the *i*th genetic variant in sample one and the linear regression coefficients 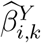 from a univariable linear regression in which the *k*th outcome is regressed on the *i*th genetic variant in sample two.

In the following, we detail each stage and how we simulate the quantities involved. We start with the first stage which is divided into two steps.

- In the first step, the exposures are generated as follows

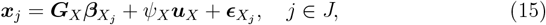

where ***G***_*X*_ and ***u***_*X*_ are the genotypes of the *n* IVs and the values of the confounder *U* measured on the same *N*_*X*_ individuals, respectively, and where 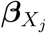 and *ψ*_*X*_ are the corresponding genetic and confounding effects. 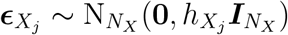 with 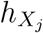 the *j*th diagonal element of the (*p × p*)-dimensional matrix

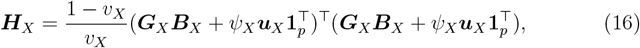

where *v*_*X*_ is the desired level of heritability, or how much variation ***G*** can explain of *X*_*j*_, fixed at 10% for all exposures and in all simulated scenarios. In (16), 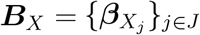 is an (*n × p*)-dimensional matrix of the effects of the genetic variants on the exposures. The confounder *U* is drawn from a multivariate standard Gaussian distribution, *i*.*e*., ***u*** ∼ N_*N*_ (**0, *I***_*N*_) and, then, split into two equally sized vectors ***u***_*X*_ and ***u***_*Y*_ with effect *ψ*_*X*_ impacting all exposures and *ψ*_*Y*_ effecting all outcomes. The effects 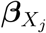 of the *n* genetic variants on the *j*th exposure are drawn following [70]. We randomly generate a topologically ordered DAG among the *p* exposures with a probability of edge inclusion 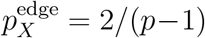 using the function `randomDAG()` in the R package *pcalg* [27]. Thus, the resulting DAG implies the following system of equations [18]

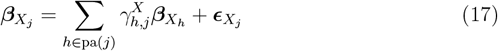

with 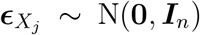. For each *j* ∈ *J*, the effect within the exposures 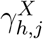 are uniformly chosen in the interval [−1.1*r*_*X*_, −0.9*r*_*X*_]∪[0.9*r*_*X*_, 1.1*r*_*X*_]. This construction procedure for 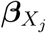 corresponds to the simulated scenario that we call “DAG_*X*_”, *i*.*e*., Directed Acyclic Graph within ***X***, which, in turn, is paired with two different simulated scenarios for the effects 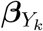 described in the second step (first stage) of the simulation study. We also simulate the effects 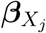 following [2]. Specifically, we simulate 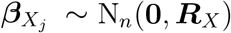, where ***R***_*X*_ is the (*p × p*)-dimensional Toeplitz matrix with 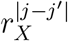 for *j, j*^*′*^ ∈ *J*. The matrix ***R***_*X*_ implies a tridiagonal sparse inverse correlation matrix 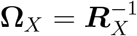. The interpretation of non-zero elements of **Ω**_*X*_ coincides with the effects simulated in (17). We call this second scenario for the effects of the genotypes on the exposures “UG_*X*_”, *i*.*e*., Undirected Graph within ***X***. In both simulated scenarios for ***X***, we use different levels of *r*_*X*_, ranging from independence to a strong dependence, *i*.*e*., *r*_*X*_ = {0, 0.2, 0.4, 0.6, 0.8}, where *r*_*X*_ = 0.6 represents a medium dependence between the genetic associations with the exposures. We use this value in the figures presented in Section ‘Simulation study’.
- In the second step (first stage) of the simulation study, the outcomes are generated on another independent set of *N*_*Y*_ individuals based on the following set of equations

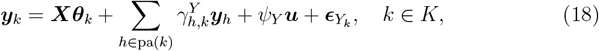

where ***X*** is the (*N*_*X*_ *× p*)-dimensional matrix of exposures simulated using (15), ***θ***_*k*_ = (*θ*_1*k*_, …, *θ*_*pk*_)^⊤^ is *p*-dimensional (sparse) vector the causal effects from the exposures to the *k*th outcome and where *ψ*_*Y*_ is the effect of the confounder *U* on the outcomes. 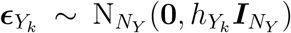 with 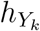 the *k*th diagonal element of the (*q × q*)-dimensional matrix

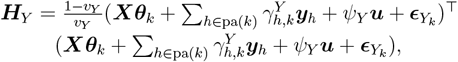

where *v*_*Y*_ is the desired level of the proportion of variance explained, fixed at 25% for all outcomes and in all simulated scenarios.

In (18), the term 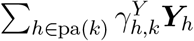 depends on a randomly generated topologically ordered DAG among the *q* outcomes with probability of edge inclusion 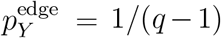. For each *k* ∈ *K*, the effects within the outcomes 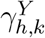 are uniformly drawn in the interval [0.9*m*_*Y*_, 1.1*m*_*Y*_ ]. In analogy with the first step, we call this scenario “DAG_*Y*_ ”, *i*.*e*., Directed Acyclic Graph within ***Y***.

We also simulate a simplified scenario where

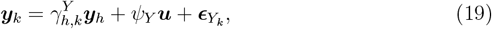

*i*.*e*., a randomly selected outcome *k* is completed mediated by another randomly selected response chosen between the remaining ones. We call this scenario “Med_*Y*_ ”, *i*.*e*., complete mediation of an outcome, since in the previous scenario “DAG_*Y*_ ” partial mediations [73] are likely simulated, while here we exclude this case. In this second simulated scenario for the outcomes, the matrix ***H***_*Y*_ is calculated according to (19). Moreover, we use different levels of *m*_*Y*_, ranging from small to a strong level of (partial or complete) mediation, *i*.*e*., *m*_*Y*_ = {0.25, 0.50, 0.75, 1, 1.5, 2}, where *m*_*Y*_ = 1 represents a medium (partial or complete) mediation effect. We use this value in the figures presented in Section ‘Simulation study’.

Finally, the causal effects ***θ***_*k*_ are drawn independently from a multivariate Gaussian distribution, *i*.*e*., ***θ***_*k*_ ∼ N_*p*_(**0, *I***_*p*_).

In both simulated scenarios for ***Y***, we consider a (*q × p*)-dimensional sparse matrix of causal effects **Θ** = {***θ***_*k*_}_*k*∈*K*_, where 30 cells of the matrix are non-zero and where several exposures are either *shared* or *distinct* for the outcomes. Specifically, we select at random the same proportion of cells in the matrix **Θ** and assign them the simulated values, while the other cells are set to zero.

After the first stage, four scenarios are created by combining the simulations for ***X*** and ***Y*** : (i) “UndG_*X*_-Med_*Y*_ ”, *i*.*e*., undirected graph within ***X*** and complete mediation of an outcome in ***Y*** ; (ii) DAG_*X*_-Med_*Y*_, *i*.*e*., topologically ordered DAG within ***X*** and complete mediation of a response within ***Y*** ; (iii) UndG_*X*_-DAG_*Y*_, *i*.*e*., undirected graph within ***X*** and topologically ordered DAG within ***Y*** ; (iv) DAG_*X*_-DAG_*Y*_, topologically ordered DAGs within ***X*** and ***Y***. In (ii) and (iv) the overall DAGs, obtained by combining different simulation patterns for ***X*** and ***Y***, are fully oriented while in (i) and (iii) they are partially oriented.

After creating the data at the individual level, in the second stage, we compute the summary-level statistics from the two independent groups of individuals. The input data for the simulation study are the summary-level statistics 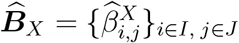, an (*n × p*)-dimensional matrix, and 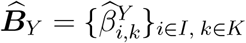, an (*n×q*)-dimensional matrix, derived from a univariable linear regression model, where each genetic variant *G*_*i*_ is regressed against each exposure *X*_*j*_ and each outcome *Y*_*k*_ at-a-time.

#### Real data application: Pre-processing and data preparation

The first step of the data processing merges the summary-level data (beta regression coefficients, their standard errors and associated *p*-values) of all exposures by their unique “rs” identifier and aligns the effect direction of the genetic associations with each exposure according to the same effect allele. As IVs, we select the genetic variants which are associated with any of the exposures at genome-wide significance (minimum *p*-value *<* 5*×*10^−8^ across all exposures). Next, we merge the genetic variants selected as IVs with the outcome data by their unique “rs” identifier and align the effect direction of the genetic associations with each outcome according to the same effect allele. Finally, we clump the genetic variants to be independent at *r*^2^ *<* 0.01 using a European reference panel [74]. This results in *n* = 708 independent genetic variants selected as IVs. See Supplementary Table 1 for the description of the summary-level statistics, the data sources, the number of non-unique IVs which were genome-wide significant for each exposure along with the contribution (%) of each exposure on the selected IVs.

Finally, we perform reverse causation using the same traits with mental health phenotypes as exposures and lifestyle and behavioural traits as outcomes. We apply the same procedure described above resulting in 470 IVs. See Supplementary Table 1 for details regarding the number of non-unique IVs which were genome-wide significant for each exposure along with the contribution (%) of each exposure on the selected IVs.

## Supporting information

Technical details along with figures and tables to support the results of the simulation study and the real data analysis

## Data availability

Data sources are presented in Supplementary Information with associated URL links. Social Science Genetic Association Consortium (SSGAC) summary-level statistics are available through a standard registration procedure (https://thessgac.com/register/).

## Code availability

The Mendelian randomization with Directed Acyclic Graph learning R package `MrDAG`is freely available on https://github.com/lb664/MrDAG/. It includes the data of the real data examples and how to run the algorithm. Post-processing routines to estimate the posterior causal effects presented in the manuscript are also included along with the Posterior Probability of Edge Inclusion.

## Acknowledgements

The authors are thankful to Federico Castelletti and Guido Consonni for their insightful comments and suggestions and to the participants of the 1st Danish International Conference on Personalised Medicine, Aarhus, DK, whose remarks regarding preliminary results of the real data application led to its substantial improvement.

## Author contributions

Conceptualization: L.B., V.Z., D.G.; Methodology: L.B, V.Z.; Formal Analysis: V.Z., L.B.; Resources: (all collaborators); Data curation: V.Z., D.G.; Writing – original draft: V.Z., L.B., T.C., G.D., N.C.; Visualization: V.Z., L.B.; Supervision: L.B., D.G.

## Additional information

Supplementary Information includes technical details regarding the identifiability and the estimation of causal effects for a given DAG, the consistency of the effects of the regressions of the exposures and the outcomes on the selected genetic variants along with figures and tables to support the results of the simulation study and the real data analysis.

