## Supplementary material for "Mendelian randomization for multiple exposures and outcomes with Bayesian Directed Acyclic Graphs exploration and causal effects estimation": Technical details along with figures and tables to support the results of the simulation study and the real data analysis

### Contents

1 Supplementary Text 3

2 Supplementary Tables 17

3 Supplementary Figures 19

### 1 Supplementary Text

In the following, we assume that the conditional expectation of the outcome $Y_k$, $k \in K$, and the conditional expectation of the exposure $X_j$, $j \in J$, are linear with regard to their DAG parents without interactions and all dependencies only affect the mean. These assumptions lead to

$$\mathbb{E}(Y_k \mid \boldsymbol{Y}_{\mathrm{pa}(k)} = \boldsymbol{y}_{\mathrm{pa}(k)}, \boldsymbol{X}_{\mathrm{pa}(k)} = \boldsymbol{x}_{\mathrm{pa}(k)}, U = u) = \sum_{j \in \mathrm{pa}(k)} \theta_{j,k} x_j + \sum_{h \in \mathrm{pa}(k)} \gamma^Y_{h,k} y_h + \psi_Y u,$$
$$\mathbb{E}(X_j \mid \boldsymbol{X}_{\mathrm{pa}(j)} = \boldsymbol{x}_{\mathrm{pa}(j)}, \boldsymbol{G} = \boldsymbol{g}, U = u) = \sum_{i \in I} \beta^X_{i,j} g_i + \sum_{h \in \mathrm{pa}(j)} \gamma^X_{h,j} x_h + \psi_X u,$$

where $\mathrm{pa}_{\mathcal{D}}(v)$ denotes the parent set of the node $v$ in the DAG $\mathcal{D}$ (for ease of notation, we removed the subscript $\mathcal{D}$), $\gamma^Y_{h,k}$ is the effect of the $h$th outcomes on the $k$th outcome, $h \neq k$, $\theta_{j,k}$, is the causal effect of the $j$th exposure on the $k$th outcome and $\gamma^X_{h,j}$ is the effect of the $h$th exposure on the $j$th exposure, $h \neq j$.

In addition, we assume that the expectation of $Y_k$ conditionally on an intervention on $X_h$, $h \in \mathrm{pa}(k)$, is

$$\mathbb{E}(Y_k \mid \mathrm{do}(X_h = \widetilde{x}_h), \boldsymbol{X}_{j \in \mathrm{pa}(h)} = \boldsymbol{x}_{j \in \mathrm{pa}(h)}, U = u) = \theta_{h,k} \widetilde{x}_h + \sum_{j \in \mathrm{pa}(h)} \theta_{j,k} x_j + \psi_Y u. \quad (1)$$

#### Regressions of outcomes and exposures on $\boldsymbol{G}$

The proof that, for a given DAG $\mathcal{D}$, the effects of $\mathbb{E}(Y_k \mid \boldsymbol{G} = \boldsymbol{g})$, $k \in K$, and $\mathbb{E}(X_j \mid \boldsymbol{G} = \boldsymbol{g})$, $j \in J$, can be obtained without adjustment on $U$ because $\boldsymbol{G}$ is randomly assigned at conception [1], as shown for standard MR [2], is provided in the following proposition. The proof depends on the multivariate "core conditions" (MCC) for valid IVs presented in the main text Methods, the Markov properties (MP) of the DAG and the different expectations being additive in the conditioning variables. For $\mathbb{E}(Y_k \mid \boldsymbol{G} = \boldsymbol{g})$, we compute the conditional expectation of $Y_k$ given $(\boldsymbol{Y}_{\mathrm{pa}(k)}, \boldsymbol{X}_{\mathrm{pa}(k)}, U)$, then (for each outcome in $\boldsymbol{Y}_{\mathrm{pa}(k)}$ we proceed similarly) integrate out each exposure in $\boldsymbol{X}_{\mathrm{pa}(k)}$ with respect to its conditional distribution given $(\boldsymbol{G} = \boldsymbol{g}, U)$ until there are no exposures that have other exposures as parents and, finally, integrate out the unmeasured confounder $U$ with regard to its marginal distribution. We use the same strategy to compute the conditional expectation of $X_j$ given $\boldsymbol{G}$.

**Proposition 1.** For a given DAG $\mathcal{D}$ that represents the conditional dependencies between $\boldsymbol{Y}$, $\boldsymbol{X}$, $\boldsymbol{G}$ and $U$, $\mathbb{E}(Y_k \mid \boldsymbol{G} = \boldsymbol{g})$, $\forall k \in K$, and $\mathbb{E}(X_j \mid \boldsymbol{G} = \boldsymbol{g})$, $\forall j \in J$, are unconfounded by $U$.

*Proof.* Let's assume that the multivariate extension of the "core conditions" for a valid instrumental variable hold:

(IV1) $G_i \perp\!\!\!\perp U$, $\forall i \in I$, *i.e.*, $G_i$ must be independent of $U$;

(IV2) $G_i \not\perp\!\!\!\perp X_j \mid \boldsymbol{X}_{\setminus j}$, $\forall i \in I$ and $\forall j \in J$, *i.e.*, $G_i$ must *not* be independent of $X_j$ conditionally on $\boldsymbol{X}_{\setminus j}$;

(IV3) $G_i \perp\!\!\!\perp Y_k \mid (\boldsymbol{X}, U)$, $\forall i \in I$ and $\forall k \in K$, *i.e.*, $G_i$ must be independent of $Y_k$ conditionally on $\boldsymbol{X}$ and $U$.

For a given DAG, let's assume that, among the outcomes and exposures, the parents of $Y_k$ are $\boldsymbol{Y}_{\mathrm{pa}(k)}$ and $\boldsymbol{X}_{\mathrm{pa}(k)}$, $\boldsymbol{Y}_{\mathrm{pa}(k)}$ do not have any outcomes among their parents and $\boldsymbol{X}_{\mathrm{pa}(k)}$ have parents among the exposures. We denote the parents of $\boldsymbol{X}_{\mathrm{pa}(k)}$ as $\boldsymbol{X}_{\mathrm{pa}(\mathrm{pa}(k))}$ which, in turn, do not have any exposures among their parents. We can write the conditional expectation of $Y_k$ given $\boldsymbol{G}$ as

$$
\begin{aligned}
\mathbb{E}(Y_k \mid \boldsymbol{G} = \boldsymbol{g}) &= \mathbb{E}_{U|G=g}\, \mathbb{E}_{Y_{\mathrm{pa}(k)}, X_{\mathrm{pa}(k)}, X_{\mathrm{pa}(\mathrm{pa}(k))}|G=g,U} \\
&\quad \mathbb{E}(Y_k \mid \boldsymbol{Y}_{\mathrm{pa}(k)}, \boldsymbol{X}_{\mathrm{pa}(k)}, \boldsymbol{X}_{\mathrm{pa}(\mathrm{pa}(k))}, \boldsymbol{G} = \boldsymbol{g}, U) \\
&= \mathbb{E}_{U|G=g}\, \mathbb{E}_{X_{\mathrm{pa}(\mathrm{pa}(k))}|G=g,U}\, \mathbb{E}_{X_{\mathrm{pa}(k)}|X_{\mathrm{pa}(\mathrm{pa}(k))},G=g,U}\, \mathbb{E}_{Y_{\mathrm{pa}(k)}|X_{\mathrm{pa}(k)},X_{\mathrm{pa}(\mathrm{pa}(k))},G=g,U} \\
&\quad \mathbb{E}(Y_k \mid \boldsymbol{Y}_{\mathrm{pa}(k)}, \boldsymbol{X}_{\mathrm{pa}(k)}, \boldsymbol{X}_{\mathrm{pa}(\mathrm{pa}(k))}, \boldsymbol{G} = \boldsymbol{g}, U) \\
&= \mathbb{E}_{U|G=g}\, \mathbb{E}_{X_{\mathrm{pa}(\mathrm{pa}(k))}|G=g,U}\, \mathbb{E}_{X_{\mathrm{pa}(k)}|X_{\mathrm{pa}(\mathrm{pa}(k))},G=g,U} \\
&\quad \mathbb{E}_{Y_{\mathrm{pa}(k)}|X_{\mathrm{pa}(k)},X_{\mathrm{pa}(\mathrm{pa}(k))},U}\, \mathbb{E}(Y_k \mid \boldsymbol{Y}_{\mathrm{pa}(k)}, \boldsymbol{X}_{\mathrm{pa}(k)}, \boldsymbol{G} = \boldsymbol{g}, U) \text{ since MP} \\
&= \mathbb{E}_{U}\, \mathbb{E}_{X_{\mathrm{pa}(\mathrm{pa}(k))}|G=g,U}\, \mathbb{E}_{Y_{\mathrm{pa}(k)}|X_{\mathrm{pa}(k)},X_{\mathrm{pa}(\mathrm{pa}(k))},U} \\
&\quad \mathbb{E}_{X_{\mathrm{pa}(k)}|X_{\mathrm{pa}(\mathrm{pa}(k))},G=g,U}\, \mathbb{E}(Y_k \mid \boldsymbol{Y}_{\mathrm{pa}(k)}, \boldsymbol{X}_{\mathrm{pa}(k)}, U) \text{ since IV1 and IV3} \\
&= \mathbb{E}_{U}\, \mathbb{E}_{X_{\mathrm{pa}(\mathrm{pa}(k))}|G=g,U}\, \mathbb{E}_{Y_{\mathrm{pa}(k)}|X_{\mathrm{pa}(k)},X_{\mathrm{pa}(\mathrm{pa}(k))},U} \\
&\quad \mathbb{E}_{X_{\mathrm{pa}(k)}|X_{\mathrm{pa}(\mathrm{pa}(k))},G=g,U}\Big(\sum_{j\in\mathrm{pa}(k)} \theta_{j,k} X_j + \sum_{h\in\mathrm{pa}(k)} \gamma^{Y}_{h,k} Y_h + \psi_Y U\Big) \\
&= \sum_{j\in\mathrm{pa}(k)} \theta_{j,k} \underbrace{\mathbb{E}_U\, \mathbb{E}_{X_{\mathrm{pa}(j)}|G=g,U}\, \mathbb{E}_{X_j|X_{\mathrm{pa}(j)}G=g,U}\, X_j}_{T_1} + \\
&\quad \sum_{h\in\mathrm{pa}(k)} \gamma^{Y}_{h,k} \underbrace{\mathbb{E}_U\, \mathbb{E}_{X_{\mathrm{pa}(h)}|G=g,U}\, \mathbb{E}_{Y_h|X_{\mathrm{pa}(h)},U}\, Y_h}_{T_2} + \psi_Y\, \mathbb{E}\, U.
\end{aligned}
\tag{2}
$$

Under the assumptions regarding the parents of $\boldsymbol{Y}_{\mathrm{pa}(k)}$ and $\boldsymbol{X}_{\mathrm{pa}(\mathrm{pa}(k))}$, we have

$$
\begin{aligned}
T_1 &= \mathbb{E}_U\, \mathbb{E}_{X_{\mathrm{pa}(j)}|G=g,U}\Big(\sum_{s\in\mathrm{pa}(j)} \gamma^{X}_{s,j} X_s + \boldsymbol{g}^\top \boldsymbol{\beta}_{X_j} + \psi_X U\Big) \\
&= \boldsymbol{g}^\top \boldsymbol{\beta}_{X_j} + \sum_{s\in\mathrm{pa}(j)} \gamma^{X}_{s,j}\, \mathbb{E}_U\, \mathbb{E}_{X_s|G=g,U}\, X_s + \psi_X\, \mathbb{E}\, U \\
&= \boldsymbol{g}^\top \boldsymbol{\beta}_{X_j} + \sum_{s\in\mathrm{pa}(j)} \gamma^{X}_{s,j}(\boldsymbol{g}^\top \boldsymbol{\beta}_{X_s} + \psi_X\, \mathbb{E}\, U) + \psi_X\, \mathbb{E}\, U \\
&= \boldsymbol{g}^\top \boldsymbol{\beta}_{X_j} + \sum_{s\in\mathrm{pa}(j)} \gamma^{X}_{s,j} \boldsymbol{g}^\top \boldsymbol{\beta}_{X_s} + \{(1 + \sum_{s\in\mathrm{pa}(j)} \gamma^{X}_{s,j})\}\psi_X\, \mathbb{E}\, U
\end{aligned}
\tag{3}
$$

and

$$
\begin{aligned}
T_2 &= \mathbb{E}_U\, \mathbb{E}_{X_{\mathrm{pa}(h)}|G=g,U}\Big(\sum_{r\in\mathrm{pa}(h)} \theta_{r,h} X_r + \psi_Y U\Big) \\
&= \sum_{r\in\mathrm{pa}(h)} \theta_{r,h}\, \mathbb{E}_U\, \mathbb{E}_{X_r|G=g,U}\, X_r + \psi_Y\, \mathbb{E}\, U \\
&= \sum_{r\in\mathrm{pa}(h)} \theta_{r,h}(\boldsymbol{g}^\top \boldsymbol{\beta}_{X_r} + \psi_X\, \mathbb{E}\, U) + \psi_Y\, \mathbb{E}\, U \\
&= \sum_{r\in\mathrm{pa}(h)} \theta_{r,h} \boldsymbol{g}^\top \boldsymbol{\beta}_{X_r} + \Big(\sum_{r\in\mathrm{pa}(h)} \theta_{r,h} \psi_X + \psi_Y\Big)\, \mathbb{E}\, U.
\end{aligned}
\tag{4}
$$

Combining (2), (3) and (4), we obtain

$$\begin{aligned}\mathbb{E}(Y_k \mid \boldsymbol{G}=\boldsymbol{g}) = \alpha_{Y_k} + \sum_{j\in\mathrm{pa}(k)} \theta_{j,k}\boldsymbol{g}^\top\boldsymbol{\beta}_{X_j} + \sum_{j\in\mathrm{pa}(k)} \theta_{j,k} \sum_{s\in\mathrm{pa}(j)} \gamma^X_{s,j}\boldsymbol{g}^\top\boldsymbol{\beta}_{X_s} + \\ \sum_{h\in\mathrm{pa}(k)} \gamma^Y_{h,k} \sum_{r\in\mathrm{pa}(h)} \theta_{r,h}\boldsymbol{g}^\top\boldsymbol{\beta}_{X_r},\end{aligned} \tag{5}$$

where $\sum_{j\in\mathrm{pa}(k)} \theta_{j,k}\boldsymbol{g}^\top\boldsymbol{\beta}_{X_j}$, $\sum_{j\in\mathrm{pa}(k)} \theta_{j,k} \sum_{s\in\mathrm{pa}(j)} \gamma^X_{s,j}\boldsymbol{g}^\top\boldsymbol{\beta}_{X_s}$ and $\sum_{h\in\mathrm{pa}(k)} \gamma^Y_{h,k} \sum_{r\in\mathrm{pa}(h)} \theta_{r,h}\boldsymbol{g}^\top\boldsymbol{\beta}_{X_r}$ are the genetic components of $Y_k$ determined by the genetic components of the exposures among its parents and mediated by other exposures and outcomes.

Similarly, assuming that $\boldsymbol{X}_{\mathrm{pa}(\mathrm{pa}(j))}$ do have parents among the exposures, we have

$$\begin{aligned}\mathbb{E}(X_j \mid \boldsymbol{G}=\boldsymbol{g}) &= \mathbb{E}_{U|G=g}\,\mathbb{E}_{X_{\mathrm{pa}(\mathrm{pa}(j))}|G=g,U}\,\mathbb{E}_{X_{\mathrm{pa}(j)}|X_{\mathrm{pa}(\mathrm{pa}(j))},G=g,U} \\ &\quad \mathbb{E}(X_j \mid \boldsymbol{X}_{\mathrm{pa}(j)}, \boldsymbol{X}_{\mathrm{pa}(\mathrm{pa}(j))}, \boldsymbol{G}=\boldsymbol{g}, U) \\ &= \mathbb{E}_{U|G=g}\,\mathbb{E}_{X_{\mathrm{pa}(\mathrm{pa}(j))}|G=g,U}\,\mathbb{E}_{X_{\mathrm{pa}(j)}|X_{\mathrm{pa}(\mathrm{pa}(j))},G=g,U} \\ &\quad \mathbb{E}(X_j \mid \boldsymbol{X}_{\mathrm{pa}(j)}, \boldsymbol{G}=\boldsymbol{g}, U) \text{ since MP} \\ &= \mathbb{E}_U\,\mathbb{E}_{X_{\mathrm{pa}(\mathrm{pa}(j))}|G=g,U}\,\mathbb{E}_{X_{\mathrm{pa}(j)}|X_{\mathrm{pa}(\mathrm{pa}(j))},G=g,U}(\boldsymbol{g}^\top\boldsymbol{\beta}_{X_j} + \sum_{r\in\mathrm{pa}(j)} \gamma^X_{r,j}X_r + \psi_X U) \\ &= \boldsymbol{g}^\top\boldsymbol{\beta}_{X_j} + \sum_{r\in\mathrm{pa}(j)} \gamma^X_{r,j}\,\mathbb{E}_U\,\mathbb{E}_{X_{\mathrm{pa}(r)}|G=g,U}\,\mathbb{E}_{X_r|X_{\mathrm{pa}(r)},G=g,U} X_r + \psi_X\,\mathbb{E}\,U \\ &\quad \text{since IV1} \\ &= \boldsymbol{g}^\top\boldsymbol{\beta}_{X_j} + \sum_{r\in\mathrm{pa}(j)} \gamma^X_{r,j}\,\mathbb{E}_U\,\mathbb{E}_{X_{\mathrm{pa}(r)}|G=g,U}(\boldsymbol{g}^\top\boldsymbol{\beta}_{X_r} + \sum_{s\in\mathrm{pa}(r)} \gamma^X_{s,r}X_s + \psi_X U) + \\ &\quad \psi_X\,\mathbb{E}\,U \\ &= \boldsymbol{g}^\top\boldsymbol{\beta}_{X_j} + \sum_{r\in\mathrm{pa}(j)} \gamma^X_{r,j}(\boldsymbol{g}^\top\boldsymbol{\beta}_{X_r} + \sum_{s\in\mathrm{pa}(r)} \gamma^X_{s,r}\,\mathbb{E}_U\,\mathbb{E}_{X_s|G=g,U} X_s + \psi_X U) + \\ &\quad \psi_X\,\mathbb{E}\,U \\ &= \boldsymbol{g}^\top\boldsymbol{\beta}_{X_j} + \sum_{r\in\mathrm{pa}(j)} \gamma^X_{r,j}\{\boldsymbol{g}^\top\boldsymbol{\beta}_{X_r} + \sum_{s\in\mathrm{pa}(r)} \gamma^X_{s,r}(\boldsymbol{g}^\top\boldsymbol{\beta}_{X_s} + \psi_X\,\mathbb{E}\,U) + \psi_X\,\mathbb{E}\,U\} + \\ &\quad \psi_X\,\mathbb{E}\,U \\ &= \alpha_{X_j} + \boldsymbol{g}^\top\boldsymbol{\beta}_{X_j} + \sum_{r\in\mathrm{pa}(j)} \gamma^X_{r,j}\boldsymbol{g}^\top\boldsymbol{\beta}_{X_r} + \sum_{r\in\mathrm{pa}(j)} \gamma^X_{r,j} \sum_{s\in\mathrm{pa}(r)} \gamma^X_{s,r}\boldsymbol{g}^\top\boldsymbol{\beta}_{X_s},\end{aligned} \tag{6}$$

where $\boldsymbol{g}^\top\boldsymbol{\beta}_{X_j}$, $\sum_{r\in\mathrm{pa}(j)} \gamma^X_{r,j}\boldsymbol{g}^\top\boldsymbol{\beta}_{X_r}$ and $\sum_{r\in\mathrm{pa}(j)} \gamma^X_{r,j} \sum_{s\in\mathrm{pa}(r)} \gamma^X_{s,r}\boldsymbol{g}^\top\boldsymbol{\beta}_{X_s}$ are the direct and mediated genetic components of $X_j$ which, in turn, are function of the genetic components of other exposures. ∎

**Remark 1.** In (2) and (6), we made some assumptions regarding the dependency structure of the parents of $\boldsymbol{Y}_{\mathrm{pa}(k)}$ and $\boldsymbol{X}_{\mathrm{pa}(\mathrm{pa}(k))}$, and the parents of $\boldsymbol{X}_{\mathrm{pa}(\mathrm{pa}(j))}$, respectively. The proof does not change if further dependency relations are considered among the outcomes and the exposures given that the expectations are additive in the conditioning variables and each exposure can be integrated out with regard to its conditional distribution. Under these assumptions, regardless of the dependency structure associated with a given DAG, it is possible to integrate out $U$ with regard to its marginal distribution.

**Remark 2.** In (5), the regression coefficient of a regression of $Y_k$ on $\boldsymbol{G}$ is "consistent" for the causal parameters of interest as well as the mediation parameters within the outcomes and the exposures. In (6), the regression coefficient of a regression of $X_j$ on $\boldsymbol{G}$ yields a consistent estimate of the mediation parameters within the exposures.

#### Identification and estimation of the causal effects

In this section, we first specify what the causal parameters of interest are, and second determine how they relate to the parameters of a regression of $Y_k$ on $\boldsymbol{G}$ (2) and a regression of $X_j$ on $\boldsymbol{G}$ (6).

To prove that

$$\theta_{h,k} = \frac{\partial}{\partial x_h} \mathbb{E}(Y_k \mid \mathrm{do}(X_h = x_h))\bigg|_{x_h=\widetilde{x}_h}, \tag{7}$$

we assume general linear models for the dependencies among the variables $\boldsymbol{Y}$, $\boldsymbol{X}$, $\boldsymbol{G}$ and $U$ and compute the conditional expectation of $Y_k$ given $(\boldsymbol{X}_{\mathrm{pa}(h)}, U)$. Then, we remove the conditioning on $\mathrm{do}(X_h = x_h)$ since there cannot be dependence between any random variables and an intervention, integrate out each exposure in $\boldsymbol{X}_{\mathrm{pa}(h)}$ with respect to its conditional distribution and, finally, integrate out $\boldsymbol{G}$ and $U$ with regard to their marginal distributions.

**Proposition 2.** For a given DAG $\mathcal{D}$ that represents the conditional dependencies between $\boldsymbol{Y}$, $\boldsymbol{X}$, $\boldsymbol{G}$ and $U$, the causal effect of an intervention in $X_h = \widetilde{x}_h$ on $Y_k$ is $\theta_{h,k}$.

*Proof.* Let's assume MCC and model (1) hold. To show (7), it is sufficient to prove that

$$\begin{aligned}
\mathbb{E}(Y_k \mid \mathrm{do}(X_h = x_h)) &= \mathbb{E}_{X_{\mathrm{pa}(h)},U|\mathrm{do}(X_h=x_h)} \mathbb{E}(Y_k \mid \mathrm{do}(X_h = x_h), \boldsymbol{X}_{\mathrm{pa}(h)}, U) \\
&= \mathbb{E}_{U|\mathrm{do}(X_h=x_h)} \mathbb{E}_{X_{\mathrm{pa}(h)}|\mathrm{do}(X_h=x_h),U} \mathbb{E}(Y_k \mid \mathrm{do}(X_h = x_h), \boldsymbol{X}_{\mathrm{pa}(h)}, U) \\
&= \mathbb{E}_U \mathbb{E}_{X_{\mathrm{pa}(h)}|U} \mathbb{E}(Y_k \mid \mathrm{do}(X_h = x_h), \boldsymbol{X}_{\mathrm{pa}(h)}, U) \\
&= \mathbb{E}_U \mathbb{E}_{X_{\mathrm{pa}(h)}|U} (\theta_{h,k} x_h + \textstyle\sum_{j\in \mathrm{pa}(h)} \theta_{j,k} X_j + \psi_Y U) \\
&= \theta_{h,k} x_h + \mathbb{E}_U (\textstyle\sum_{j\in \mathrm{pa}(h)} \theta_{j,k} \mathbb{E}_{X_j|U} X_j + \psi_Y U) \\
&= \theta_{h,k} x_h + \mathbb{E}_U (\textstyle\sum_{j\in \mathrm{pa}(h)} \theta_{j,k} \mathbb{E}_G \mathbb{E}_{X_j|G,U} X_j + \psi_Y U) \text{ since IV1} \\
&= \theta_{h,k} x_h + \mathbb{E}_U (\textstyle\sum_{j\in \mathrm{pa}(h)} \theta_{j,k} \mathbb{E}_G (\sum\limits_{i\in I} G_i \beta^X_{i,j} + \psi_X U) + \psi_Y U) \\
&= \theta_{h,k} x_h + \textstyle\sum_{j\in \mathrm{pa}(h)} \theta_{j,k} (\sum\limits_{i\in I} \mathbb{E}\, G_i \beta^X_{i,j} + \psi_X \mathbb{E}\, U) + \psi_Y \mathbb{E}\, U.
\end{aligned}$$

■

The next proposition shows that the causal parameter of interest is related to the regression parameters of a regression of $Y_k$ on $\boldsymbol{G}$ and a regression of $X_j$ on $\boldsymbol{G}$.

**Proposition 3.** The estimand of the causal parameter of interest $\theta_{h,k}$ is the solution of the linear least squares (LLS) regression of $\boldsymbol{\beta}_{Y_k}$ on $\boldsymbol{B}_{X_{\mathrm{fa}(h)}} = \{\boldsymbol{\beta}_{X_j}\}_{j\in \mathrm{pa}(h)}$.

*Proof.* Define $\overline{G}_i = G_i - \mathbb{E}(G_i)$, $\forall i \in I$, so that $\mathrm{Cov}(Y_k, G_i) = \mathbb{E}(Y_k \overline{G}_i)$. With MCC and model (1), we have

$$
\begin{aligned}
\mathbb{E}(Y_k\overline{G}_i) &= \mathbb{E}_U\,\mathbb{E}_{G_i|U}\,\mathbb{E}_{X_{\mathrm{pa}(h)}|G_i,U}\,\mathbb{E}_{X_h|X_{\mathrm{pa}(h)},G_i,U}\,\mathbb{E}(Y_k\overline{G}_i \mid X_h, \boldsymbol{X}_{\mathrm{pa}(h)}, \overline{G}_i, U)\\
&= \mathbb{E}_U\,\mathbb{E}_{G_i}\,\mathbb{E}_{X_{\mathrm{pa}(h)}|G_i,U}\\
&\quad \mathbb{E}_{X_h|X_{\mathrm{pa}(h)},G_i,U}(\{\theta_{h,k}X_h + \textstyle\sum_{j\in\mathrm{pa}(h)}\theta_{j,k}X_j + \psi_Y U\}\overline{G}_i) \text{ since IV1}\\
&= \theta_{h,k}\,\mathbb{E}_{G_i}\,\mathbb{E}_{X_h|G_i}(X_h\overline{G}_i) + \textstyle\sum_{j\in\mathrm{pa}(h)}\theta_{j,k}\,\mathbb{E}_{G_i}\,\mathbb{E}_{X_j|G_i}(X_j\overline{G}_i) + \psi_Y\,\mathbb{E}_U\,\mathbb{E}_{G_i}(U\overline{G}_i)\\
&= \theta_{h,k}\,\mathbb{E}(X_h\overline{G}_i) + \textstyle\sum_{j\in\mathrm{pa}(h)}\theta_{j,k}\,\mathbb{E}(X_j\overline{G}_i) + \psi_Y\,\mathbb{E}\,U\,\mathbb{E}\,\overline{G}_i \text{ since IV1}\\
&= \theta_{h,k}\,\mathbb{E}(X_h\overline{G}_i) + \textstyle\sum_{j\in\mathrm{pa}(h)}\theta_{j,k}\,\mathbb{E}(X_j\overline{G}_i).
\end{aligned}
$$

Dividing both sides by $\mathbb{E}\,\overline{G}_i^2$, we obtain

$$
\begin{aligned}
\frac{\mathbb{E}(Y_k\overline{G}_i)}{\mathbb{E}\,\overline{G}_i^2} &= \theta_{h,k}\frac{\mathbb{E}(X_h\overline{G}_i)}{\mathbb{E}\,\overline{G}_i^2} + \sum_{j\in\mathrm{pa}(h)}\theta_{j,k}\frac{\mathbb{E}(X_j\overline{G}_i)}{\mathbb{E}\,\overline{G}_i^2}\\
\beta^Y_{i,k} &= \theta_{h,k}\beta^X_{i,h} + \sum_{j\in\mathrm{pa}(h)}\theta_{j,k}\beta^X_{i,j}
\end{aligned}
$$

and in matrix form for all $i \in I$

$$
\begin{aligned}
\boldsymbol{\beta}_{Y_k} &= \boldsymbol{\beta}_{X_h}\theta_{h,k} + \textstyle\sum_{j\in\mathrm{pa}(h)}\boldsymbol{\beta}_{X_j}\theta_{j,k}\\
&= \boldsymbol{B}_{X_{\mathrm{fa}(h)}}\boldsymbol{\theta}_{\mathrm{fa}(h),k},
\end{aligned}
$$

where $\mathrm{fa}(v) = v \cup \mathrm{pa}(v)$ is the family of $v$. Assuming that $n > p$, we obtain

$$
\theta_{h,k} = [(\boldsymbol{B}^\top_{X_{\mathrm{fa}(h)}}\boldsymbol{B}_{X_{\mathrm{fa}(h)}})^{-1}\boldsymbol{B}^\top_{X_{\mathrm{fa}(h)}}\boldsymbol{\beta}_{Y_k}]_1, \tag{8}
$$

where the subscript indicates the first element of the solution of the LLS regression which is well defined because $\boldsymbol{B}_{X_{\mathrm{fa}(h)}} \neq \boldsymbol{0}$ since IV2. Finally, the estimator of the causal parameter of interest is

$$
\widehat{\theta}_{h,k} = [(\widehat{\boldsymbol{B}}^\top_{X_{\mathrm{fa}(h)}}\widehat{\boldsymbol{B}}_{X_{\mathrm{fa}(h)}})^{-1}\widehat{\boldsymbol{B}}^\top_{X_{\mathrm{fa}(h)}}\widehat{\boldsymbol{\beta}}_{Y_k}]_1. \tag{9}
$$

■

**Remark 3.** The IVW estimator of the main parameter of interest $\theta_{h,k}$ is

$$
\begin{aligned}
\widehat{\theta}_{h,k} &= [(\widehat{\boldsymbol{B}}^\top_{X_{\mathrm{fa}(h)}}\overline{\sigma}_Y^{-2}\boldsymbol{V}\widehat{\boldsymbol{B}}_{X_{\mathrm{fa}(h)}})^{-1}\widehat{\boldsymbol{B}}^\top_{X_{\mathrm{fa}(h)}}\overline{\sigma}_Y^{-2}\boldsymbol{V}\widehat{\boldsymbol{\beta}}_{Y_k}]_1\\
&= [(\widehat{\boldsymbol{B}}^{*\top}_{X_{\mathrm{fa}(h)}}\widehat{\boldsymbol{B}}^{*}_{X_{\mathrm{fa}(h)}})^{-1}\widehat{\boldsymbol{B}}^{*\top}_{X_{\mathrm{fa}(h)}}\widehat{\boldsymbol{\beta}}^{*}_{Y_k}]_1,
\end{aligned}
$$

where $\overline{\sigma}_Y^2\boldsymbol{V}^{-1} = q^{-1}\sum_{k\in K}\mathbb{V}(\widehat{\boldsymbol{\beta}}_{Y_k})$ [3] with $\boldsymbol{V}$ the LD matrix of the population from which the traits are drawn.

**Remark 4.** Assuming linearity, no interactions and no unmeasured confounders, back-door adjustment [4] can be applied to derive the effects of a mediator unconfounded by other mediators, *i.e.*, the mediation parameter $\gamma^X_{h,j}$ between the $h$th and $j$th exposures,

$j \neq h \in J$, unconfounded by other exposures and $\gamma^Y_{h,k}$ between the $h$th and $k$th outcomes $j \neq h \in J$, unconfounded by other outcomes and/or exposures. In the case of unmeasured confounders, as in the proposed MR framework, mediation parameters can be derived, but they do not have a causal interpretation. This is due to the selection of IVs discussed in the main text Section 'Selection of instrumental variables'. When taken on their own, the group of exposures violates condition IV3 (each exposure is not conditionally independent of $\boldsymbol{G}$ given the other exposures and $U$) and the group of outcomes violates condition IV2 (each outcome is independent of $\boldsymbol{G}$).

#### Covariances between outcomes and exposures conditionally on $\boldsymbol{G}$

We show that, under MCC for valid IVs, $\text{Cov}(X_j, X_h \mid \boldsymbol{G} = \boldsymbol{g})$, $j, h \in J$, $\text{Cov}(Y_k, X_j \mid \boldsymbol{G} = \boldsymbol{g})$, $k \in K, j \in J$, and $\text{Cov}(Y_k, Y_h \mid \boldsymbol{G} = \boldsymbol{g})$, $k, h \in K$ is unaffected by $U$. The proofs use the property of the cross-moment of two random variables for which it is possible to compute the conditional expectation of one variable while conditioning on the other, and then compute the conditional expectation of the second variable. Given that the expectations are additive in the conditioning variables, we integrate the exposures with respect to their conditional distributions. This strategy is repeated until there are no exposures that have other exposures as parents, and thus, there are no cross-moments of exposures after computing their conditional expectation. Finally, we integrate out $U$ with regard to its marginal distribution.

**Proposition 4.** For a given DAG $\mathcal{D}$ that represents the conditional dependencies between $\boldsymbol{Y}$, $\boldsymbol{X}$, $\boldsymbol{G}$ and $U$, $\text{Cov}(X_h, X_j \mid \boldsymbol{G} = \boldsymbol{g})$, $h, j \in J$, is unconfounded by $U$.

*Proof.* Let's assume that MCC hold and, for a given DAG, among the exposures, the parents of $X_j$ are $\boldsymbol{X}_{\text{pa}(j)}$ and the parents of $X_h$ are $\boldsymbol{X}_{\text{pa}(h)}$ which, in turn, do not have any exposures among their parents. Since $\text{Cov}(X_h, X_j \mid \boldsymbol{G} = \boldsymbol{g}) = \mathbb{E}(X_j X_h \mid \boldsymbol{G} = \boldsymbol{g}) - \mathbb{E}(X_j \mid \boldsymbol{G} = \boldsymbol{g})\, \mathbb{E}(X_h \mid \boldsymbol{G} = \boldsymbol{g})$, and we have already shown the consistency of the coefficients of the a regression of an exposure on $\boldsymbol{G}$ (see Proposition 1), we need to prove that $\mathbb{E}(X_j X_h \mid \boldsymbol{G}=\boldsymbol{g})$ can be estimated without adjustment on $U$. We have

$$
\begin{aligned}
\mathbb{E}(X_j X_h \mid \boldsymbol{G}=\boldsymbol{g}) &= \mathbb{E}_{U|G=g}\,\mathbb{E}_{X_{\mathrm{pa}(j)}|G=g,U}\,\mathbb{E}_{X_{\mathrm{pa}(h)}|X_{\mathrm{pa}(j)},G=g,U}\,\mathbb{E}_{X_h|X_{\mathrm{pa}(j)},X_{\mathrm{pa}(h)},G=g,U}\\
&\quad \mathbb{E}(X_j X_h \mid X_h, \boldsymbol{X}_{\mathrm{pa}(j)}, \boldsymbol{X}_{\mathrm{pa}(h)}, \boldsymbol{G}=\boldsymbol{g}, U)\\
&= \mathbb{E}_{U}\,\mathbb{E}_{X_{\mathrm{pa}(j)}|G=g,U}\,\mathbb{E}_{X_{\mathrm{pa}(h)}|X_{\mathrm{pa}(j)},G=g,U}\,\mathbb{E}_{X_h|X_{\mathrm{pa}(j)},X_{\mathrm{pa}(h)},G=g,U}\\
&\quad \mathbb{E}(X_j X_h \mid X_h, \boldsymbol{X}_{\mathrm{pa}(j)}, \boldsymbol{G}=\boldsymbol{g}, U) \text{ since IV1 and MP}\\
&= \mathbb{E}_{U}\,\mathbb{E}_{X_{\mathrm{pa}(j)}|G=g,U}\,\mathbb{E}_{X_{\mathrm{pa}(h)}|X_{\mathrm{pa}(j)},G=g,U}\\
&\quad \mathbb{E}_{X_h|X_{\mathrm{pa}(j)},X_{\mathrm{pa}(h)},G=g,U}(\{\boldsymbol{g}^\top\boldsymbol{\beta}_{X_j} + \sum_{r\in\mathrm{pa}(j)}\gamma^X_{r,j}X_r + \psi_X U\}X_h)\\
&= \boldsymbol{g}^\top\boldsymbol{\beta}_{X_j}\underbrace{\mathbb{E}_{U}\,\mathbb{E}_{X_{\mathrm{pa}(h)}|G=g,U}\,\mathbb{E}_{X_h|X_{\mathrm{pa}(h)},G=g,U}\,X_h}_{T_1} +\\
&\quad \sum_{r\in\mathrm{pa}(j)}\gamma^X_{r,j}\underbrace{\mathbb{E}_{U}\,\mathbb{E}_{X_r|G=g,U}\,\mathbb{E}_{X_{\mathrm{pa}(h)}|X_r,G=g,U}\,\mathbb{E}_{X_h|X_r,X_{\mathrm{pa}(h)},G=g,U}(X_r X_h)}_{T_2} +\\
&\quad \psi_X\underbrace{\mathbb{E}_{U}\,\mathbb{E}_{X_{\mathrm{pa}(h)}|G=g,U}\,\mathbb{E}_{X_h|X_{\mathrm{pa}(h)},G=g,U}(U X_h)}_{T_3}.
\end{aligned}
\tag{10}
$$

Under the assumption regarding the parents of $X_h$, we have

$$
\begin{aligned}
T_1 &= \mathbb{E}_{U}\,\mathbb{E}_{X_{\mathrm{pa}(h)}|G=g,U}(\boldsymbol{g}^\top\boldsymbol{\beta}_{X_h} + \sum_{s\in\mathrm{pa}(h)}\gamma^X_{s,h}X_s + \psi_X U\})\\
&= \boldsymbol{g}^\top\boldsymbol{\beta}_{X_h} + \mathbb{E}_U \sum_{s\in\mathrm{pa}(h)}\gamma^X_{s,h}\,\mathbb{E}_{X_s|G=g,U}\,X_s + \psi_X\,\mathbb{E}\,U\\
&= \boldsymbol{g}^\top\boldsymbol{\beta}_{X_h} + \sum_{s\in\mathrm{pa}(h)}\gamma^X_{s,h}(\boldsymbol{g}^\top\boldsymbol{\beta}_{X_s} + \psi_X\,\mathbb{E}\,U) + \psi_X\,\mathbb{E}\,U\\
&= \boldsymbol{g}^\top\boldsymbol{\beta}_{X_h} + \sum_{s\in\mathrm{pa}(h)}\gamma^X_{s,h}\boldsymbol{g}^\top\boldsymbol{\beta}_{X_s} + \sum_{s\in\mathrm{pa}(h)}\gamma^X_{s,h}\psi_X\,\mathbb{E}\,U + \psi_X\,\mathbb{E}\,U,
\end{aligned}
\tag{11}
$$

$$
\begin{aligned}
T_2 &= \mathbb{E}_{U}\,\mathbb{E}_{X_r|G=g,U}\,\mathbb{E}_{X_{\mathrm{pa}(h)}|X_r,G=g,U}\,\mathbb{E}_{X_h|X_r,X_{\mathrm{pa}(h)},G=g,U}(X_r X_h)\\
&= \mathbb{E}_{U}\,\mathbb{E}_{X_r|G=g,U}\,\mathbb{E}_{X_{\mathrm{pa}(h)}|X_r,G=g,U}(X_r\{\boldsymbol{g}^\top\boldsymbol{\beta}_{X_h} + \sum_{s\in\mathrm{pa}(h)}\gamma^X_{s,h}X_s + \psi_X U\})\\
&= \mathbb{E}_{U}\,\mathbb{E}_{X_r|G=g,U}(X_r\{\boldsymbol{g}^\top\boldsymbol{\beta}_{X_h} + \sum_{s\in\mathrm{pa}(h)}\gamma^X_{s,h}\,\mathbb{E}_{X_s|G=g,U}\,X_s + \psi_X U\})\\
&= \boldsymbol{g}^\top\boldsymbol{\beta}_{X_h}\,\mathbb{E}_{U}\,\mathbb{E}_{X_r|G=g,U}\,X_r + \mathbb{E}_{U}\,\mathbb{E}_{X_r|G=g,U}(X_r\psi_X U) +\\
&\quad \mathbb{E}_U(\mathbb{E}_{X_r|G=g,U}\,X_r\sum_{s\in\mathrm{pa}(h)}\gamma^X_{s,h}\,\mathbb{E}_{X_s|G=g,U}\,X_s)\\
&= \boldsymbol{g}^\top\boldsymbol{\beta}_{X_h}\boldsymbol{g}^\top\boldsymbol{\beta}_{X_r} + \boldsymbol{g}^\top\boldsymbol{\beta}_{X_h}\psi_X\,\mathbb{E}\,U + \boldsymbol{g}^\top\boldsymbol{\beta}_{X_r}\psi_X\,\mathbb{E}\,U + \psi_X^2\,\mathbb{E}\,U^2 +\\
&\quad \boldsymbol{g}^\top\boldsymbol{\beta}_{X_r}\sum_{s\in\mathrm{pa}(h)}\gamma^X_{s,h}\boldsymbol{g}^\top\boldsymbol{\beta}_{X_s} + \boldsymbol{g}^\top\boldsymbol{\beta}_{X_r}\sum_{s\in\mathrm{pa}(h)}\gamma^X_{s,h}\psi_X\,\mathbb{E}\,U +\\
&\quad \psi_X\,\mathbb{E}\,U\sum_{s\in\mathrm{pa}(h)}\gamma^X_{s,h}\boldsymbol{g}^\top\boldsymbol{\beta}_{X_s} + \sum_{s\in\mathrm{pa}(h)}\gamma^X_{s,h}\psi_X^2\,\mathbb{E}\,U^2,
\end{aligned}
\tag{12}
$$

and

$$
\begin{aligned}
T_3 &= \mathbb{E}_U \, \mathbb{E}_{X_{\text{pa}(h)}|G=g,U}(U\{\boldsymbol{g}^\top \boldsymbol{\beta}_{X_h} + \sum_{s\in\text{pa}(h)} \gamma^X_{s,h} X_s + \psi_X U\}) \\
&= \boldsymbol{g}^\top \boldsymbol{\beta}_{X_h} \, \mathbb{E}\, U + \mathbb{E}_U \sum_{s\in\text{pa}(h)} \gamma^X_{s,h} U \, \mathbb{E}_{X_s|G,U} X_s + \psi_X \, \mathbb{E}\, U^2 \\
&= \boldsymbol{g}^\top \boldsymbol{\beta}_{X_h} \, \mathbb{E}\, U + \sum_{s\in\text{pa}(h)} \gamma^X_{s,h} (\boldsymbol{g}^\top \boldsymbol{\beta}_{X_s} \, \mathbb{E}\, U + \psi_X \, \mathbb{E}\, U^2) + \psi_X \, \mathbb{E}\, U^2 \\
&= \boldsymbol{g}^\top \boldsymbol{\beta}_{X_h} \, \mathbb{E}\, U + \sum_{s\in\text{pa}(h)} \gamma^X_{s,h} \boldsymbol{g}^\top \boldsymbol{\beta}_{X_s} \, \mathbb{E}\, U + \sum_{s\in\text{pa}(h)} \gamma^X_{s,h} \psi_X \, \mathbb{E}\, U^2 + \psi_X \, \mathbb{E}\, U^2.
\end{aligned} \tag{13}
$$

Combining (10), (11), (12) and (13), we obtain

$$
\begin{aligned}
\mathbb{E}(X_j X_h \mid \boldsymbol{G} = \boldsymbol{g}) = {}& \boldsymbol{g}^\top \boldsymbol{\beta}_{X_j} \boldsymbol{g}^\top \boldsymbol{\beta}_{X_h} + \boldsymbol{g}^\top \boldsymbol{\beta}_{X_j} \sum_{s\in\text{pa}(h)} \gamma^X_{s,h} \boldsymbol{g}^\top \boldsymbol{\beta}_{X_s} + \boldsymbol{g}^\top \boldsymbol{\beta}_{X_h} \sum_{r\in\text{pa}(j)} \gamma^X_{r,j} \boldsymbol{g}^\top \boldsymbol{\beta}_{X_r} + \\
& \sum_{r\in\text{pa}(j)} \gamma^X_{r,j} \boldsymbol{g}^\top \boldsymbol{\beta}_{X_r} \sum_{s\in\text{pa}(h)} \gamma^X_{s,h} \boldsymbol{g}^\top \boldsymbol{\beta}_{X_s} + \\
& \{(1 + \sum_{s\in\text{pa}(h)} \gamma^X_{s,h}) \boldsymbol{g}^\top \boldsymbol{\beta}_{X_j} + (1 + \sum_{r\in\text{pa}(j)} \gamma^X_{r,j}) \boldsymbol{g}^\top \boldsymbol{\beta}_{X_h} + (1 + \sum_{s\in\text{pa}(h)} \gamma^X_{s,h}) \\
& \sum_{r\in\text{pa}(j)} \gamma^X_{r,j} \boldsymbol{g}^\top \boldsymbol{\beta}_{X_r} + (1 + \sum_{r\in\text{pa}(j)} \gamma^X_{r,j}) \sum_{s\in\text{pa}(h)} \gamma^X_{s,h} \boldsymbol{g}^\top \boldsymbol{\beta}_{X_s}\} \psi_X \, \mathbb{E}\, U + \\
& (1 + \sum_{r\in\text{pa}(j)} \gamma^X_{r,j} + \sum_{s\in\text{pa}(h)} \gamma^X_{s,h} + \sum_{r\in\text{pa}(j)} \gamma^X_{r,j} \sum_{s\in\text{pa}(h)} \gamma^X_{s,h}) \psi_X^2 \, \mathbb{E}\, U^2 \\
= {}& \alpha_{X_j X_h} + \boldsymbol{g}^\top \boldsymbol{\beta}_{X_j} \boldsymbol{g}^\top \boldsymbol{\beta}_{X_h} + \boldsymbol{g}^\top \boldsymbol{\beta}_{X_j} \sum_{s\in\text{pa}(h)} \gamma^X_{s,h} \boldsymbol{g}^\top \boldsymbol{\beta}_{X_s} + \\
& \boldsymbol{g}^\top \boldsymbol{\beta}_{X_h} \sum_{r\in\text{pa}(j)} \gamma^X_{r,j} \boldsymbol{g}^\top \boldsymbol{\beta}_{X_r} + \sum_{r\in\text{pa}(j)} \gamma^X_{r,j} \boldsymbol{g}^\top \boldsymbol{\beta}_{X_r} \sum_{s\in\text{pa}(h)} \gamma^X_{s,h} \boldsymbol{g}^\top \boldsymbol{\beta}_{X_s}.
\end{aligned}
$$

■

**Proposition 5.** For a given DAG $\mathcal{D}$ that represents the conditional dependencies between $\boldsymbol{Y}$, $\boldsymbol{X}$, $\boldsymbol{G}$ and $U$, $\text{Cov}(Y_k, X_h \mid \boldsymbol{G} = \boldsymbol{g})$, $k \in K, h \in J$ and $\text{Cov}(Y_k, Y_h \mid \boldsymbol{G} = \boldsymbol{g})$, $k, h \in K$ are unconfounded by $U$.

*Proof.* Given condition IV3, all information between the outcomes and the IVs is mediated by the exposures (and the unmeasured confounder). Thus, it is sufficient to show that by computing the conditional expectation of an outcome, we can obtain a linear combination of conditional cross-moments and conditional expectations of exposures as in (10).

Let's assume that MCC hold and, for a given DAG, the parents of $Y_k$ are $\boldsymbol{Y}_{\text{pa}(k)}$ and $\boldsymbol{X}_{\text{pa}(k)}$ and $X_h$ has parents $\boldsymbol{X}_{\text{pa}(h)}$. Finally, let's assume that $\boldsymbol{Y}_{\text{pa}(k)}$ do not have any outcomes among their parents and both $\boldsymbol{X}_{\text{pa}(k)}$ and $\boldsymbol{X}_{\text{pa}(h)}$ have parents that we denote by $\boldsymbol{X}_{\text{pa}(\text{pa}(k))}$ and $\boldsymbol{X}_{\text{pa}(\text{pa}(h))}$, respectively, which, in turn, do not have any exposures among their parents.

Since $\text{Cov}(Y_k, X_h \mid \boldsymbol{G} = \boldsymbol{g}) = \mathbb{E}(Y_k X_h \mid \boldsymbol{G} = \boldsymbol{g}) - \mathbb{E}(Y_k \mid \boldsymbol{G} = \boldsymbol{g}) \, \mathbb{E}(X_h \mid \boldsymbol{G} = \boldsymbol{g})$, and we have already shown that the regressions of each outcome and exposure on $\boldsymbol{G}$ are unaffected by $U$ (see Proposition 1), we need to prove that $\mathbb{E}(Y_k X_h \mid \boldsymbol{G} = \boldsymbol{g})$ can be estimated without adjustment on $U$. We have

$$\begin{aligned}
\mathbb{E}(Y_k X_h \mid \boldsymbol{G}=\boldsymbol{g}) &= \mathbb{E}_{U|G=g}\,\mathbb{E}_{X_{\mathrm{pa}(\mathrm{pa}(k))}|G=g,U}\,\mathbb{E}_{X_{\mathrm{pa}(k)}|X_{\mathrm{pa}(\mathrm{pa}(k))},G=g,U}\,\mathbb{E}_{Y_{\mathrm{pa}(k)}|X_{\mathrm{pa}(k)},X_{\mathrm{pa}(\mathrm{pa}(k))},G=g,U}\\
&\quad \mathbb{E}_{X_{\mathrm{pa}(\mathrm{pa}(h))}|Y_{\mathrm{pa}(k)},X_{\mathrm{pa}(k)},X_{\mathrm{pa}(\mathrm{pa}(k))},G=g,U}\\
&\quad \mathbb{E}_{X_{\mathrm{pa}(h)}|X_{\mathrm{pa}(\mathrm{pa}(h))},Y_{\mathrm{pa}(k)},X_{\mathrm{pa}(k)},X_{\mathrm{pa}(\mathrm{pa}(k))},G=g,U}\\
&\quad \mathbb{E}_{X_h|X_{\mathrm{pa}(h)},X_{\mathrm{pa}(\mathrm{pa}(h))},Y_{\mathrm{pa}(k)},X_{\mathrm{pa}(k)},X_{\mathrm{pa}(\mathrm{pa}(k))},G=g,U}\\
&\quad \mathbb{E}(Y_k X_h \mid X_h, \boldsymbol{X}_{\mathrm{pa}(h)}, \boldsymbol{X}_{\mathrm{pa}(\mathrm{pa}(h))}, \boldsymbol{Y}_{\mathrm{pa}(k)}, \boldsymbol{X}_{\mathrm{pa}(k)}, \boldsymbol{X}_{\mathrm{pa}(\mathrm{pa}(k))}, \boldsymbol{G}=\boldsymbol{g}, U)\\
&= \mathbb{E}_U\,\mathbb{E}_{X_{\mathrm{pa}(\mathrm{pa}(k))}|G=g,U}\,\mathbb{E}_{X_{\mathrm{pa}(k)}|X_{\mathrm{pa}(\mathrm{pa}(k))},G=g,U}\,\mathbb{E}_{Y_{\mathrm{pa}(k)}|X_{\mathrm{pa}(k)},X_{\mathrm{pa}(\mathrm{pa}(k))},U}\\
&\quad \mathbb{E}_{X_{\mathrm{pa}(\mathrm{pa}(h))}|X_{\mathrm{pa}(k)},X_{\mathrm{pa}(\mathrm{pa}(k))},G=g,U}\,\mathbb{E}_{X_{\mathrm{pa}(h)}|X_{\mathrm{pa}(\mathrm{pa}(h))},X_{\mathrm{pa}(k)},X_{\mathrm{pa}(\mathrm{pa}(k))},G=g,U}\\
&\quad \mathbb{E}_{X_h|X_{\mathrm{pa}(h)},X_{\mathrm{pa}(\mathrm{pa}(h))},X_{\mathrm{pa}(k)},X_{\mathrm{pa}(\mathrm{pa}(k))},G=g,U}\\
&\quad \mathbb{E}(Y_k X_h \mid X_h, \boldsymbol{Y}_{\mathrm{pa}(k)}, \boldsymbol{X}_{\mathrm{pa}(k)}, \boldsymbol{G}=\boldsymbol{g}, U)\\
&\quad \text{since IV1, IV3 and MP}\\
&= \mathbb{E}_U(\{\sum_{r\in\mathrm{pa}(k)} \theta_{r,k}\,\mathbb{E}_{X_{\mathrm{pa}(r)}|G=g,U}\,\mathbb{E}_{X_r|X_{\mathrm{pa}(r)},G=g,U}\,X_r +\\
&\quad \sum_{t\in\mathrm{pa}(k)} \gamma^Y_{t,k}\,\mathbb{E}_{X_{\mathrm{pa}(t)}|G=g,U}\,\mathbb{E}_{Y_t|X_{\mathrm{pa}(t)},U}\,Y_t + \psi_Y U\}\\
&\quad \{\boldsymbol{g}^\top \boldsymbol{\beta}_{X_h} + \sum_{s\in\mathrm{pa}(h)} \gamma^X_{s,h}\,\mathbb{E}_{X_s|X_{\mathrm{pa}(s)},G=g,U}\,X_s + \psi_X U\})\\
&= \{\sum_{r\in\mathrm{pa}(k)} \theta_{r,k}\,\mathbb{E}_U\,\mathbb{E}_{X_{\mathrm{pa}(r)}|G=g,U}\,\mathbb{E}_{X_r|X_{\mathrm{pa}(r)},G=g,U}\,X_r +\\
&\quad \sum_{t\in\mathrm{pa}(k)} \gamma^Y_{t,k} \sum_{v\in\mathrm{pa}(t)} \theta_{v,t}\,\mathbb{E}_U\,\mathbb{E}_{X_v|G=g,U}\,X_v + \sum_{t\in\mathrm{pa}(k)} \gamma^Y_{t,k}\psi_Y\,\mathbb{E}\,U + \psi_Y\,\mathbb{E}\,U\}\\
&\quad \{\boldsymbol{g}^\top \boldsymbol{\beta}_{X_h} + \sum_{s\in\mathrm{pa}(h)} \gamma^X_{s,h}\,\mathbb{E}_U\,\mathbb{E}_{X_s|X_{\mathrm{pa}(s)},G=g,U}\,X_s + \psi_X\,\mathbb{E}\,U\}.
\end{aligned}$$

Similarly, let's assume that MCC hold and, for a given DAG, the parents of $Y_k$ are $\boldsymbol{Y}_{\mathrm{pa}(k)}$ and $\boldsymbol{X}_{\mathrm{pa}(k)}$ and the parents of $Y_h$ are $\boldsymbol{Y}_{\mathrm{pa}(h)}$ and $\boldsymbol{X}_{\mathrm{pa}(h)}$. Finally, let's assume that neither $\boldsymbol{Y}_{\mathrm{pa}(k)}$ nor $\boldsymbol{Y}_{\mathrm{pa}(h)}$ have any outcomes among their parents and both $\boldsymbol{X}_{\mathrm{pa}(k)}$ and $\boldsymbol{X}_{\mathrm{pa}(h)}$ have parents that we denote by $\boldsymbol{X}_{\mathrm{pa}(\mathrm{pa}(h))}$ and $\boldsymbol{X}_{\mathrm{pa}(\mathrm{pa}(h))}$, respectively, which, in turn, do not have any exposures among their parents. The conditional cross-moment is

$$\begin{aligned}
\mathbb{E}(Y_k Y_h \mid \boldsymbol{G}=\boldsymbol{g}) = {} & \mathbb{E}_{U|G=g}\, \mathbb{E}_{X_{\mathrm{pa}(\mathrm{pa}(k))}|G=g,U}\, \mathbb{E}_{X_{\mathrm{pa}(k)}|X_{\mathrm{pa}(\mathrm{pa}(k))},G=g,U}\, \mathbb{E}_{Y_{\mathrm{pa}(k)}|X_{\mathrm{pa}(k)},X_{\mathrm{pa}(\mathrm{pa}(k))},G=g,U} \\
& \mathbb{E}_{X_{\mathrm{pa}(\mathrm{pa}(h))}|Y_{\mathrm{pa}(k)},X_{\mathrm{pa}(k)},X_{\mathrm{pa}(\mathrm{pa}(k))},G=g,U} \\
& \mathbb{E}_{X_{\mathrm{pa}(h)}|X_{\mathrm{pa}(\mathrm{pa}(h))},Y_{\mathrm{pa}(k)},X_{\mathrm{pa}(k)},X_{\mathrm{pa}(\mathrm{pa}(k))},G=g,U} \\
& \mathbb{E}_{Y_{\mathrm{pa}(h)}|X_{\mathrm{pa}(h)},X_{\mathrm{pa}(\mathrm{pa}(h))},Y_{\mathrm{pa}(k)},X_{\mathrm{pa}(k)},X_{\mathrm{pa}(\mathrm{pa}(k))},G=g,U} \\
& \mathbb{E}_{Y_h|Y_{\mathrm{pa}(h)},X_{\mathrm{pa}(h)},X_{\mathrm{pa}(\mathrm{pa}(h))},Y_{\mathrm{pa}(k)},X_{\mathrm{pa}(k)},X_{\mathrm{pa}(\mathrm{pa}(k))},G=g,U} \\
& \mathbb{E}(Y_k Y_h \mid Y_h, \boldsymbol{Y}_{\mathrm{pa}(h)}\boldsymbol{X}_{\mathrm{pa}(h)}, \boldsymbol{X}_{\mathrm{pa}(\mathrm{pa}(h))}, \boldsymbol{Y}_{\mathrm{pa}(k)}, \boldsymbol{X}_{\mathrm{pa}(k)}, \boldsymbol{X}_{\mathrm{pa}(\mathrm{pa}(k))}, \boldsymbol{G}=\boldsymbol{g}, U) \\
= {} & \mathbb{E}_U\, \mathbb{E}_{X_{\mathrm{pa}(\mathrm{pa}(k))}|G=g,U}\, \mathbb{E}_{X_{\mathrm{pa}(k)}|X_{\mathrm{pa}(\mathrm{pa}(k))},G=g,U}\, \mathbb{E}_{Y_{\mathrm{pa}(k)}|X_{\mathrm{pa}(k)},X_{\mathrm{pa}(\mathrm{pa}(k))},U} \\
& \mathbb{E}_{X_{\mathrm{pa}(\mathrm{pa}(h))}|X_{\mathrm{pa}(k)},X_{\mathrm{pa}(\mathrm{pa}(k))},G=g,U}\, \mathbb{E}_{X_{\mathrm{pa}(h)}|X_{\mathrm{pa}(\mathrm{pa}(h))},X_{\mathrm{pa}(k)},X_{\mathrm{pa}(\mathrm{pa}(k))},G=g,U} \\
& \mathbb{E}_{Y_{\mathrm{pa}(h)}|X_{\mathrm{pa}(h)},X_{\mathrm{pa}(\mathrm{pa}(h))},Y_{\mathrm{pa}(k)},X_{\mathrm{pa}(k)},X_{\mathrm{pa}(\mathrm{pa}(k))},U} \\
& \mathbb{E}_{Y_h|Y_{\mathrm{pa}(h)},X_{\mathrm{pa}(h)},X_{\mathrm{pa}(\mathrm{pa}(h))},Y_{\mathrm{pa}(k)},X_{\mathrm{pa}(k)},X_{\mathrm{pa}(\mathrm{pa}(k))},U} \\
& \mathbb{E}(Y_k Y_h \mid Y_h, \boldsymbol{Y}_{\mathrm{pa}(k)}, \boldsymbol{X}_{\mathrm{pa}(k)}, \boldsymbol{G}=\boldsymbol{g}, U) \\
& \text{since IV1 and MP} \\
= {} & \mathbb{E}_U(\{\sum_{r\in\mathrm{pa}(k)} \theta_{r,k}\, \mathbb{E}_{X_{\mathrm{pa}(r)}|G=g,U}\, \mathbb{E}_{X_r|X_{\mathrm{pa}(r)},G=g,U}\, X_r + \\
& \sum_{t\in\mathrm{pa}(k)} \gamma^Y_{t,k}\, \mathbb{E}_{X_{\mathrm{pa}(t)}|G=g,U}\, \mathbb{E}_{Y_t|X_{\mathrm{pa}(t)},U}\, Y_t + \psi_Y U\} \\
& \{\sum_{s\in\mathrm{pa}(h)} \theta_{s,h}\, \mathbb{E}_{X_{\mathrm{pa}(s)}|G=g,U}\, \mathbb{E}_{X_s|X_{\mathrm{pa}(s)},G=g,U}\, X_s) \\
& \sum_{v\in\mathrm{pa}(h)} \gamma^Y_{v,h}\, \mathbb{E}_{X_{\mathrm{pa}(v)}|G=g,U}\, \mathbb{E}_{Y_v|X_{\mathrm{pa}(h)},U}\, Y_v + \psi_Y U\}) \\
= {} & \{\sum_{r\in\mathrm{pa}(k)} \theta_{r,k}\, \mathbb{E}_U\, \mathbb{E}_{X_{\mathrm{pa}(r)}|G=g,U}\, \mathbb{E}_{X_r|X_{\mathrm{pa}(r)},G=g,U}\, X_r + \\
& \sum_{t\in\mathrm{pa}(k)} \gamma^Y_{t,k} \sum_{w\in\mathrm{pa}(t)} \theta_{w,t}\, \mathbb{E}_U\, \mathbb{E}_{X_w|G=g,U}\, X_w + \sum_{t\in\mathrm{pa}(k)} \gamma^Y_{t,k}\psi_Y\, \mathbb{E}\, U + \psi_Y\, \mathbb{E}\, U\} \\
& \{\sum_{s\in\mathrm{pa}(h)} \theta_{s,h}\, \mathbb{E}_{X_{\mathrm{pa}(s)}|G=g,U}\, \mathbb{E}_U\, \mathbb{E}_{X_s|X_{\mathrm{pa}(s)},G=g,U}\, X_s + \\
& \sum_{v\in\mathrm{pa}(h)} \gamma^Y_{v,h} \sum_{z\in\mathrm{pa}(t)} \theta_{z,v}\, \mathbb{E}_U\, \mathbb{E}_{X_z|G=g,U}\, X_z + \sum_{v\in\mathrm{pa}(h)} \gamma^Y_{v,h}\psi_Y\, \mathbb{E}\, U + \psi_Y\, \mathbb{E}\, U\}.
\end{aligned}$$

■

**Remark 5.** As noted in Proposition 1, to prove that the covariances between the traits are unconfounded by $U$ conditionally on the instrumental variables, we made some assumptions regarding the dependency structure associated with the DAG. The proofs do not change if further dependency relations are considered among the outcomes and the exposures given that is possible to calculate the cross-moment of two random variables by computing the conditional expectation of one variable while conditioning on the other, and then deriving the conditional expectation of the second variable. Assuming general linear models, the expectations are additive in the conditioning variables. This allows us to integrate out the exposures with respect to their conditional distributions and $U$ with regard to its marginal distribution.

### Simulation study and real data application

#### MrDAG parameters setting

In the following, we detail the setting of MrDAG Markov chain Monte Carlo (MCMC) algorithm presented in the main text Methods that we used in the simulation study and the real data application. In the simulation study, MrDAG algorithm is run for $75,000$ MCMC sweeps, of which $25,000$ as burn-in and results saved every 10 sweeps, resulting in $5,000$ posterior samples for all the unknowns. Since *a priori* we expect one edge for each of the $(q+p)$ nodes, the prior probability of edge inclusion in (10) is

$$\begin{aligned}\pi^{\text{edge}} &= \tfrac{q+p}{(q+p)(q+p-1)/2} \\ &= \tfrac{2}{q+p-1},\end{aligned} \tag{14}$$

where $q$ is the number of responses, $p$ is the number of exposures and the denominator is the maximum number of edges in a graph with $(q+p)$ nodes. Using (14) with $q = 5$ and $p = 15$, the prior probability of edge inclusion is $\pi^{\text{edge}} = 0.10$. Finally, the value of the burn-in annealing parameter $T$ is set at 5. No other hyper-parameters need to be defined given the specification of objective priors for model selection based on the fractional Bayes factor, leading to a closed-form expression for the marginal likelihood [5] and the use of objective priors also for the inverse of the covariance matrix [6].

To select a suitable number of MCMC sweeps, and in particular the length of the burn-in, we executed some preliminary runs and checked by visual inspection the trace plots of the marginal likelihood and the model size, *i.e.*, the number of edges selected during the MCMC. No signs of slow convergence or aberrant behaviour of the designed MCMC sampler were identified in a variety of scenarios and parameter settings, suggesting that the number of sweeps during burn-in is sufficient to reach convergence and sample from the equilibrium distribution of the Markov chain.

In the real data analysis, we considerably increase the number of MCMC sweeps to $10^6$ of which $10^5$ as burn-in and results saved every 100 sweep, resulting in $9,000$ posterior samples for all the unknowns. Assuming *a priori* an edge for each of the $(q+p)$ nodes and using (14), with $q = 7$ and $p = 6$, $\pi^{\text{edge}} = 0.16$. Finally, the value of the burn-in annealing parameter $T$ is set at 10. As illustrated in Supplementary Figure 8, the burn-in and the total length of the MCMC far exceed the number of sweeps required to reach convergence to the equilibrium distribution of the Markov chain and explore the graph space faithfully. No signs of slow convergence or aberrant behaviour of the designed MCMC sampler appear. We used the same number of MCMC sweeps, annealing parameter $T$ and $\pi^{\text{edge}}$ to test reverse causation in the real data analysis.

We conclude with the description of MrDAG parameters we used in the bootstrap analysis [7] to check the robustness of the findings in the real data application. We bootstrapped the IVs with replacement $B = 100$ times and ran MrDAG algorithm on each bootstrap summary-level statistics. All parameters are left unchanged compared to the original real data analysis, $\pi^{\text{edge}} = 0.16$ and $T = 10$. However, due to the expensive computational time of each run, we decreased the MCMC sweeps to $75,000$, of which $25,000$ as burn-in without thinning, resulting in $50,000$ posterior samples for each bootstrap sample. The length of the burn-in was suggested by looking at the trace plot of the marginal-log likelihood in a few trail runs. Overall, considering the whole bootstrap analysis, we obtain $7.5 \times 10^6$ posterior samples, of which $5 \times 10^6$ after burn-in. Finally, we calculate the bootstrap frequency of edge inclusion as the average of the posterior probability of edge inclusion (PPeI) obtained in each bootstrap sample.

##### Alternative methods parameters setting

In the simulation study, as an alternative method we would like to compare, we include MR-BMA [8], a Bayesian variable selection approach for multivariable MR, with a prior probability of inclusion set at 0.10. In this way, we match the prior specification of the edge inclusion probability in MrDAG. MR-BMA only considers one outcome at-a-time and is performed on each response separately. We use also MR-BMA in the real data application and we set the prior probability of inclusion set at 0.16 to match that used in MrDAG.

We also include MRPC algorithm [9] based on [10]. It requires the specification of the "sufficient statistic" for the data and the number of "genetic variants". For the former, we calculate the Pearson correlation between the summary-level statistics and provide the number of simulated IVs, $n = 100$. For the latter, we indicate the number of exposures simulated, *i.e.*, $p = 15$. Since the genetic associations with the exposures and the responses are normally distributed, we select `gaussCItest` option to test the conditional independence between the simulated summary-level statistics. We do not correct for multiplicity. Instead, a type I error rate $\alpha$ is used for the conditional independence test. The detected Partial DAG (PDAG) are obtained at different values of $\alpha = \{0.01, 0.05, 0.10, 0.20\}$. Subsequently, they are used to estimate causal effects under intervention by using the function `o.ida()` in the R package *pcalg* [11] which considers "only those edges necessary to compute the possible optimal valid adjustment sets, using these as adjustment sets to estimate the unique possible causal effects" and specifying the option `type = "pdag"`.

We also consider Partition DAG algorithm [12] implemented in the R package *PDAG* [13]. It performs structure learning under constraints and causal effect estimation based on Lasso penalisation of the negative log-likelihood. The constraints correspond to a pre-specified partition of the nodes in which domain-specific knowledge provides information regarding their partial ordering. In our setup, the prior knowledge allows us to partition the nodes into exposures and outcomes and the partial ordering information corresponds to the edges' orientation from the exposures to the responses. Since two disjoint groups of nodes are considered, we use the function `partitionDAG::partial2()` and specify `m1 = 15` where the partition between the exposures and the outcomes occurs. Results are recorded for three different values for the Lasso penalisation $\lambda = \{0.5, 0.7, 0.9\}$. Preliminary runs suggest using these values since at zero no penalisation occurs with extremely dense solutions and at 1 the shrinkage is too strong. We also tried a stability selection approach to set the penalisation parameter as suggested in [12]. However, in all simulation scenarios, the estimated penalisation parameter $\widehat{\lambda}$ is quite small, ranging from 0.05 to 0.10 with dense solutions. We do not report these results given that they were not sufficiently sparse.

##### Definition of True Positive, False Negative and False Positive

To evaluate the performance of the methods considered and summarize it by a precision-recall curve (PRC), with recall (= sensitivity = TP/(TP + FN)) in the $x$-axis and precision (= positive predictive value = TP/(TP + FP)) in the $y$-axis, a definition of TP = True Positive, FN = False Negative and FN = False Positive is required.

However, since the methods considered in the simulation study are quite heterogeneous, there are substantial differences in what they estimate. MrDAG and MRPC algorithms return as output PDAGs. In MrDAG this happens when, at a defined threshold of the PPeI, there are bidirected edges [5], while in the MRPC algorithm, this occurs since undirected edges might still exist after determining all $v$-structures and applying rules R1-R3 of the PC algorithm as given in Algorithm 2 detailed in [14]. In contrast, Partition DAG provides a fully oriented DAG as output.

For a fair comparison, it is paramount to define TP, FN and FP in a way that they do not favour implicitly any method. We depart from [15, 5] who transform a simulated DAG (*true* DAG) into a CPDAG and [12] who modify a CPDAG output by reorienting all undirected edges according to the simulated DAG (as well as to the opposite direction). Instead, we compare the results of the different methods by considering the simulated *true* (partially- and fully-oriented) DAG as described in the main text Methods.

We motivate this choice as follows:

- Transforming a *true* DAG into a CPDAG might violate the partial ordering. If the resulting CPDAG is considered as the background truth and a violation of the partial ordering is present, all methods will experience an increase of the FNs since either they automatically satisfy the constraint of the causal effects from the exposures to the responses (MR-BMA) or this condition is enforced (MRPC, ParDAG and MrDAG).

- In MRPC and MrDAG, undirected edges might exist only within the exposures and outcomes given the partial ordering enforced between the phenotypes. Reorienting undirected edges according to the *true* DAG can be a solution, but it provides a substantial advantage to these methods. Moreover, the divergence of the results when reorienting the undirected edges based on the true DAG or flipping to the opposite direction might be so large that it is not clear their overall performance.

- Since in two scenarios of the simulation study DAGs are fully oriented and in two scenarios they are partially oriented (but only within the exposures), this should provide alternative advantages to the competing algorithms and, overall, a fair comparison.

We define TP when an algorithm can detect the simulated oriented edge, FN when an algorithm does not declare any oriented edge when instead it should and FP when an algorithm detects an oriented edge when instead it shouldn't. In the case of partially oriented DAGs, TP, FN and FP are defined with respect to the detection of bidirectional edges. In practice, their estimation is simplified by utilising the simulated and estimated non-symmetrical adjacency matrix for each replicate and in each scenario (see in the main text Figure 1).

#### Further results of MrDAG model in the real data application

Here, we present extended results concerning the findings obtained by alternative methods to detect the impact of lifestyle and behavioural traits on mental health phenotypes and internal checks of the results provided by MrDAG algorithm when applied to the same data set.

The risk of detecting spurious shared causal effects is very high when a standard MR method is used separately on each trait as well as when multiple exposures are considered for each outcome [8]. To show that this is a widespread issue which can be solved only if dependency relations are considered in the model, we contrast the results of MrDAG with multivariable single-response MR, implemented in the MR-BMA algorithm which can perform exposure selection [8] but is not designed for multiple outcomes.

Supplementary Table 2 shows the results MR-BMA algorithm when applied to the same data set. By contrasting Figure 6 in the main text and Supplementary Table 2 some general comments can be made. First, the signs of the causal effects from the exposures to the outcomes mostly agree, for instance, the causal effects under intervention on EDU, although the effects sizes are almost two times bigger than those estimated by MrDAG, except for the exposure-outcome pair EDU-COG where they coincide. Second, the overestimation of the causal effects by MR-BMA is a general feature since the pleiotropic effects within the outcomes are not considered in MR-BMA which tries to ascribe the whole effect to the exposures. See the effect of genetically predicted lifetime smoking index (SM) on ADHD. Third, as expected from the simulation study, MR-BMA detects many more associations than MrDAG. For instance, genetically predicated SP has been found causally associated two times (with AN and SCZ), ALC once (ASD), and LST five times (AN, ADHD, ASD, BD, and SCZ). Except for the causal association of LST with AN, MrDAG does not identify any of these associations. Finally, MR-BMA detects genetically predicated SM on SCZ while MrDAG lists it as a spurious causal effect.

Internal validation is an important step to assess the validity of the results obtained by MrDAG algorithm. We divide this internal check into sensitivity to hyper-prior specifications and robustness of structure learning.

Regarding the first point, MrDAG is built under an objective Bayes framework which has the advantage of not depending on priors hyper-parameters (see Methods in the main text). The only parameter that needs to be specified is the prior probability of edge inclusion that controls the level of sparsity. The results presented in the main text Section 'Real data application: The impact of lifestyle and behavioural traits on mental health' were obtained by setting $\pi^{\text{edge}} = 0.16$, *i.e.*, *a priori* we expect one edge for each of the 13 nodes. We repeat the analysis, and specify $\pi^{\text{edge}} = 0.08$ and $\pi^{\text{edge}} = 0.04$. Supplementary Figure 11 shows that the posterior causal effects as well as the 95% credible intervals (CIs) are not influenced by this choice, with only a handful of cases at $\pi^{\text{edge}} = 0.04$ where the posterior causal effects and the CIs are slightly different.

Although MrDAG explores the space of alternative Essential Graphs (EGs) [16] that best fit the data and produces the posterior estimate of the causal effects averaging over the space of the visited graphical models and thus, taking into account model uncertainty without focusing on a single model, the second question is how robust are the results and how much they should be trusted. We answer these questions by bootstrapping MrDAG repeatedly on the data [7]. In Supplementary Figures 12 we present the bootstrap frequency of edge inclusion for each permitted combination of exposures and outcomes and the scatterplot of the posterior probability of edge inclusion (PPeI) against the bootstrap frequency of edge inclusion. The results show that there is a satisfactory agreement between a single run of the algorithm and the bootstrap results for the casual associations. In four cases we report less alignment. The exposure-outcome pairs SM-SCZ and PA-MDD receive more weight in the bootstrap analysis than in the single run of the algorithm, while the opposite happens for EDU-BD and SM-COG, although, in all cases, the bootstrap frequency of edge inclusion is around 50%.

### 2 Supplementary Tables

| Type | Trait | Acronym | Population | Sample size | No. IVs (%) | Reference | Data source |
|---|---|---|---|---|---|---|---|
| Expsoure | Education (in years) | EDU | European (meta-analysis) | 766,345 | $426^{\dagger}$ (0.57) | Lee *et. al.*, 2018 [17] | `https://thessgac.com/` |
| | Physical activity | PA | European (meta-analysis + UK Biobank) | 608,595 | $5^{\dagger}$ (0.01) | Wang *et. al.*, 2022 [18] | `https://www.ebi.ac.uk/gwas/studies/GCST90104341` |
| | Overall sleep duration | SP | European (UK Biobank) | 446,118 | $56^{\dagger}$ (0.07) | Dashi *et. al.*, 2019 [19] | `https://sleep.hugeamp.org/dinspector.html?dataset=GWAS_UKBB_eu` |
| | Alcohol consumption | ALC | European (UK Biobank) | 480,842 | $63^{\dagger}$ (0.08) | Evangelou *et. al.*, 2019 [20] | `http://ftp.ebi.ac.uk/pub/databases/gwas/summary_statistics/` |
| | Lifetime smoking index | SM | European (UK Biobank) | 462,690 | $111^{\dagger}$ (0.15) | Wootton *et. al.*, 2020 [21] | `https://data.bris.ac.uk/data/dataset/10i96zb8gm0j81yz0q6ztei23d` |
| | Leisure screen time | LST | European (meta-analysis + UK Biobank) | 526,725 | $92^{\dagger}$ (0.12) | Wang *et. al.*, 2022 [18] | `https://www.ebi.ac.uk/gwas/studies/GCST90104339` |
| Outcome | Major depressive disorder | MDD | European (meta-analysis) | 170,756 cases and 329,443 controls | $45^{\ddagger}$ (0.10) | Howard *et. al.*, 2019 [22] | |
| | Anorexia nervosa | AN | European (meta-analysis) | 16,992 cases and 55,525 controls | $5^{\ddagger}$ (0.01) | Watson *et. al.*, 2019 [23] | |
| | Attention deficit hyperactivity disorder | ADHD | European (meta-analysis) | 20,183 cases and 35,191 controls | $5^{\ddagger}$ (0.01) | Demontis *et. al.*, 2019 [24] | `https://pgc.unc.edu/for-researchers/download-results/` |
| | Bipolar disorder | BD | European (meta-analysis) | 41,917 cases and 371,549 controls | $47^{\ddagger}$ (0.10) | Mullins *et. al.*, 2021 [25] | |
| | Autism spectrum disorder | ASD | European (meta-analysis) | 18,382 cases and 27,969 controls | $1^{\ddagger}$ (¿0.01) | Grove *et. al.*, 2019 [26] | |
| | Schizophrenia | SCZ | Trans-ethnic (meta-analysis) | | $233^{\ddagger}$ (0.50) | Trubetskoy *et. al.*, 2022 [27] | |
| | Cognition | CO | European (meta-analysis) | 257,828 | $154^{\ddagger}$ (33%) | Lee *et. al.*, 2018 [17] | `https://thessgac.com/` |

**Table 1. Overview of summary-level statistics used in the real data application to detect the impact of lifestyle and behavioural traits on mental health phenotypes.** $^{\dagger}$Number of non-unique genetic variants selected as IVs for each expsosure. $^{\ddagger}$Number of non-unique genetic variants selected as IVs for each phenotypic trait in the reverse causal analysis.

| Outcome | Exposure | MACE | mPPI | Empirical $p$-value | FDR |
|---|---|---|---|---|---|
| MDD | EDU | -0.005 | 0.136 | 0.693 | 0.693 |
| | **PA** | -0.168 | 0.971 | 0.001 | 0.002 |
| | SP | -0.023 | 0.291 | 0.251 | 0.301 |
| | ALC | -0.078 | 0.482 | 0.208 | 0.301 |
| | **SM** | 0.608 | 1.000 | $1.00\times10^{-7}$ | $1.00\times10^{-7}$ |
| | LST | 0.023 | 0.362 | 0.054 | 0.107 |
| AN | EDU | 0.049 | 0.286 | 0.115 | 0.173 |
| | PA | 0.085 | 0.388 | 0.087 | 0.173 |
| | **SP** | -0.374 | 0.943 | 0.003 | 0.009 |
| | ALC | -0.047 | 0.249 | 0.945 | 0.945 |
| | SM | 0.094 | 0.347 | 0.228 | 0.273 |
| | **LST** | -0.397 | 1.000 | $1.00\times10^{-7}$ | $6.00\times10^{-5}$ |
| ADHD | **EDU** | -0.750 | 1.000 | $1.00\times10^{-7}$ | $3.00\times10^{-5}$ |
| | PA | 0.029 | 0.203 | 0.377 | 0.565 |
| | SP | -0.010 | 0.135 | 0.965 | 0.965 |
| | ALC | -0.037 | 0.236 | 0.965 | 0.965 |
| | **SM** | 1.464 | 1.000 | $1.00\times10^{-7}$ | $3.00\times10^{-5}$ |
| | **LST** | 0.273 | 0.983 | $1.00\times10^{-7}$ | $3.00\times10^{-5}$ |
| ASD | **EDU** | 0.723 | 1.000 | $1.00\times10^{-7}$ | $6.00\times10^{-5}$ |
| | **PA** | -0.251 | 0.769 | 0.009 | 0.014 |
| | SP | 0.002 | 0.114 | 1.000 | 1.000 |
| | **ALC** | -0.608 | 0.909 | 0.010 | 0.014 |
| | **SM** | 0.480 | 0.908 | 0.004 | 0.011 |
| | **LST** | 0.073 | 0.423 | 0.037 | 0.044 |
| BD | **EDU** | 0.396 | 0.994 | $9.00\times10^{-5}$ | 0.000 |
| | PA | 0.107 | 0.476 | 0.053 | 0.079 |
| | SP | 0.099 | 0.462 | 0.092 | 0.110 |
| | ALC | 0.057 | 0.282 | 0.864 | 0.864 |
| | **SM** | 1.115 | 1.000 | $1.00\times10^{-7}$ | $6.00\times10^{-5}$ |
| | **LST** | -0.133 | 0.712 | 0.008 | 0.016 |
| SCZ | EDU | 0.041 | 0.266 | 0.134 | 0.201 |
| | PA | 0.005 | 0.121 | 0.974 | 0.974 |
| | **SP** | 0.423 | 0.986 | 0.001 | 0.001 |
| | ALC | 0.063 | 0.282 | 0.863 | 0.974 |
| | **SM** | 1.233 | 1.000 | $1.00\times10^{-7}$ | $6.00\times10^{-5}$ |
| | **LST** | -0.329 | 0.998 | $3.00\times10^{-5}$ | $9.00\times10^{-5}$ |
| COG | **EDU** | 0.820 | 1.000 | $1.00\times10^{-7}$ | $6.00\times10^{-5}$ |
| | **PA** | -0.062 | 0.724 | 0.013 | 0.031 |
| | SP | -0.018 | 0.324 | 0.198 | 0.297 |
| | ALC | 0.016 | 0.270 | 0.898 | 0.898 |
| | **SM** | 0.094 | 0.785 | 0.016 | 0.031 |
| | LST | 0.002 | 0.106 | 0.751 | 0.898 |

**Table 2. Results of Bayesian model averaging (MR-BMA) algorithm regarding how life-style and behavioural exposures impact mental health outcomes.** MR-BMA [8] is run on each outcome separately. Results include the model-averaged causal effect estimate (MACE), the marginal posterior probability of inclusion (mPPI) and the corresponding empirical $p$-value [28]. We adjust for multiple testing using Benjamini-Hochberg False Discovery Rate (FDR). Exposures selected at 5% FDR are highlighted in bold.

### 3 Supplementary Figures

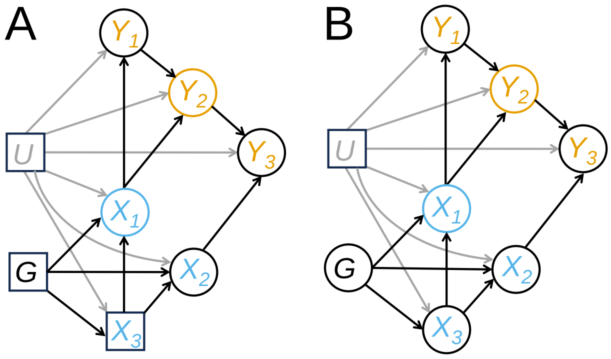

**Figure 1. Sufficient adjustment sets required by the back-door criterion** to guarantee that the association between $X_1$ and $Y_2$ is purely causative in the example depicted in the main text Figure 1C. (**A**) $Y_2(x_1) \perp\!\!\!\perp X_1 \mid C, \forall x_1$, where $C = \{X_3, \boldsymbol{G}, U\}$ is the sufficient adjustment set [29]. By conditioning on $\{X_3, \boldsymbol{G}, U\}$, depicted as squares, there are no back-door paths from $X_1$ to $Y_2$. (**B**) $Y_2(x_1) \perp\!\!\!\perp X_1 \mid C, \forall x_1$, where $C = \{U\}$. The second sufficient adjustment set is obtained by using the function `adjustmentSets()` in the R package *dagitty* which implements the adjustment criterion presented in [30], an extension of Pearl's back-door criterion [4]. In (A) and (B), both sufficient adjustment sets include the unmeasured confounder $U$.

#### Simulation study

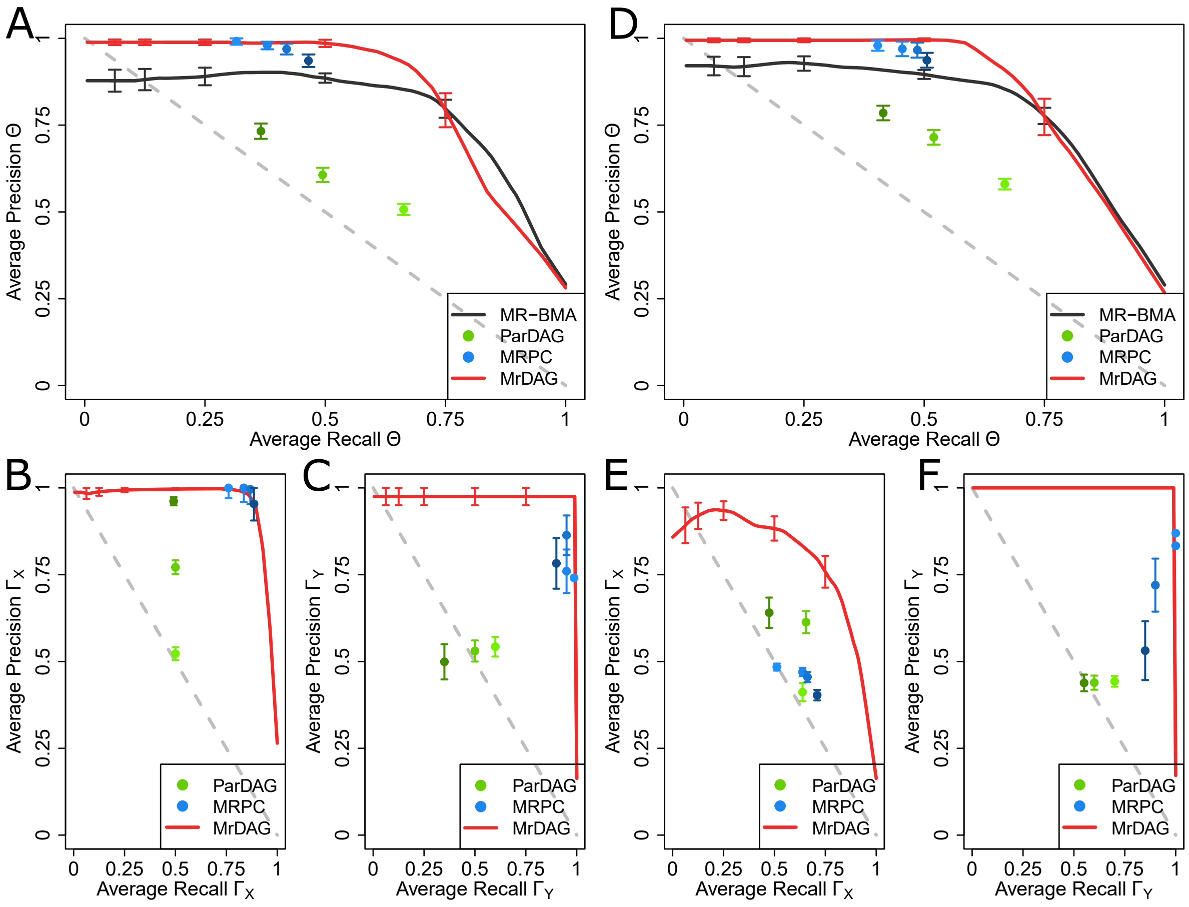

**Figure 2. Precision-Recall Curves (PRCs) for all methods considered in the simulated scenarios UndG$_X$-Med$_Y$ and DAG$_X$-Med$_Y$** show recall (= sensitivity = TP/(TP+FN)) in the $x$-axis and precision (= positive predictive value = TP/(TP+FP)) in the $y$-axis with TP = True Positive, FN = False Negative and FN = False Positive averaged over 25 replicates in each scenario and for all methods considered. In scenario UndG$_X$-Med$_Y$ (**A-C**), the strength of the correlation between consecutive $\boldsymbol{X}$ is set at $r_X = 0.6$, and then it decreases exponentially for non-consecutive exposures, and the level of complete mediation in $\boldsymbol{Y}$ is set at $m_Y = 1$, while in scenario DAG$_X$-Med$_Y$ (**D-F**), the average level of the mediation parameters within $\boldsymbol{X}$ and the level of complete mediation in $\boldsymbol{Y}$ is set at $r_X = 0.6$ and $m_Y = 1$, respectively. For details, see in the main text Methods. In both scenarios, the results are presented separately for the simulated dependency structures from the exposures to the outcomes (A and D), within the exposures (B and E) and the outcomes (C and D), respectively. Vertical bars in each PRCs, at specific recall levels $0.0625, 0.125, 0.25, 0.50$ and $0.75$, indicate standard error. Vertical bars in each PRC, at specific recall levels $0.0625, 0.125, 0.25, 0.50$ and $0.75$, indicate standard error. For the MRPC algorithm, type I error rate for the conditional independence test is set at $\alpha = \{0.01, 0.05, 0.10, 0.20\}$ (from light- to dark-blue dots) and for the ParDAG algorithm we specify three different values for the Lasso penalisation $\lambda = \{0.5, 0.7, 0.9\}$ (from light- to dark-green dots). See Section 'Simulation study and real data application' for details.

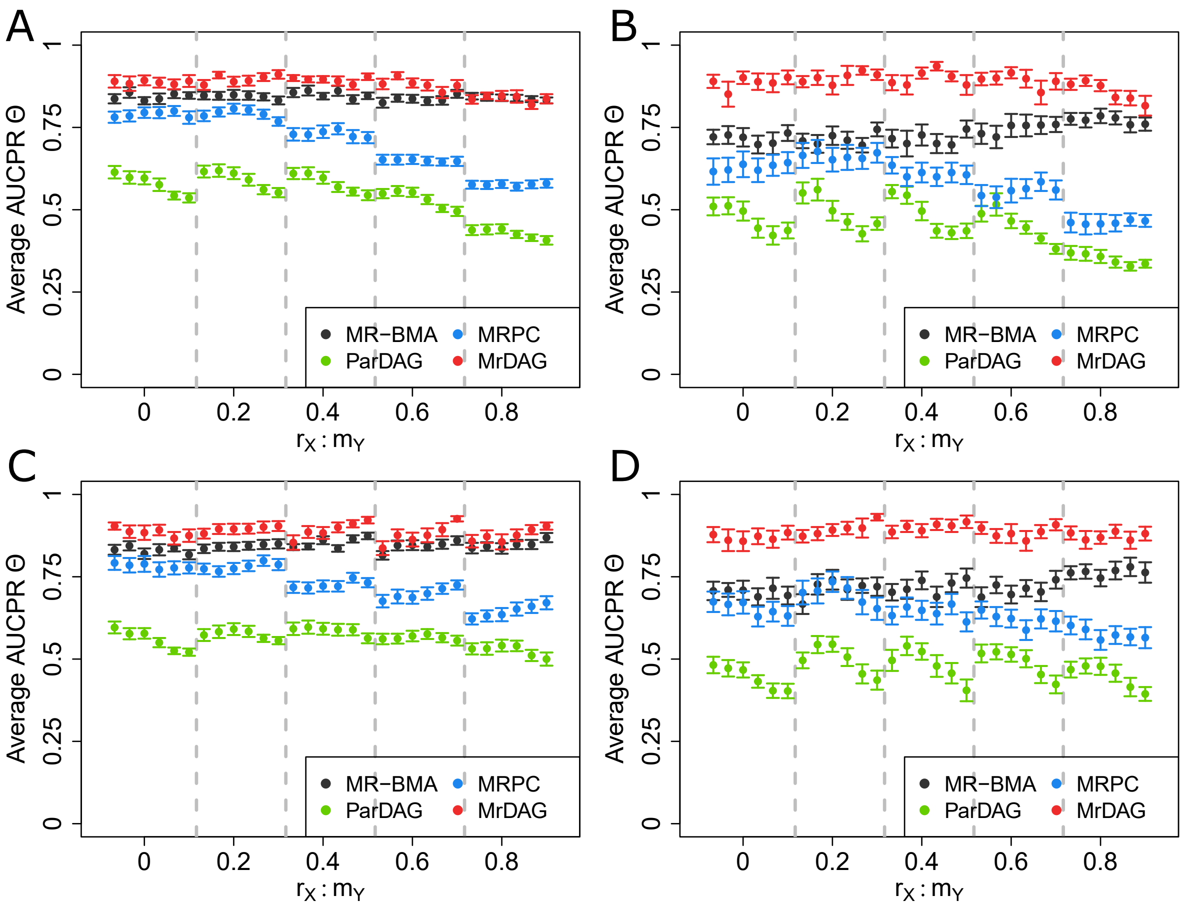

**Figure 3. Area Under the Curve of Precision-Recall (AUCPR) of the causal effects Θ between the exposures and the outcomes for all methods considered and simulated scenarios** averaged across 25 replicates for different value of the parameters $r_X = \{0, 0.2, 0.4, 0.6, 0.8\}$ which controls the strength of correlation between consecutive $\boldsymbol{X}$, and then it decreases exponentially for non-consecutive exposures, in scenarios (**A**) UndG$_X$-Med$_Y$ and (**B**) UndG$_X$-DAG$_Y$ or the average level of the mediation parameters within $\boldsymbol{X}$ in scenarios (**C**) DAG$_X$-Med$_Y$ and (**D**) DAG$_X$-DAG$_Y$, and $m_Y = \{0.25, 0.50, 0.75, 1, 1.5, 2\}$ which regulates the level of complete mediation in scenarios (**A**) UndG$_X$-Med$_Y$ and (**C**) DAG$_X$-Med$_Y$ or the average level of the mediation parameters within $\boldsymbol{Y}$ in scenarios (**B**) UndG$_X$-DAG$_Y$ and (**D**) DAG$_X$-DAG$_Y$, respectively. For details, see in the main text Methods. At each value of $r_X$ in the $x$-axis and delimited by vertical-dotted lines, the AUCPRs are plotted against all values of $m_Y$. The average value of the AUCPR and its standard error (vertical bar for each AUCPR) has been calculated using the functions `prediction()` and `performance()` in the R package *ROCR* [31, 32]. For MRPC and ParDAG algorithms, we only show the results obtained at type I error rate for the conditional independence test $\alpha = 0.01$ and Lasso penalisation $\lambda = 0.9$, respectively. These values provide the best results for the two algorithms as shown in the main text Figure 4 and Supplementary Figure 2.

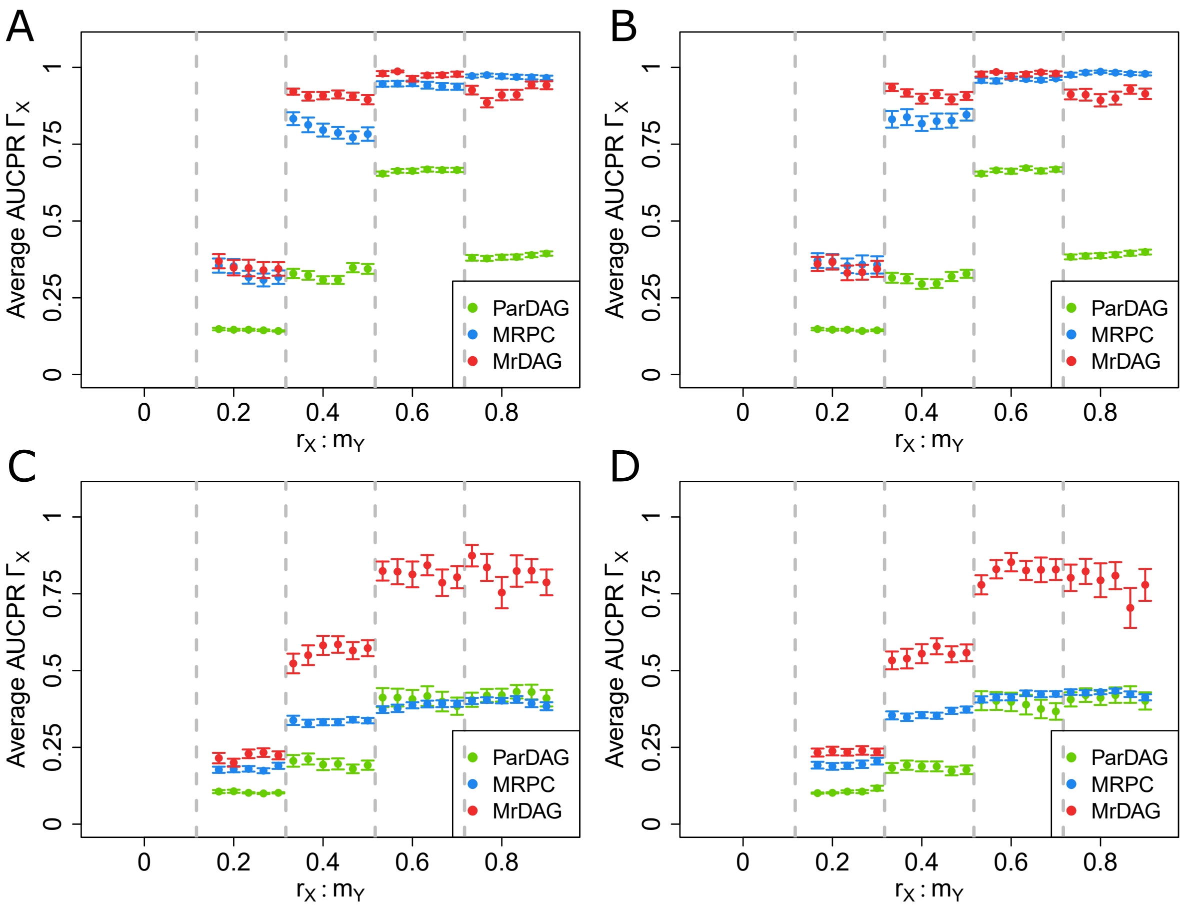

**Figure 4. Area Under the Curve of Precision-Recall (AUCPR) of the detection of the dependency structure within the exposures for all methods considered and simulated scenarios** averaged across 25 replicates for different value of the parameters $r_X = \{0, 0.2, 0.4, 0.6, 0.8\}$ which controls the strength of correlation between consecutive $\boldsymbol{X}$, and then it decreases exponentially for non-consecutive exposures, in scenarios (**A**) UndG$_X$-Med$_Y$ and (**B**) UndG$_X$-DAG$_Y$ or the average level of the mediation parameters within $\boldsymbol{X}$ in scenarios (**C**) DAG$_X$-Med$_Y$ and (**D**) DAG$_X$-DAG$_Y$, and $m_Y = \{0.25, 0.50, 0.75, 1, 1.5, 2\}$ which regulates the average level of complete mediation in scenarios (**A**) UndG$_X$-Med$_Y$ and (**C**) DAG$_X$-Med$_Y$ or the average level of the mediation parameters within $\boldsymbol{Y}$ in scenarios (**B**) UndG$_X$-DAG$_Y$ and (**D**) DAG$_X$-DAG$_Y$, respectively. For details, see in the main text Methods. At each value of $r_X$ in the $x$-axis and delimited by vertical-dotted lines, the AUCPRs are plotted against all values of $m_Y$. The average value of the AUCPR and its standard error (vertical bar for each AUCPR) has been calculated using the functions `prediction()` and `performance()` in the R package *ROCR* [31, 32]. For MRPC and ParDAG algorithms, we only show the results obtained at type I error rate for the conditional independence test $\alpha = 0.01$ and Lasso penalisation $\lambda = 0.9$, respectively. These values provide the best results for the two algorithms as shown in the main text Figure 4 and Supplementary Figure 2.

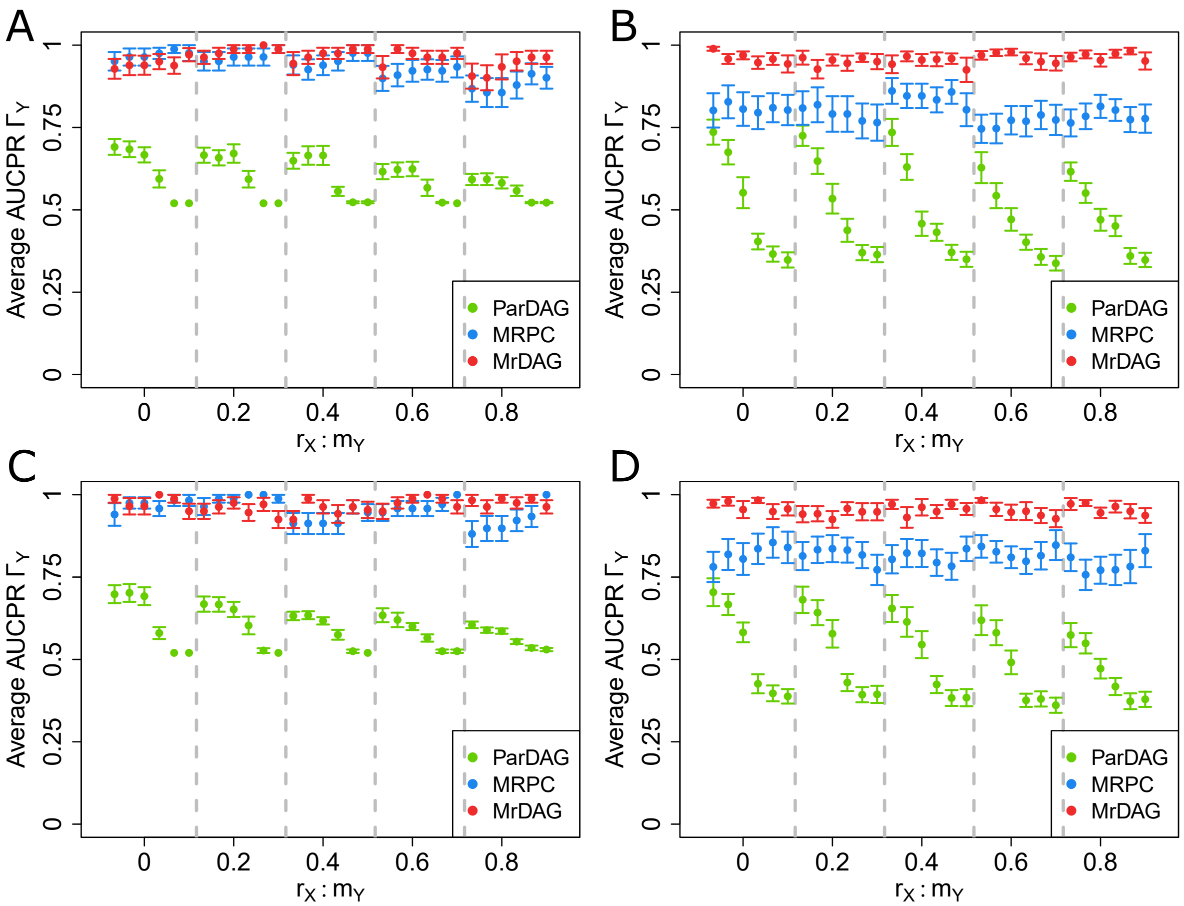

**Figure 5. Area Under the Curve of Precision-Recall (AUCPR) of the detection of the dependency structure within the outcomes for all methods considered and simulated scenarios** averaged across 25 replicates for different value of the parameters $r_X = \{0, 0.2, 0.4, 0.6, 0.8\}$ which controls the strength of correlation between consecutive $\boldsymbol{X}$, and then it decreases exponentially for non-consecutive exposures, in scenarios (**A**) UndG$_X$-Med$_Y$ and (**B**) UndG$_X$-DAG$_Y$ or the average level of the mediation parameters within $\boldsymbol{X}$ in scenarios (**C**) DAG$_X$-Med$_Y$ and (**D**) DAG$_X$-DAG$_Y$, and $m_Y = \{0.25, 0.50, 0.75, 1, 1.5, 2\}$ which regulates the level of complete mediation in scenarios (**A**) UndG$_X$-Med$_Y$ and (**C**) DAG$_X$-Med$_Y$ or the average level of the mediation parameters within $\boldsymbol{Y}$ in scenarios (**B**) UndG$_X$-DAG$_Y$ and (**D**) DAG$_X$-DAG$_Y$, respectively. For details, see in the main text Methods. At each value of $r_X$ in the $x$-axis and delimited by vertical-dotted lines, the AUCPRs are plotted against all values of $m_Y$. The average value of the AUCPR and its standard error (vertical bar for each AUCPR) has been calculated using the functions `prediction()` and `performance()` in the R package *ROCR* [31, 32]. For MRPC and ParDAG algorithms, we only show the results obtained at type I error rate for the conditional independence test $\alpha = 0.01$ and Lasso penalisation $\lambda = 0.9$, respectively. These values provide the best results for the two algorithms as shown in the main text Figure 4 and Supplementary Figure 2.

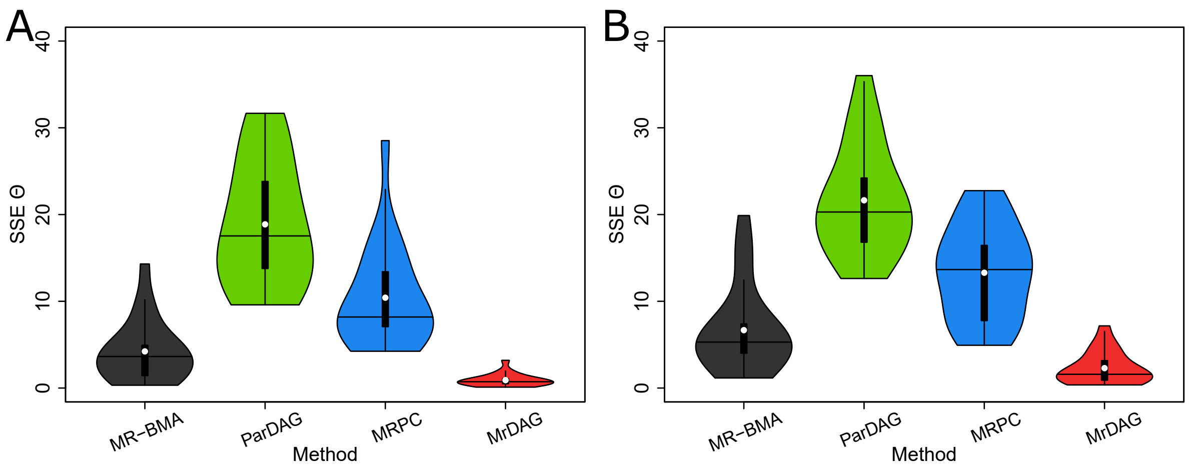

**Figure 6. Violin plots of the Sum of Squares Error (SSE) of the causal effects $\mathbf{\Theta}$ between the exposures and the outcomes for all methods considered in the simulated scenarios UndG$_X$-Med$_Y$ and DAG$_X$-Med$_Y$** across 25 replicates in each scenario. (**A**) In scenario UndG$_X$-Med$_Y$, the strength of correlation between consecutive $\boldsymbol{X}$ is set at $r_X = 0.6$, and then it decreases exponentially for non-consecutive exposures, and the level of complete mediation in $\boldsymbol{Y}$ is set at $m_Y = 1$. (**B**) In scenario DAG$_X$-Med$_Y$, the average level of the mediation parameters within $\boldsymbol{X}$ and the level of complete mediation in $\boldsymbol{Y}$ is set at $r_X = 0.6$ and $m_Y = 1$, respectively. For details, see in the main text Methods. In each violin plot, the vertical black thick line displays the interquartile range, the black horizontal line denotes the median and the white dot the mean. For MRPC and ParDAG algorithms, we only show the results obtained at type I error rate for the conditional independence test $\alpha = 0.01$ and Lasso penalisation $\lambda = 0.9$, respectively. These values provide the best results for the two algorithms as shown in the main text Figure 4 and Supplementary Figure 2.

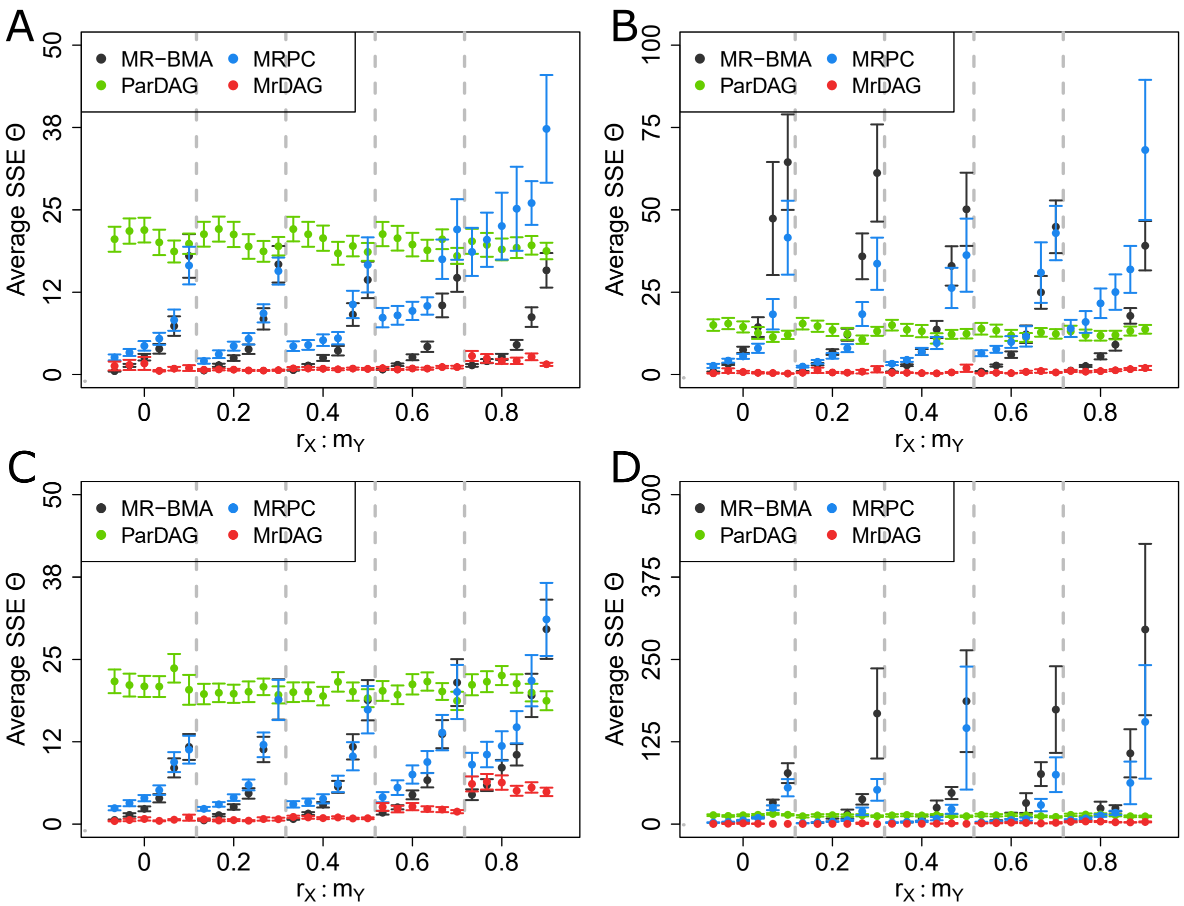

**Figure 7. Sum of Squared Error (SSE) of the causal effects Θ between the exposures and the outcomes for all methods considered and simulated scenarios** averaged across 25 replicates for different value of the parameters $r_X = \{0, 0.2, 0.4, 0.6, 0.8\}$ which controls the strength of correlation between consecutive exposures, and then it decreases exponentially for non-consecutive exposures, in scenarios (**A**) UndG$_X$-Med$_Y$ and (**B**) UndG$_X$-DAG$_Y$ or the average level of the mediation parameters within $\boldsymbol{X}$ in scenarios (**C**) DAG$_X$-Med$_Y$ and (**D**) DAG$_X$-DAG$_Y$, and $m_Y = \{0.25, 0.50, 0.75, 1, 1.5, 2\}$ which regulates the level of complete mediation in scenarios (**A**) UndG$_X$-Med$_Y$ and (**C**) DAG$_X$-Med$_Y$ or the average level of the mediation parameters within $\boldsymbol{Y}$ in scenarios (**B**) UndG$_X$-DAG$_Y$ and (**D**) DAG$_X$-DAG$_Y$, respectively. For details, see in the main text Methods. At each value of $r_X$ in the $x$-axis and delimited by vertical-dotted lines, the SSEs are plotted against all values of $m_Y$. Vertical bars for each SSE indicate standard error. For MRPC and ParDAG algorithms, we only show the results obtained at type I error rate for the conditional independence test $\alpha = 0.01$ and Lasso penalisation $\lambda = 0.9$, respectively. These values provide the best results for the two algorithms as shown in the main text Figure 4 and Supplementary Figure 2

.

#### Real data application

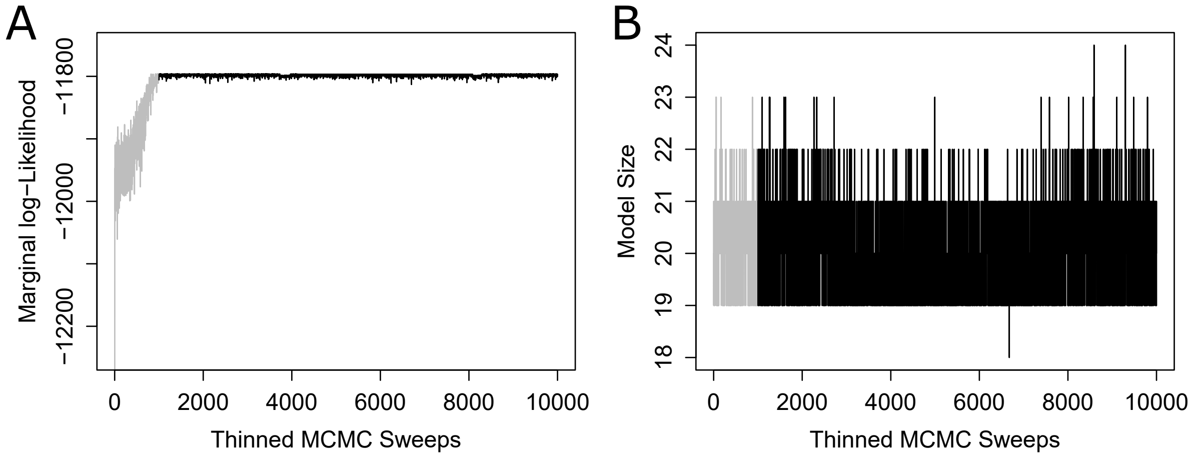

**Figure 8. Convergence diagnostics of MrDAG algorithm to assess how lifestyle and behavioural exposures impact mental health outcomes.** Grey colour denotes burn-in up to $10^5$ MCMC sweeps, while black colour shows values after the burn-in. Quantities reported in the $y$-axis are recorded at every 100 MCMC sweeps, resulting in $10,000$ thinned recorded values ($x$-axis). (**A**) Trace plot of the log-marginal likelihood $m_{\mathcal{G}}(\text{data})$ during $10^6$ MCMC sweeps. In the burn-in, the effect of the annealing parameter produces a larger excursion of the log-marginal likelihood, allowing the algorithm to escape from local maxima and thus favouring an efficient exploration of regions of high posterior mass. The acceptance rate of the proposed DAGs (which belongs to the Markov Equivalent Class) under partial ordering from the exposures to the outcomes is $\approx 90\%$ and thus it does not influence the convergence of the algorithm and structure learning. (**B**) Trace plot of the posterior model size, *i.e.*, the number of directed edges selected during MCMC sweeps. The range of the model size is between 18 and 24 with an average of $\approx 21$ selected edges. After burn-in, the posterior model size is much larger than the prior model size which is set at one edge for each of the $(q + p)$ nodes with $q = 7$ for the outcomes and $p = 6$ for the exposures, resulting in 13 nodes.

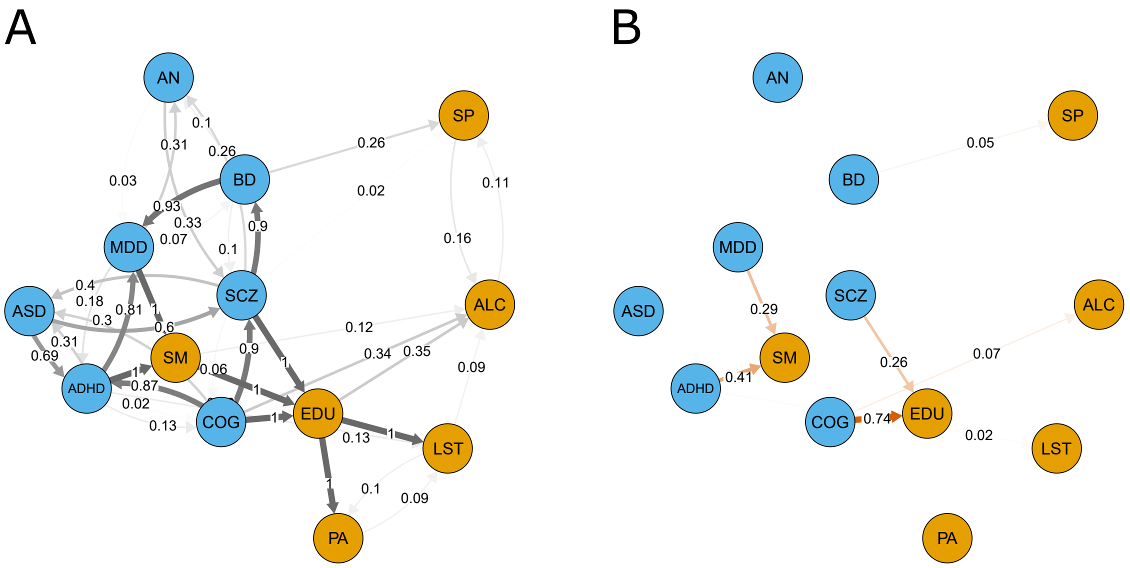

**Figure 9. Partially DAGs (PDAGs) representation of MrDAG results regarding how liability to mental health phenotypes affects lifestyle and behavioural traits.** (**A**) PDAG of the posterior probability of edge inclusion (PPeI) (12) within the exposures (mental health phenotypes, blue nodes), the outcomes (lifestyle and behavioural traits, orange nodes) and between them. Undirected edges are represented as bidirectional edges, see, for instance, edges between PA (physical activity) and LST (leisure screen time) or ASD (autism spectrum disorder) and ADHD (attention deficit hyperactivity disorder). Neither reverse causation from the outcomes to the exposures nor feedback loops are allowed. (**B**) Posterior causal effects on the outcomes (orange nodes) under intervention on the exposures (blue nodes). Red and green edges indicate positive and negative posterior causal effects, respectively (14).

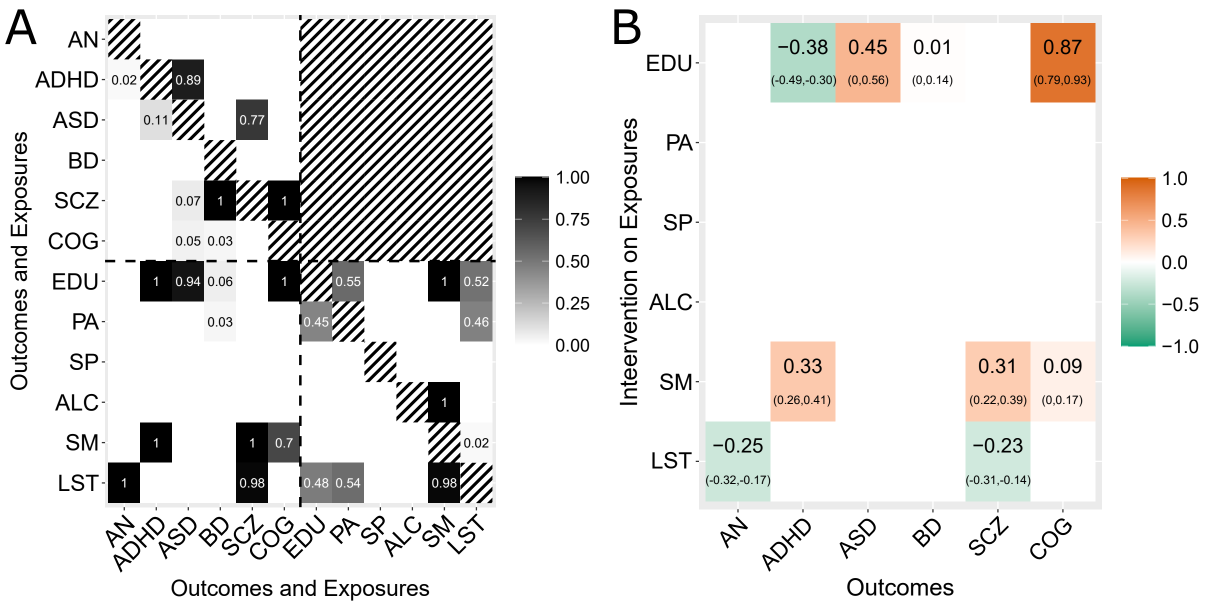

**Figure 10. Results of MrDAG regarding how lifestyle and behavioural exposures impact mental health outcomes when major depression disorder (MDD) is removed from the list of outcomes.** (**A**) Posterior probability of edge inclusion (PPeI) for each combination of outcomes (mental health phenotypes) and exposures (lifestyle and behavioural traits) when MDD is removed from the list of outcomes. Horizontal and vertical dotted lines separate the exposures (bottom-right submatrix) from the outcomes (top-left submatrix). PPEIs between exposures and outcomes are depicted in the bottom-left submatrix. Neither reverse causation (top-right submatrix) nor feedback loops (main diagonal) are allowed (black-white strips). (**B**) Posterior causal effects (95% credible intervals) on the outcomes ($y$-axis) under intervention on the exposures ($x$-axis) when MDD is excluded from the list of outcomes. The results are obtained by specifying the same prior probability of edge inclusion $\pi^{\text{edge}} = 0.16$ used in the main text Section 'Real data application: The impact of lifestyle and behavioural traits on mental health'.

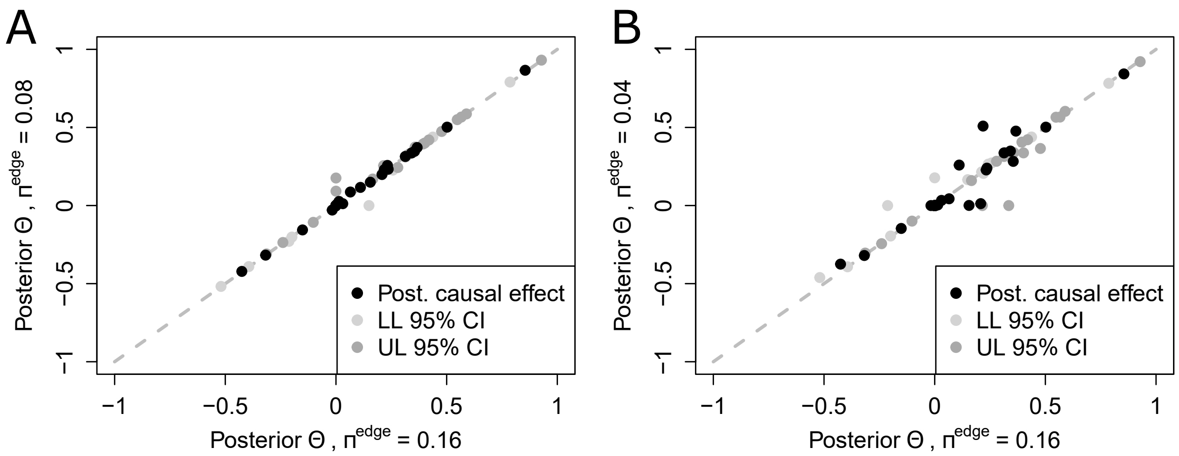

**Figure 11. Sensitivity analysis of MrDAG results regarding how lifestyle and behavioural exposures impact mental health outcomes for different values of the *a priori* probability of edge inclusion.** Posterior causal effects (black dots) and 95% credible intervals (Lower Limit - light-grey dots, Upper Limit - dark-grey dots) obtained by MrDAG algorithm for different levels of sparsity controlled by the hyper-parameter $\pi^{\text{edge}}$. In each panel, the $x$-axis reports the posterior causal effects at $\pi^{\text{edge}} = 0.16$ used in the main text Section 'Real data application: The impact of lifestyle and behavioural traits on mental health' and in the $y$-axis the posterior causal effects at (**A**) $\pi^{\text{edge}} = 0.08$ and (**B**) $\pi^{\text{edge}} = 0.04$. Overall, MrDAG algorithm is robust with only a handful of cases (panel B) where the posterior causal effects and the credible intervals slightly do not agree at different levels of $\pi^{\text{edge}}$.

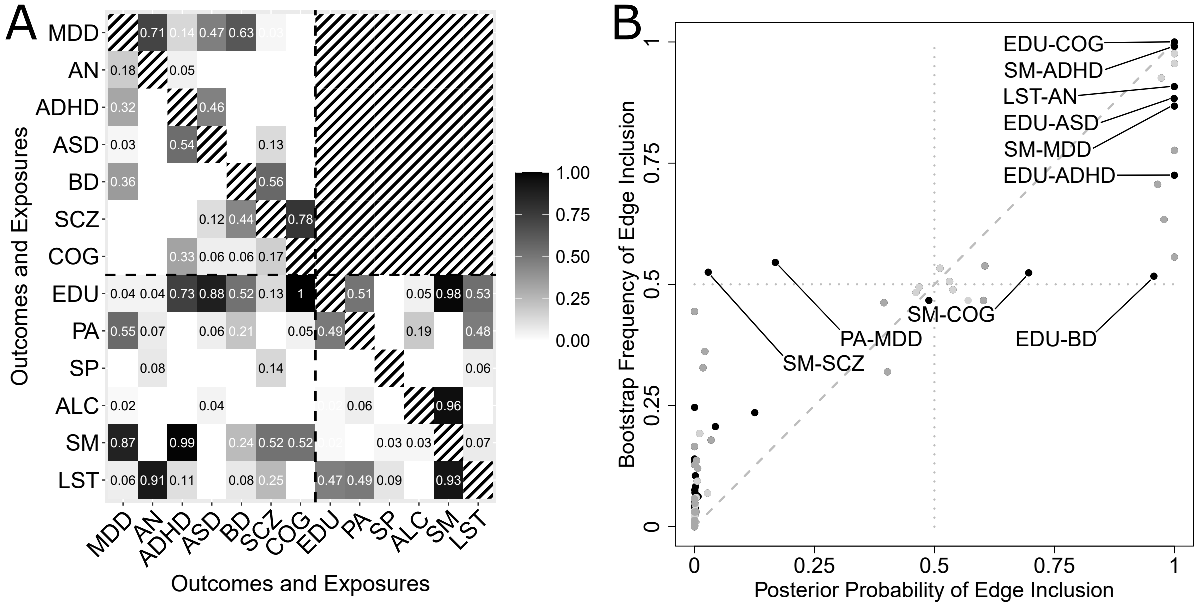

**Figure 12. Robustness of MrDAG results regarding how lifestyle and behavioural exposures impact mental health outcomes.** (**A**) Bootstrap frequency of edge inclusion is calculated as the average of the posterior probability of edge inclusion (PPeI) obtained by running MrDAG algorithm on each bootstrap summary-level statistics. Horizontal and vertical dotted lines separate the exposures (bottom-right submatrix) from the outcomes (top-left submatrix). Bootstrap frequencies between exposures and outcomes are depicted in the bottom-left submatrix. (**B**) Scatterplot of PPeI against the bootstrap frequency of edge inclusion. For values of the bootstrap frequency of edge inclusion ($y$-axis) $\geq 0.5$ (horizontal dotted line), exposure-outcome pairs (black dots) are highlighted.
